# Transient Water Wires Mediate Selective Proton Transport in Designed Channel Proteins

**DOI:** 10.1101/2022.03.28.485852

**Authors:** Huong T. Kratochvil, Laura C. Watkins, Marco Mravic, Noah H. Somberg, Jessica L. Thomaston, John M. Nicoludis, Lijun Liu, Mei Hong, Gregory A. Voth, William F. DeGrado

## Abstract

Selective proton transport through proteins is essential for forming and utilizing proton gradients in cells. Protons are conducted along hydrogen-bonded “wires” of water molecules and polar sidechains, which, somewhat surprisingly, are often interrupted by dry apolar stretches in the conduction pathways inferred from static protein structures. We hypothesize that protons are conducted through such dry spots by forming transient water wires, often highly correlated with the presence of the excess proton itself in the water wire. To test this hypothesis, we used molecular dynamics simulations to design transmembrane channels with stable water pockets interspersed by apolar segments capable of forming flickering water wires. The minimalist designed channels conduct protons at rates similar to viral proton channels, and they are at least 10^6^-fold more selective for H^+^ over Na^+^. These studies inform mechanisms of biological proton conduction and principles for engineering proton-conductive materials.

---

The controlled diffusion of protons through transmembrane proteins is critical for many aspects of physiological function, including substrate transport ^1^, control of cellular and organelle pH ^2^, the creation and utilization of pH gradients required for bioenergetics ^3,4^, and cellular signaling ^5^. Proteins conduct protons along precisely defined pathways that prevent wasteful collapse of Na^+^ and K^+^ gradients. Because the proton concentration in the cellular cytoplasm is about 10^6^-fold lower than that of other ions at neutral pH, proton channels must have selectivities significantly greater than this value. The conduction of protons in water and aqueous pores is facilitated by the formation of water wires consisting of hydrogen-bonded chains of waters. In the classical Grotthuss mechanism, protons pass from one water molecule to the next to achieve long-range net transport without the need to move oxygen atoms (Fig. 1a) ^6^. In this mechanism, the excess proton first forms a hydrated structure with a hydronium-like core ^7^, creating a net positive charge defect or hole that propagates “down-stream” of its initial position along the water wire ^8^.

**Fig. 1.**
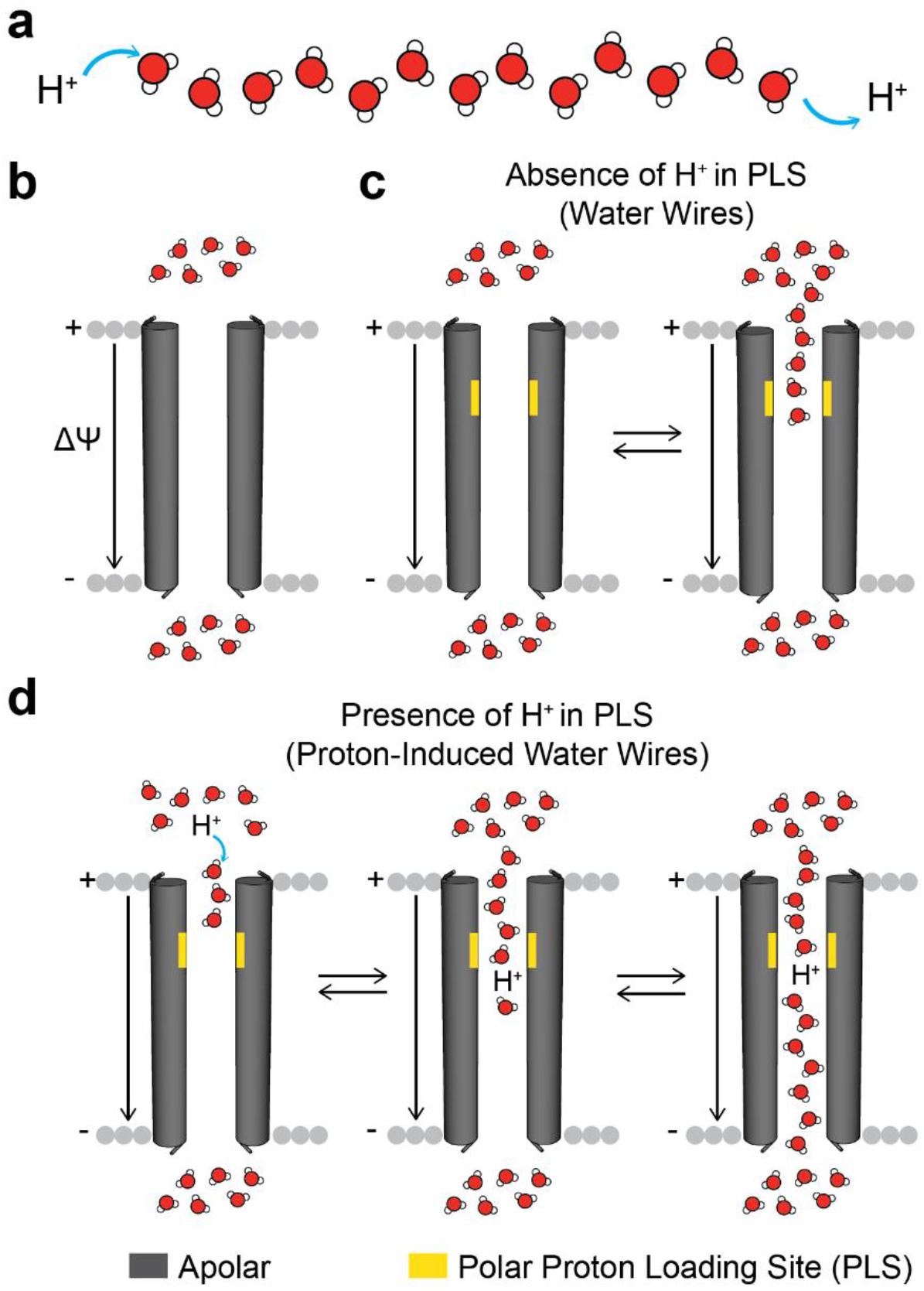
Hypothesis of proton-selective transport along transient water wires. **a**, Protons hop across dynamically rearranging hydrogen-bonded wires of water in the Grotthuss mechanism. **b**, In a wholly apolar channel, the pore remains devoid of water regardless of membrane polarization (ΔΨ), but a polar PLS, **c**, can mediate the flickering of neutral water in and out of a relatively short apolar sectors of the channel. **d**, The presence of a hydrated excess proton facilitates water wire formation and transport of an excess proton through the hydrophobic sector by Grotthuss shuttling.

Proteins achieve high proton selectivity by organizing and gating such water wires in proton conduction pathways often interspersed with ionizable sidechains, which explicitly participate in Grotthuss shuttling ^9,10^.Somewhat surprisingly, however, protons often appear to be conducted through dry stretches of hydrophobic residues that feature in X-ray and cryo-electron microscopic structures of proteins ^8,11–13^. Classical molecular dynamics (MD), reactive molecular dynamics (RMD), and quantum mechanical calculations suggest that water can occasionally penetrate through such apolar sectors with the help of polar residues, forming transient water wires not apparent in the time-averaged structures (Fig. 1c) ^8,11^. Thus, rare equilibrium fluctuations mediated by polar proton loading sites (PLSs) provide one mechanism for transient protonated water wire formation. Additionally, in systems that are energetically activated by chemical reactions, light or a transmembrane potential, the arrival of an excess proton can induce the formation of transient water wires through confined, hydrophobic spaces (Fig. 1d) ^8,11,14^. One would expect very high proton selectivity from such a mechanism because a single column of connected water wires in a hydrophobic environment is unable to accommodate or stabilize a hydrated Na^+^ or K^+^ ion ^15–20^. However, experimental evidence for such mechanisms has been indirect or lacking, with the exception of the very extensively studied protein bacteriorhodopsin where transient spectroscopy and serial crystallography have identified water molecules that form during portions of its photo-cycle ^13,21–23^. However, this system represents only a single example in which large changes are induced by photo-isomerization.

We therefore turned to de novo design ^24–28^ to test and expand the transient water wire hypothesis. The design of a highly proton-selective channel operating by this mechanism not only tests an important concept in biophysics, but it also represents an important challenge in *de novo* protein design. While it has been possible to design stably-packed proteins that bind small molecules and protein interfaces ^29–31^ more complex functions that rely on chemical dynamics have been difficult to design from scratch. Except for early studies where transition states were considered as targets in the design of catalytic proteins ^32,33^, *de novo* design has generally focused on ground states rather than non-equilibrium high-energy states, as are generated during ion conduction. Computational design algorithms also favor tight and efficient packing are likely to dampen essential fluctuations required for catalysis and transport ^34–36^. Moreover, *de novo* protein design does not consider explicit water molecules, and instead relies on approximations of the effect of solvent. Here, we not only consider water explicitly, but we also account for the dynamic formation and breaking of covalent bonds as protons are passed from one water to the next. Finally, despite a few successes ^27,37–40^, the design and high-resolution structural characterization of membrane proteins remains a difficult endeavor. Two *de novo* channel-forming proteins have been structurally characterized, but they were not highly selective, nor were their per-channel conductance rates determined^38^. Other work focused on the conversion of water-soluble nanopores into membrane-spanning channels yielded channels with well-defined single-channel conductance characteristics, but the structure of the ion-conducting form of the channel was not determined^37^. Finally, tetrameric TM bundles were designed to use transition metal ion-binding to drive proton translocation and vice versa, but the anti-porting efficiency was limited by leakiness to protons ^39^. Thus, the purposeful design of highly proton-selective channels that operate by a dynamic wetting/dewetting mechanism represents a significant advance.

## MD-guided design of proton channels

To test the transient water wire hypothesis, we designed a series of channels containing a polar PLS adjacent to a hydrophobic pore (Fig. 2). The expected length (*l_exp_*) of the longest uninterrupted hydrophobic stretch along the pore axis, and the number and position of protein loading sites were varied. We chose a cluster of neutral Gln residues as the PLS to help stabilize a proton in the pore without falling into any deep energy wells that might occur with an ionizable residue. Gln and Asn feature in the proton conduction pathways of the S31N mutant of the influenza A M2 and OTOP proton channels^41,42^ Due to its two methylene groups, Gln is also flexible so that it should be able to stabilize multiple polarizations of water wires that are created during a conduction cycle.

**Fig. 2.**
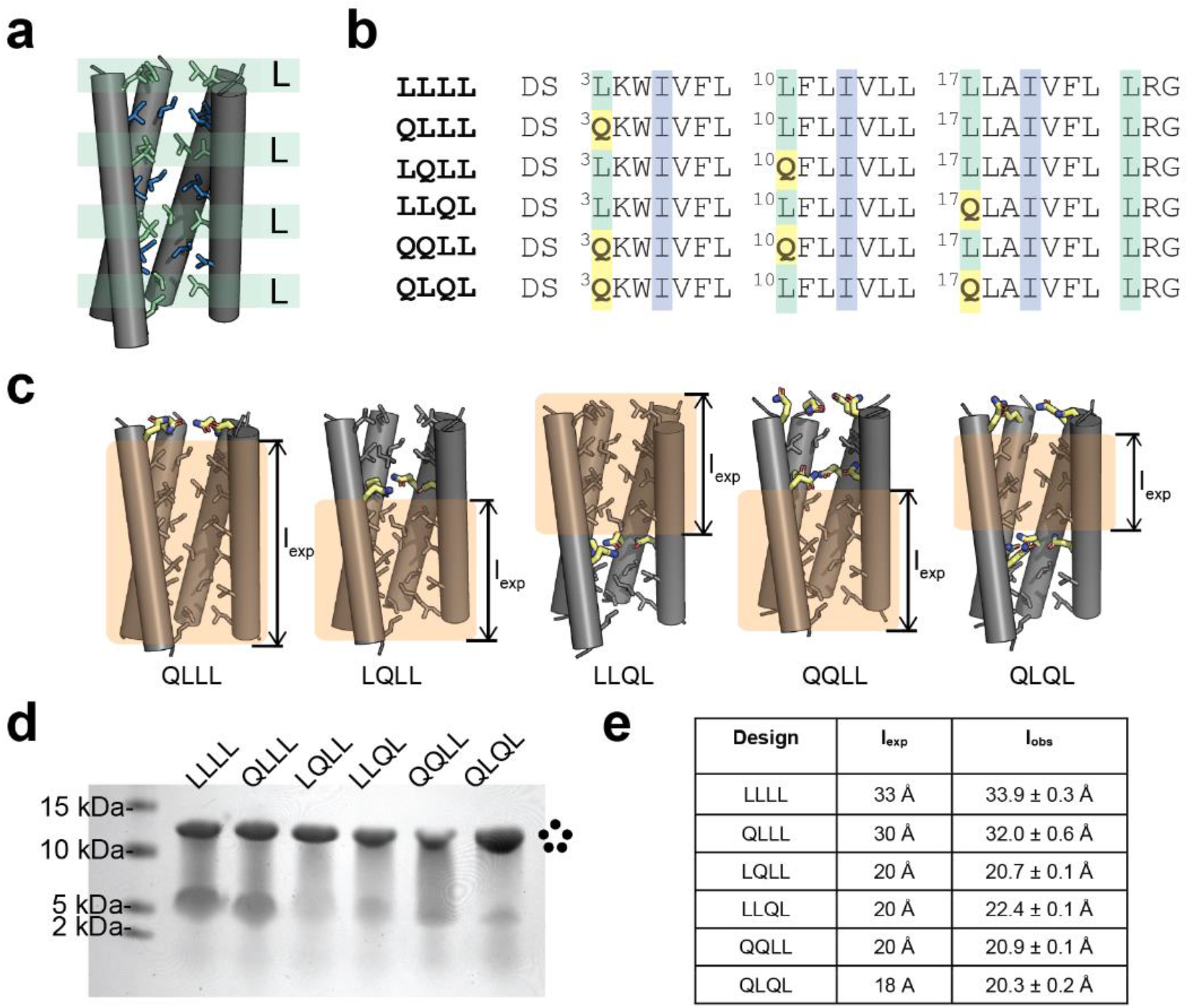
De novo channels incorporate PLSs at key positions to modulate hydrophobic lengths. **a**, The parent scaffold, LLLL (pdb 6mct), contains layers of interdigitating Leu (green) and Ile (blue) residues. **b**, Five Leu-to-Gln variants were designed. **c**, Models of the designed channels illustrates the different expected hydrophobic lengths (l_exp_) of the channels relative to the parent scaffold. **d**, SDS-PAGE revealed that these designs formed stable pentamers. **e**, Comparison of the longest hydrophobic length expected (l_exp_) and observed (l_obs_, determined by classical MD simulations; see Methods and Extended Data Table 1). Side views of models in Fig. 2, 3, 4 and 6 have the fifth helix removed for clarity.

Our constructs began with a previously characterized *de novo* homopentameric TM helical bundle with an interior stabilized by efficient van der Waals packing of alternating layers of apolar Leu and Ile residues (Fig. 2a, pdb 6mct) ^43^. A narrow, fully hydrophobic pore (app. 2 – 3 Å diameter), which is impervious to water in classical MD simulations, runs the entire length of the bundle ^43^. We introduced single and double Leu-to-Gln substitutions into the pore (Fig. 2b,d), thereby creating a site that filled with water in MD simulations. The Leu residues were targeted for substitution because they project directly towards the pore without steric clashes. The variants are designated as LLLL, QLLL, QQLL, etc., according to the identity of the Leu-to-Gln substitutions (Fig. 2). Seven variants with 0, 1, or 2 Gln substitutions were synthesized; six formed pentamers based on gel electrophoresis and were structurally and functionally characterized. By design, the expected length of the longest apolar path, *l_exp_*, was intended to be 33 Å for LLLL and 30 Å for QLLL and it was held constant at approximately 18 Å to 20 Å for the remaining five variants (Fig. 2c). Both single and double proton-loading sites were evaluated by varying the positions of the introduced Gln residues. A peptide with 3 Gln residues failed to form a pentameric bundle, so it was not possible to further decrease the value of *l_exp_*.

The hydration of each channel was evaluated in three independent 200 ns classical MD simulations starting with the computational design model in phospholipid bilayers (Fig. 3, Supplementary Figs. S1-S6). The pore of the starting LLLL (Fig. 3a) pentamer was devoid of significant water density throughout the trajectories. The length of the longest apolar path in its pore (*l_obs_* =33.9 ± 0.3 Å Fig. 3a) was in good agreement with design (*l_exp_* of 33 Å). QLLL (Fig. 3b) showed strong water density near its Gln sidechain near the entry to the pore, but the remainder of the pore was dry (*l_obs_* = 32.0 ± 0.6 Å). The single-site variant LQLL (Fig. 3c) has strong hydration near the Gln carboxamide sidechain, which communicates with the bulk water at the top of the channel via infrequent, flickering water-wires. The remaining C-terminal pore region is fully dehydrated over approximately 21 Å (*l_obs_* = 20.7 ± 0.1 Å). LLQL (Fig. 3d) shows inverse behavior with hydrated Gln residues, fluctuating water wires near the bottom of the channel, and a 22.4 ± 0.04 Å dehydrated pore at the top. The doubly-substituted pentamers had greater hydration associated with the additional interfacial Gln (Fig. 3e,f). However, by design, they still have three consecutive layers of pore-lining Leu and Ile residues, which results in having dry regions of similar length to the longest stretch seen in the single-Gln variants (*l_obs_* = 20.9 ± 0.04 Å and 20.3 ± 0.2 Å for QQLL and QLQL, respectively). Thus, by incorporating classical MD simulations into the design process, we can assess the potential of design candidates for experimental validation.

**Fig. 3.**
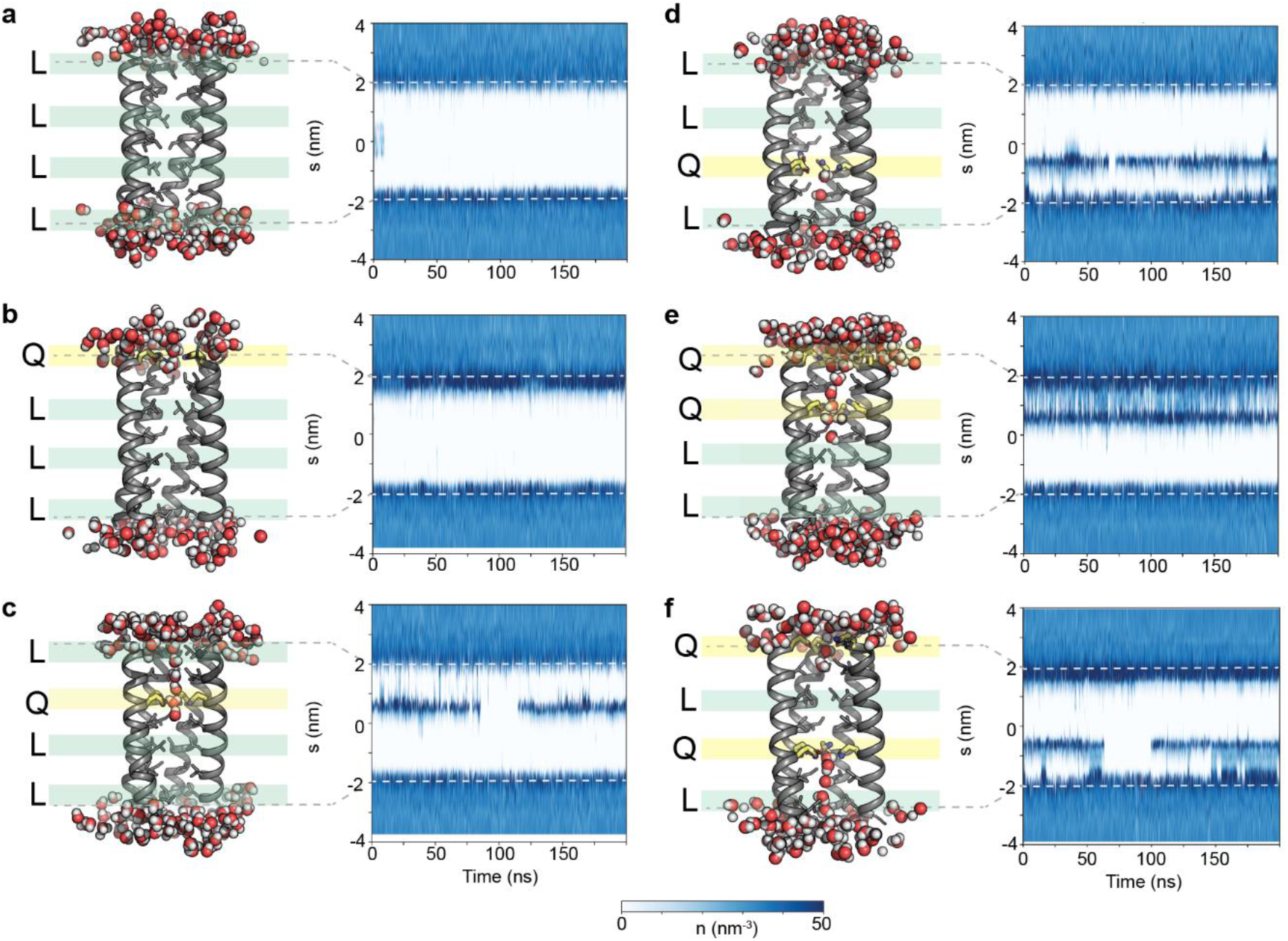
Permeation of water into the designed channels correlates with the position of the luminal Gln residues. MD simulations were analyzed using the Channel Annotation Package ^44^. Plots of pore water density versus time reveal no water permeation into the hydrophobic pore in **a**, LLLL or **b**, QLLL. Pentamers with Gln in the second and third layers of the channel, including **c**, LQLL, **d**, LLQL, **e**, QQLL, and **f**, QLQL, have strong water density around the polar Gln site and flickering water molecules in the shorter apolar segment leading up to the mutation. However, the longer apolar segment remains dehydrated. Fifth helix in all figures removed for clarity.

## Structure of design proton channels

The structures of the five pentameric Gln-containing pentamers were determined by X-ray crystallography. Although they crystallized in different space groups (Extended Data Table 2), their backbone structures were nearly identical with backbone a RMSD to the starting LLLL pentamer ranging from 0.25 to 0.32 Å (Fig. 4a). The Gln sidechains converge in layers near the center of the bundle, where they form sidechain-sidechain and sidechain-backbone hydrogen bonds (Fig. 4 and Supplementary Figs. S7-S16). As expected, they also surround puncta of electron density, which were well modeled as water molecules at full occupancy (Extended Data Fig. 1). The Gln sidechains are not fully symmetric and different conformations are seen in the individual monomeric units of the pentamers. They also have higher Debye Waller factors than the surrounding main-chain and pore-lining sidechains. These findings are in good agreement with the hydration of Gln sidechains observed in MD simulations.

**Fig. 4.**
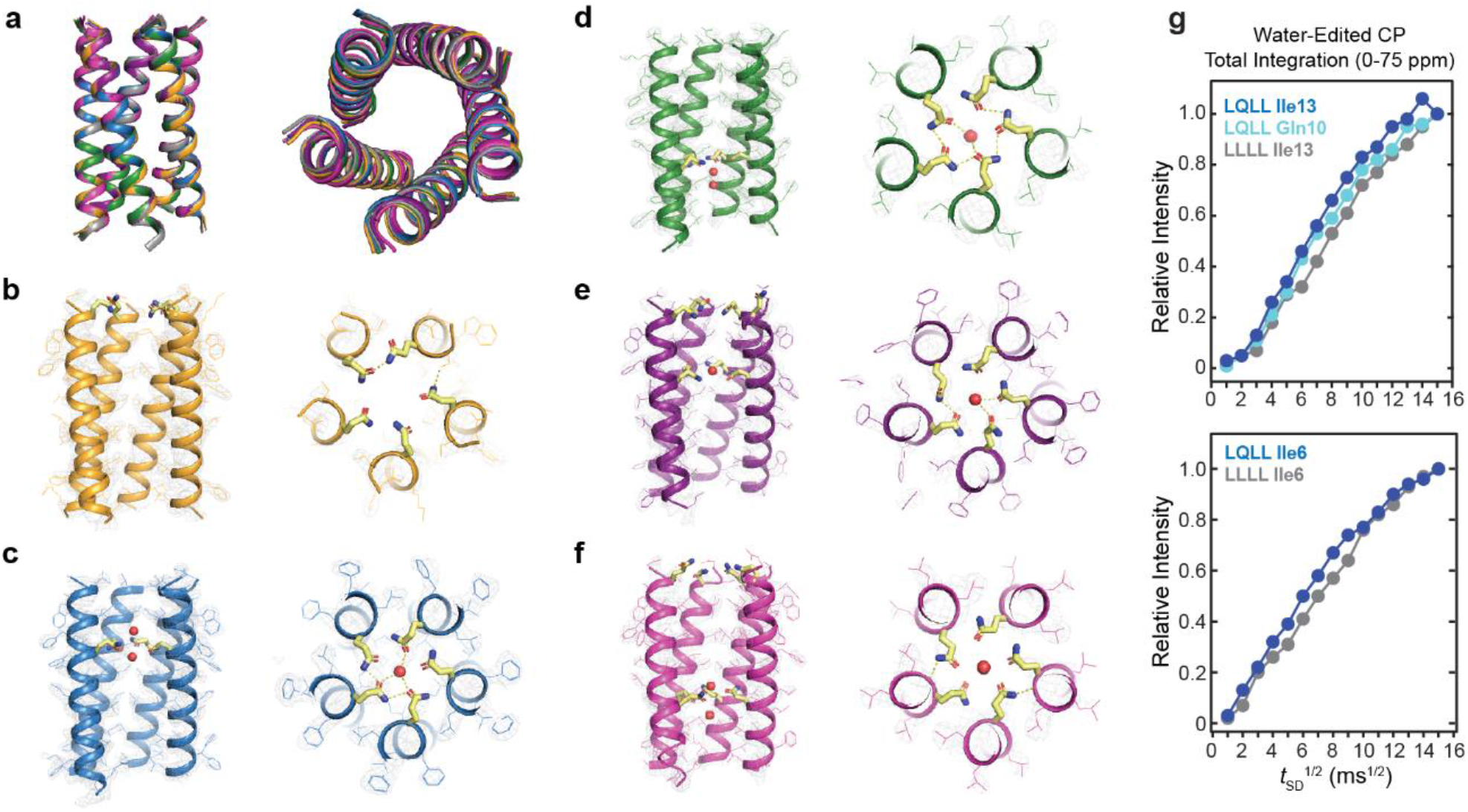
Crystal structures of the designed channels demonstrate the introduction of polar Gln residues mediates stable water pockets. The X-ray structures of the designed channels are within < 0.4 Å rmsd of the original design template, **a**, LLLL (pdb 6mct, gray), **b**, QLLL (pdb 7udy), **c**, LQLL (pdb 7udz), **d**, LLQL (pdb 7udv), **e**, QQLL (pdb 7udw), and **f**, QLQL (pdb 7udx). Fifth helix in side-view figures removed for clarity. **g,** Water buildup curves for uniformly-labeled ^13^C, ^15^N Ile13, Ile6, and Gln10 in LLLL and LQLL peptides. Both the Ile6 and Ile13 sites in the LQLL sample show faster water buildup than the corresponding sites in the LLLL sample.

To confirm the hydration observed from MD simulations and crystal structures, we measured water-edited ^13^C solid state NMR (ssNMR) spectra of a series of LLLL and LQLL peptides uniformly labeled with ^13^C, ^15^N at the Ile6 or Ile13 positions. As expected from the MD simulations, magnetization transfer from water was more efficient for Ile13 in LQLL than in LLLL for all atoms (Fig. 4g, Extended Data Fig. 2). Experiments with labeled Gln10 also show significant magnetization transfer from water to the PLS (Fig. 4g). Thus, the relative water accessibility observed for these pentamers in hydrated phospholipid bilayers is in good agreement with MD simulations and crystal structures.

However, all three of these methods probe equilibrium hydration in the absence of an excess proton in the pore. The Grotthuss proton hopping process requires consideration of covalent bond making and breaking as the proton moves from water to water. To simulate this process, we turned to RMD simulations.

## Multiscale reactive MD simulations from structures

While classical simulations are useful for evaluating the hydration of the channel in the absence of an excess proton, they do not account for changes in chemical bonding such as occurs with the Grotthuss shuttling mechanism caused by protons permeating the channel ^8,10^. We therefore turned to multiscale reactive MD (MS-RMD) simulations, which explicitly simulate the entire dynamic trajectory of proton translocation, including the covalent bonding rearrangements and transfer of protons as they pass between water molecules ^45–47^. While MS-RMD is significantly faster than explicit quantum mechanical (*ab initio* MD) calculations, the calculation of proton transport through an entire TM protein via MS-RMD is still computationally very intensive so we confined our focus to a comparison of LLLL and LQLL from the crystal structures. We used two collective variables (CVs) to enable enhanced free energy sampling in the RMD simulations: the position of the “center of excess charge” (CEC), which tracks the translocation of a proton charge defect along the pore ^45–47^, and a water wire connectivity parameter, ϕ, which quantifies the number and connectivity of hydrogen-bonded waters in the pore and also associated with the excess proton structure ^8^. A value of ϕ of 0 corresponds to a dry pore, and 1 to a pore with a fully connected water wire spanning the pore and containing an excess proton.

The computed potentials of mean force (PMFs) for LLLL vs. LQLL predict that the saddle in the two-dimensional free energy “landscape” as a function of the two CVs for proton translocation is prohibitively high in LLLL, but greatly lowered through the introduction of the single Gln in LQLL (Figs. 5a,b and Supplementary Fig. S17). The landscape and its saddle region for LQLL (rescaled in 5c to allow easier viewing of its contours) describes the free energy of an excess proton at varying positions along the channel axis (x-axis, Fig. 5a–c) and varying degrees of protonated water wire formation (y-axis). The asymmetric location of the Gln-containing PLS in LQLL separates the overall channel into a “short apolar path” near the N-terminus (right side in Fig. 5a,b, top in Fig. 5d) and a “long apolar path” near the C-terminus (left in Fig. 5a,b, bottom in Fig. 5d). The lowest energy pathway from the exterior to the PLS through the short apolar path (points 1 to 3 to 4A to 5 in Fig. 5c and 5d) features water-wires that are fully formed through only this region of the channel and then “flip” to translocate the proton through. The lower free energy saddle predicts rapid proton transport through the short apolar path. The arrival of a proton in the lower portion of the PLS was also seen to sometimes induce cooperative, fully formed water wires running through the entire long apolar path (points 1 to 3 to 4B to 5 in Fig. 5c and 5d, and Supplementary Fig. S18). The transmission of protons through this region was thus computed to proceed via two energetically similar pathways (Fig. 5c), which differ in whether water wires are present or absent in the shorter path as the proton moves through the longer path (i.e. a fully connected water wire spanning the length of the channel). It is important to note that the broad saddle region along the y-axis (the water wire connectivity) relative to the depth and narrowness of the wells for the proton entry and exit regions (between point 4A to 4B in Fig. 5c) indicates a large positive entropy change for the excess proton moving into the saddle region. This is a highly favorable entropy of activation and one must therefore not interpret the “barrier” for the proton transport as simply the one-dimensional trace along either one of the white lines in Fig. 5c, but rather the mechanism must be considered as an average over many such paths along that vertical saddle, and hence a quite favorable proton translocation behavior as seen in the experiments. In summary, RMD of LLLL versus LQLL predicts that the introduction of a Gln-rich PLS one third of the way through the pore should dramatically enhance transmission of protons, via the formation of proton-induced water wires.

**Fig. 5.**
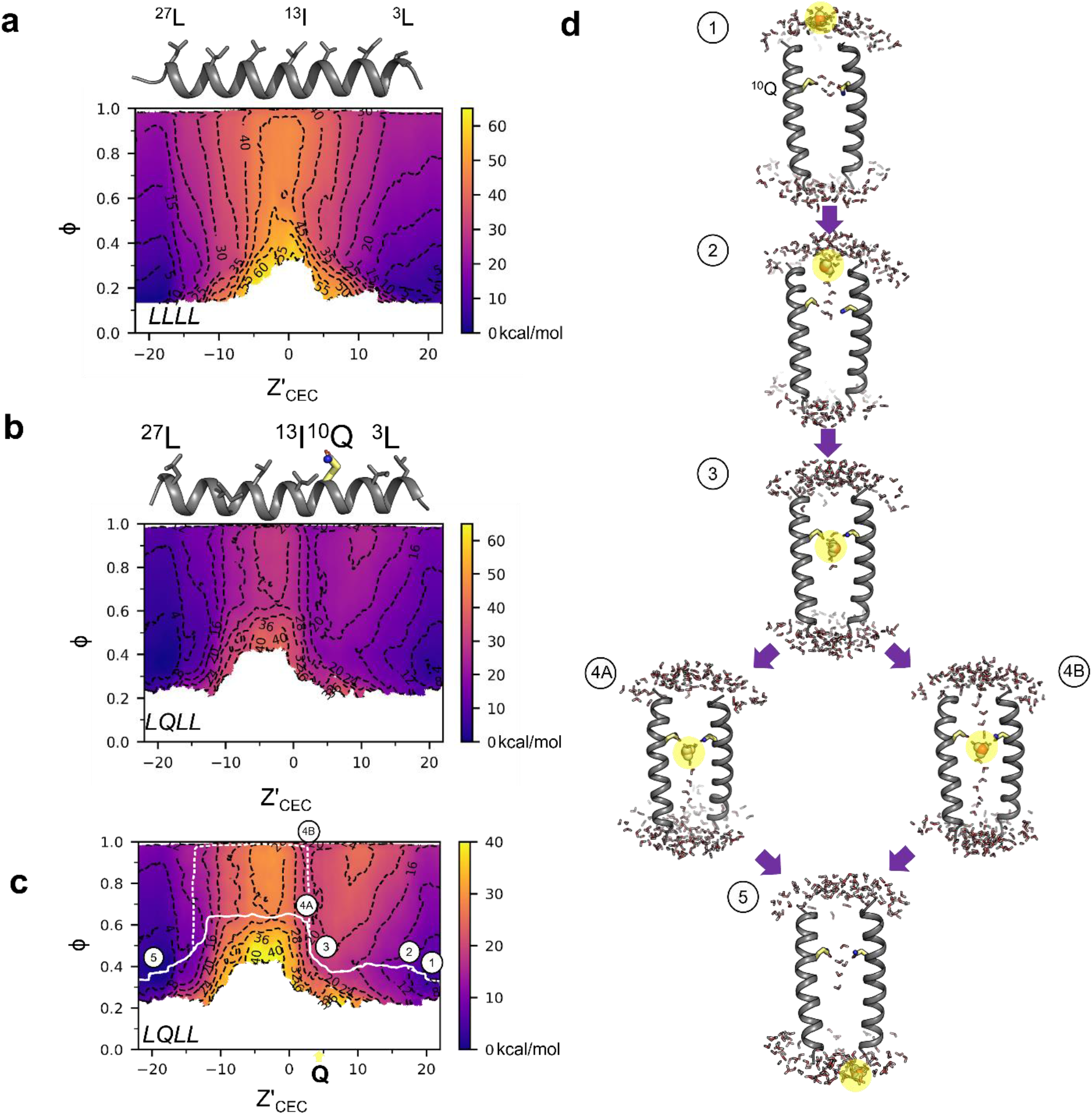
MS-RMD predicts that introduction of the PLS enables the formation of proton-conducive transient water wires. **a**, 2D-PMF of LLLL shows high barrier when the proton is at Z’_CEC_ = 0 Å, or the center of mass of the channel at the Ile13 alpha-carbons. Z’_CEC_ in **a-c** are in units of Å. **b**, Addition of the Gln residue at +4 Å in LQLL shifts the barrier to the C-terminal side of the channel and decreases the barrier height by ~20 kcal/mol. **c**, The two lowest mean free energy paths (MFEPs, white solid and dashed lines), derived from string theory (see Methods) through the LQLL channel. Note the scale change on the color bar in 5b versus 5c. **d**, Snapshots along the two pathways for LQLL (from panel c) reveal the mechanism of proton-induced water wires mediating proton translocation. The most hydronium-like structures are highlighted in yellow. Only two helices are represented for clarity.

## Function through proton flux measurements

To demonstrate the ability of these designed channels to transport protons, we experimentally measured the flux of protons driven by an electrical gradient. We used a vesicle assay (adapted from ^48,49^) that uses a chemiosmotically-induced electrical potential to drive carrier-mediated translocation of protons into phospholipid vesicles. In this assay, vesicles containing K^+^ buffer are rapidly diluted into Na^+^ buffer, creating a chemical potential across the bilayer. Valinomycin, a K^+^ carrier that is highly selective for K^+^ over Na^+^, is then added to allow K^+^ to diffuse down its chemical potential out of the vesicle, thereby creating a TM electrical potential (Fig. 6a and Extended Data Fig. 3). If a proton carrier or channel is present in the bilayer, protons will then follow the induced electrical potential and diffuse against a growing concentration gradient of protons into the vesicle leading to a drop in the interior pH (pH_in_), which is detected by the pH-sensitive fluorescent dye (Fig. 6a). It is noteworthy that this system requires that the vesicles be non-leaky and highly impermeable to Na^+^, otherwise the Na^+^ will diffuse into the vesicle dissipating the electrical potential (Extended Data Fig. 4). Figure 5c shows proton conduction data for the well-characterized proton channel, M2, a proton-selective viroporin from the influenza A virus ^50^. Induction of an electrical gradient by addition of valinomycin (t = 120 s) leads to a change in pH_in_ as expected from M2’s proton-selectivity, but no change in pH_in_ for empty vesicles which contain no protein (Fig. 6b). Following the addition of valinomycin, the protonophore, carbonyl cyanide m-chlorophenyl hydrazone (CCCP), is added (Extended Data Fig. 3); the observed additional bolus of proton flux assures that an electrical and pH gradients have been maintained throughout the experiment, even on the order of hours (Extended Data Figs. 4–7, Supplementary Figs. S19-S23).

**Fig. 6.**
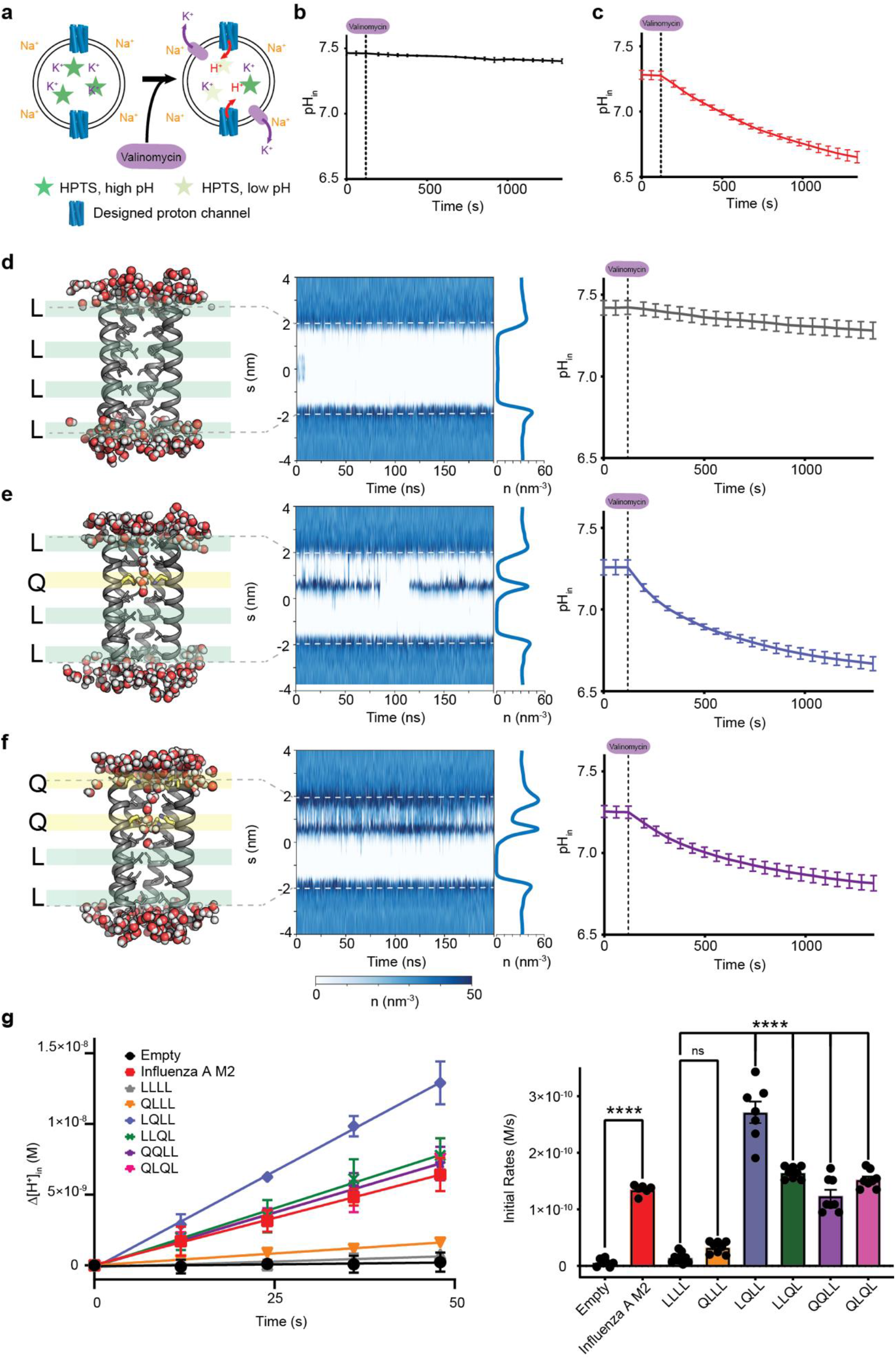
Designed channels selectively move protons across the membrane. **a**, Schematic for proton flux assays using a vesicle-entrapped pH-sensitive fluorescent dye, 8-hydroxypyrene-1,3,6-trisulfonic acid (HPTS). At t=0 the pH is the same inside and outside of the vesicles. **b**, Following addition of valinomycin, the pH_in_ of empty vesicles does not change, because there is no proton channel or carrier included. **c**, When a proton channel is present, like the influenza A M2 channel shown in this panel, addition of valinomycin enables the transport of protons down the electrochemical gradient created by the efflux of potassium. This results in a significant decrease in the pH_in_ over time as protons move into the vesicle up a concentration gradient. Water density profiles and snapshots (fifth helix removed for clarity) are shown along with the corresponding water density plots, obtained through Channel Annotation Package software ^44^, and proton flux assays for **d**, LLLL, **e**, LQLL, and **f**, QQLL indicates that addition of polar Gln near the middle of the channel enables water permeation events into the pore which facilitates proton-selective transport. **g**, Change in the H^+^ concentration of representative samples upon addition of valinomycin at t=0. Fitting of the initial rates shows that LQLL, LLQL, QQLL, and QLQL have significant proton transport activity (p < 0.0001). All data in **a-f** presented with standard deviation bars. Data in **g** shown with standard deviation (left) and standard error (right) bars.

Using this assay, we determined that LLLL (*l_obs_* = 33.9 ± 0.3 Å) had no significant conductance over background, and QLLL (*l_obs_* = 32.0 ± 0.6 Å) had a very low conduction that was not significantly different from LLLL (Fig. 6g). In contrast, the remaining pentamers with apolar tracks of approximately 20 Å (LQLL, LLQL, QQLL and QLQL) showed significant proton conduction well above background, reaching rates on par with M2 at equivalent symmetrical pH_in_ and pH_out_ and peptide/lipid ratios (Fig. 6g). The conductance rate of these four functional proton channels were all highly similar (within a factor of two), which is consistent with their similar hydrophobic lengths. This finding was confirmed in 6-9 independent replicates for each channel (p < 0.0001) (Table S3). Given that the pH is 7.5 ([H^+^] = 10^-7.5^ M) and [Na^+^]_out_ is 0.16 M at the beginning of the experiment the selectivity of the channels for protons over Na^+^ must be at least 10^6^-fold.

## Discussion

We have successfully engineered minimalist proton channels that enable us to explore the roles transient water wires through apolar regions play in proton selectivity and conduction. In keeping with our minimalist strategy, a ring of Gln sidechains was chosen as the PLS. In future work, ionizable residues in the pore would allow for pH and metal ion ^51,52^ gating and modulation of the proton conductance. Moreover, although the peptide pentamers were randomly distributed in the membrane in our preparations (Extended Data Fig. 8), methods to allow unidirectional insertion of the channels would provide information concerning the symmetry of conduction. Previous theoretical work has predicted that the arrival of a proton at a loading site can trigger transient water wires that span surprisingly long distances ^11^ which we show here can be as long as ~20 Å. While this was predicted based on MS-RMD, our classical MD simulations, which do not explicitly consider a mobile Grotthuss shuttling excess proton, predicted that water wires could only span on the order 10 Å from the exterior bulk water to a position within the Gln pentad. Thus, while classical simulations are quite useful for identifying pre-existing water wires that occasionally flicker on in the neutral state (Fig. 1c), they are intrinsically less suited for identifying water wires that are induced as a proton enters the channel (Fig. 1d). The M2 proton channel from influenza A virus ^53,54^ and Hv1 ^55–57^ use electrical or chemical gradients to drive vectorial Grothuss proton conduction across apolar constrictions within channels. In other cases, such as bacteriorhodopsin ^13^,^21–23^, light absorption leads to release of protons that induce formation of water wires. Similarly, proton transport to the proton loading site of cytochrome c oxidase traverses an apolar region ^58–62^; classical and MS-RMD simulations have revealed that transient water structures facilitate this important step of this proton pump ^63,64^. In each case, proton conduction occurs along water wires that are impervious to larger ions. By contrast, many channels that conduct larger ions have apolar pores that are dry in the “off” state but become significantly more enlarged and hydrated in response to a larger conformational gating transition ^16–19,65^.

During proton conduction, each water in a water wire changes the direction of its dipole as protons transfer from one water molecule to the next. The orientation of the waters must be reset to regenerate the initial polarization before a new proton can be transported. Water wires in restricted hydrophobic environments are well suited for stabilizing both polarizations because they do not form strong dipolar interactions with the pore that would bias their orientations. Also, the Gln sidechains are relatively mobile in our crystal structures as assessed from the B-factors, MD and MS-RMD (Supplementary Fig. S24), which shows rapid conformational fluctuations. This observation is also supported by broader peak widths in the ssNMR spectra for LQLL versus LLLL (Supplementary Fig. 25). This behavior contrasts with the requirements for water channels, like aquaporins, which feature stable water-binding sites with strong polarization that undermine orientations of the hydrogen-bonded networks and dipolar switching that would otherwise enable proton-selective transport ^66,67^.

In this work, it was necessary to move beyond the static structures to include the dynamic processes required for proton migration, including both protein and water dynamics, which *de novo* design generally ignores in favor of structural stability and computational efficiency. MS-RMD allowed even deeper consideration of the bond-making/breaking steps required for Grotthuss proton migration through water wires. While computational speed is currently too slow to incorporate RMD into early stages of protein design, it can clearly provide an important filter to assess potential designs. Indeed, Mondal and coworkers have used an empirical valence bond approach to screen combinatorial libraries of enzyme variants ^68^.

Our work also highlights enabling design principles for the development of new proton-conductive materials. We showed that close positioning of a PLS proximal to a dry pore resulted in channels that are highly selective for protons. Our minimalist designs show that the PLS need not be elaborate in design. Indeed, the present work was inspired in part by experimental and computational studies of hard materials composed of carbon nanotubes ^69,70^. Our design principles also have implications for design of soft materials proton-selective membranes. For example, Jiang and coworkers have designed proton-selective copolymers consisting of apolar segments interspersed with occasional polar ethylene glycol units ^71^, bearing similarities to the two-component design of our channels. Our current work extends a computational approach to mechanistically interrogate and design materials with even greater efficiency and selectivity. Indeed, because our designs are based on fundamental physical chemical principles and molecular rather than bioinformatic algorithms they are not limited to production of natural proteins or synthetic peptides. Instead, they should be able to translate to the design of novel non-proteinaceous molecular assemblies and polymers for applications ranging from water purification to energy storage and utilization.

## Acknowledgements

Diffraction data was collected at the GM/CA@APS and ALS BL 8.3.1. GM/CA@APS is supported by the National Cancer Institute (ACB-12002) and the National Institute of General Medical Sciences (AGM-12006, P30GM138396), and the Eiger 16M detector by NIH S10 OD012289. We also acknowledge the Advanced Photon Source, supported by U.S. Department of Energy (DE) contract DE-AC02-06CH11357, and beamline 8.3.1 at the Advanced Light Source operated by the University of California at San Francisco with support from National Institutes of Health (NIH; R01 GM124149 and P30 GM124169), Plexxikon Inc. and the Integrated Diffraction Analysis Technologies program (US Department of Energy Office of Biological and Environmental Research. The Advanced Light Source at Lawrence Berkeley National Laboratory is supported by DE-AC02-05CH11231.

## Author contributions

H.T.K. designed the channels, ran flux measurements, crystallized and collected the X-ray diffraction data, and ran and analyzed classical MD simulations. L.C.W. ran the MS-RMD simulations, L.C.W. and G.A.V. analyzed the data. N.H.S. conducted the solid-state NMR experiments. N.H.S. and M. H. analyzed the NMR data. M.M. ran classical MD simulations. J.L.T, J.M.N., and L.L. processed and refined the crystal structures. H.T.K. and W.F.D. analyzed experimental data. All authors contributed to data analysis and writing the manuscript.

## Data and materials availability

Coordinates and data files have the PDB with accession codes 7UDY (QLLL), 7UDZ (LQLL), 7UDV (LLQL), 7UDW (QQLL), and 7UDX (QLQL). Materials are available from the authors on request.

## References and notes

1 Moriyama, Y. & Futai, M. H+-ATPase, a primary pump for accumulation of neurotransmitters, is a major constituent of brain synaptic vesicles. Biochemical and biophysical research communications 173, 443–448, doi:10.1016/s0006-291x(05)81078-2 (1990).

2 Nishi, T. & Forgac, M. The vacuolar (H+)-ATPases--nature’s most versatile proton pumps. Nature reviews. Molecular cell biology 3, 94–103, doi:10.1038/nrm729 (2002).

3 Mitchell, P. Coupling of phosphorylation to electron and hydrogen transfer by a chemi-osmotic type of mechanism. Nature 191, 144–148, doi:10.1038/191144a0 (1961).

4 Nicholls, D. G. Mitochondrial ion circuits. Essays in biochemistry 47, 25–35, doi:10.1042/bse0470025 (2010).

5 Diering, G. H. & Numata, M. Endosomal pH in neuronal signaling and synaptic transmission: role of Na(+)/H(+) exchanger NHE5. Frontiers in physiology 4, 412, doi:10.3389/fphys.2013.00412 (2014).

6 Agmon, N. The Grotthuss mechanism. Chemical Physics Letters 244, 456–462, doi:10.1016/0009-2614(95)00905-j (1995).

7 Calio, P. B., Li, C. & Voth, G. A. Resolving the Structural Debate for the Hydrated Excess Proton in Water. Journal of the American Chemical Society 143, 18672–18683, doi:10.1021/jacs.1c08552 (2021).

8 Li, C. & Voth, G. A. A quantitative paradigm for water-assisted proton transport through proteins and other confined spaces. Proceedings of the National Academy of Sciences of the United States of America 118, doi:10.1073/pnas.2113141118 (2021).

9 Wraight, C. A. Chance and design--proton transfer in water, channels and bioenergetic proteins. Biochimica et biophysica acta 1757, 886–912, doi:10.1016/j.bbabio.2006.06.017 (2006).

10 Decoursey, T. E. Voltage-gated proton channels and other proton transfer pathways. Physiol Rev 83, 475–579, doi:10.1152/physrev.00028.2002 (2003).

11 Peng, Y., Swanson, J. M., Kang, S. G., Zhou, R. & Voth, G. A. Hydrated Excess Protons Can Create Their Own Water Wires. The journal of physical chemistry. B 119, 9212–9218, doi:10.1021/jp5095118 (2015).

12 Banh, R. et al. Hydrophobic gasket mutation produces gating pore currents in closed human voltage-gated proton channels. Proceedings of the National Academy of Sciences of the United States of America 116, 18951–18961, doi:10.1073/pnas.1905462116 (2019).

13 Garczarek, F. & Gerwert, K. Functional waters in intraprotein proton transfer monitored by FTIR difference spectroscopy. Nature 439, 109–112, doi:10.1038/nature04231 (2006).

14 Kaur, D., Khaniya, U., Zhang, Y. & Gunner, M. R. Protein Motifs for Proton Transfers That Build the Transmembrane Proton Gradient. Frontiers in chemistry 9, 660954, doi:10.3389/fchem.2021.660954 (2021).

15 Kalra, A., Garde, S. & Hummer, G. Osmotic water transport through carbon nanotube membranes. Proceedings of the National Academy of Sciences of the United States of America 100, 10175–10180, doi:10.1073/pnas.1633354100 (2003).

16 Ben-Abu, Y., Zhou, Y., Zilberberg, N. & Yifrach, O. Inverse coupling in leak and voltage-activated K+ channel gates underlies distinct roles in electrical signaling. Nature structural & molecular biology 16, 71–79, doi:10.1038/nsmb.1525 (2009).

17 Jensen, M. O. et al. Principles of conduction and hydrophobic gating in K+ channels. Proceedings of the National Academy of Sciences of the United States of America 107, 5833–5838, doi:10.1073/pnas.0911691107 (2010).

18 Aryal, P., Sansom, M. S. & Tucker, S. J. Hydrophobic gating in ion channels. J Mol Biol 427, 121–130, doi:10.1016/j.jmb.2014.07.030 (2015).

19 Zhu, F. & Hummer, G. Drying transition in the hydrophobic gate of the GLIC channel blocks ion conduction. Biophysical journal 103, 219–227, doi:10.1016/j.bpj.2012.06.003 (2012).

20 Rasaiah, J. C., Garde, S. & Hummer, G. Water in nonpolar confinement: from nanotubes to proteins and beyond. Annual review of physical chemistry 59, 713–740, doi:10.1146/annurev.physchem.59.032607.093815 (2008).

21 Wang, T. et al. Deprotonation of D96 in bacteriorhodopsin opens the proton uptake pathway. Structure 21, 290–297, doi:10.1016/j.str.2012.12.018 (2013).

22 Weinert, T. et al. Proton uptake mechanism in bacteriorhodopsin captured by serial synchrotron crystallography. Science 365, 61–65, doi:10.1126/science.aaw8634 (2019).

23 Freier, E., Wolf, S. & Gerwert, K. Proton transfer via a transient linear water-molecule chain in a membrane protein. Proceedings of the National Academy of Sciences of the United States of America 108, 11435–11439, doi:10.1073/pnas.1104735108 (2011).

24 Regan, L. & DeGrado, W. F. Characterization of a helical protein designed from first principles. Science 241, 976–978, doi:10.1126/science.3043666 (1988).

25 Walsh, S. T., Cheng, H., Bryson, J. W., Roder, H. & DeGrado, W. F. Solution structure and dynamics of a de novo designed three-helix bundle protein. Proceedings of the National Academy of Sciences of the United States of America 96, 5486–5491, doi:10.1073/pnas.96.10.5486 (1999).

26 Kuhlman, B. et al. Design of a novel globular protein fold with atomic-level accuracy. Science 302, 1364–1368, doi:10.1126/science.1089427 (2003).

27 Vorobieva, A. A. et al. De novo design of transmembrane beta barrels. Science 371, doi:10.1126/science.abc8182 (2021).

28 Yang, C. et al. Bottom-up de novo design of functional proteins with complex structural features. Nat Chem Biol 17, 492–500, doi:10.1038/s41589-020-00699-x (2021).

29 Polizzi, N. F. & DeGrado, W. F. A defined structural unit enables de novo design of small-molecule-binding proteins. Science 369, 1227–1233, doi:10.1126/science.abb8330 (2020).

30 Cao, L. et al. De novo design of picomolar SARS-CoV-2 miniprotein inhibitors. bioRxiv: the preprint server for biology, doi:10.1101/2020.08.03.234914 (2020).

31 Fleishman, S. J. et al. Computational design of proteins targeting the conserved stem region of influenza hemagglutinin. Science 332, 816–821, doi:10.1126/science.1202617 (2011).

32 Jiang, L. et al. De novo computational design of retro-aldol enzymes. Science 319, 1387–1391, doi:10.1126/science.1152692 (2008).

33 Lassila, J. K., Privett, H. K., Allen, B. D. & Mayo, S. L. Combinatorial methods for small-molecule placement in computational enzyme design. Proceedings of the National Academy of Sciences of the United States of America 103, 16710–16715, doi:10.1073/pnas.0607691103 (2006).

34 Polizzi, N. F. et al. De novo design of a hyperstable non-natural protein-ligand complex with sub-A accuracy. Nature chemistry 9, 1157–1164, doi:10.1038/nchem.2846 (2017).

35 Leaver-Fay, A. et al. ROSETTA3: an object-oriented software suite for the simulation and design of macromolecules. Methods in enzymology 487, 545–574, doi:10.1016/B978-0-12-381270-4.00019-6 (2011).

36 Koga, N. et al. Principles for designing ideal protein structures. Nature 491, 222–227, doi:10.1038/nature11600 (2012).

37 Scott, A. J. et al. Constructing ion channels from water-soluble alpha-helical barrels. Nature chemistry 13, 643–650, doi:10.1038/s41557-021-00688-0 (2021).

38 Xu, C. et al. Computational design of transmembrane pores. Nature 585, 129–134, doi:10.1038/s41586-020-2646-5 (2020).

39 Joh, N. H. et al. De novo design of a transmembrane Zn2+-transporting four-helix bundle. Science 346, 1520–1524, doi:10.1126/science.1261172 (2014).

40 Lu, P. et al. Accurate computational design of multipass transmembrane proteins. Science 359, 1042–1046, doi:10.1126/science.aaq1739 (2018).

41 Thomaston, J. L. et al. X-ray Crystal Structure of the Influenza A M2 Proton Channel S31N Mutant in Two Conformational States: An Open and Shut Case. Journal of the American Chemical Society 141, 11481–11488, doi:10.1021/jacs.9b02196 (2019).

42 Saotome, K. et al. Structures of the otopetrin proton channels Otop1 and Otop3. Nature structural & molecular biology 26, 518–525, doi:10.1038/s41594-019-0235-9 (2019).

43 Mravic, M. et al. Packing of apolar side chains enables accurate design of highly stable membrane proteins. Science 363, 1418–1423, doi:10.1126/science.aav7541 (2019).

44 Klesse, G., Rao, S., Sansom, M. S. P. & Tucker, S. J. CHAP: A Versatile Tool for the Structural and Functional Annotation of Ion Channel Pores. J Mol Biol 431, 3353–3365, doi:10.1016/j.jmb.2019.06.003 (2019).

45 Lee, S., Liang, R., Voth, G. A. & Swanson, J. M. Computationally Efficient Multiscale Reactive Molecular Dynamics to Describe Amino Acid Deprotonation in Proteins. Journal of chemical theory and computation 12, 879–891, doi:10.1021/acs.jctc.5b01109 (2016).

46 Knight, C., Lindberg, G. E. & Voth, G. A. Multiscale reactive molecular dynamics. J Chem Phys 137, 22A525, doi:10.1063/1.4743958 (2012).

47 Yamashita, T., Peng, Y., Knight, C. & Voth, G. A. Computationally Efficient Multiconfigurational Reactive Molecular Dynamics. Journal of chemical theory and computation 8, 4863–4875, doi:10.1021/ct3006437 (2012).

48 Moffat, J. C. et al. Proton transport through influenza A virus M2 protein reconstituted in vesicles. Biophysical journal 94, 434–445, doi:10.1529/biophysj.107.109082 (2008).

49 Ma, C. et al. Identification of the functional core of the influenza A virus A/M2 proton-selective ion channel. Proceedings of the National Academy of Sciences of the United States of America 106, 12283–12288, doi:10.1073/pnas.0905726106 (2009).

50 Leiding, T., Wang, J., Martinsson, J., DeGrado, W. F. & Arskold, S. P. Proton and cation transport activity of the M2 proton channel from influenza A virus. Proceedings of the National Academy of Sciences of the United States of America 107, 15409–15414, doi:10.1073/pnas.1009997107 (2010).

51 Slope, L. N. & Peacock, A. F. De Novo Design of Xeno-Metallo Coiled Coils. Chemistry, an Asian journal 11, 660–666, doi:10.1002/asia.201501173 (2016).

52 Pinter, T. B. J., Koebke, K. J. & Pecoraro, V. L. Catalysis and Electron Transfer in De Novo Designed Helical Scaffolds. Angewandte Chemie 59, 7678–7699, doi:10.1002/anie.201907502 (2020).

53 Khurana, E. et al. Molecular dynamics calculations suggest a conduction mechanism for the M2 proton channel from influenza A virus. Proceedings of the National Academy of Sciences of the United States of America 106, 1069–1074, doi:10.1073/pnas.0811720106 (2009).

54 Yi, M., Cross, T. A. & Zhou, H. X. A secondary gate as a mechanism for inhibition of the M2 proton channel by amantadine. The journal of physical chemistry. B 112, 7977–7979, doi:10.1021/jp800171m (2008).

55 Ramsey, I. S. et al. An aqueous H+ permeation pathway in the voltage-gated proton channel Hv1. Nature structural & molecular biology 17, 869–875, doi:10.1038/nsmb.1826 (2010).

56 Chamberlin, A. et al. Hydrophobic plug functions as a gate in voltage-gated proton channels. Proceedings of the National Academy of Sciences of the United States of America 111, E273–282, doi:10.1073/pnas.1318018111 (2014).

57 Takeshita, K. et al. X-ray crystal structure of voltage-gated proton channel. Nature structural & molecular biology 21, 352–357, doi:10.1038/nsmb.2783 (2014).

58 Wikstrom, M., Krab, K. & Sharma, V. Oxygen Activation and Energy Conservation by Cytochrome c Oxidase. Chemical reviews 118, 2469–2490, doi:10.1021/acs.chemrev.7b00664 (2018).

59 Hofacker, I. & Schulten, K. Oxygen and proton pathways in cytochrome c oxidase. Proteins: Structure, Function, and Genetics 30, 100–107, doi:10.1002/(sici)1097-0134(199801)30:1<100::aid-prot9>3.0.co;2-s (1998).

60 Wikström, M., Verkhovsky, M. I. & Hummer, G. Water-gated mechanism of proton translocation by cytochrome c oxidase. Biochimica et Biophysica Acta (BBA) - Bioenergetics 1604, 61–65, doi:10.1016/s0005-2728(03)00041-0 (2003).

61 Tashiro, M. & Stuchebrukhov, A. A. Thermodynamic properties of internal water molecules in the hydrophobic cavity around the catalytic center of cytochrome c oxidase. The journal of physical chemistry. B 109, 1015–1022, doi:10.1021/jp0462456 (2005).

62 Goyal, P., Lu, J., Yang, S., Gunner, M. R. & Cui, Q. Changing hydration level in an internal cavity modulates the proton affinity of a key glutamate in cytochrome c oxidase. Proceedings of the National Academy of Sciences of the United States of America 110, 18886–18891, doi:10.1073/pnas.1313908110 (2013).

63 Liang, R., Swanson, J. M. J., Wikstrom, M. & Voth, G. A. Understanding the essential proton-pumping kinetic gates and decoupling mutations in cytochrome c oxidase. Proceedings of the National Academy of Sciences of the United States of America 114, 5924–5929, doi:10.1073/pnas.1703654114 (2017).

64 Liang, R., Swanson, J. M., Peng, Y., Wikstrom, M. & Voth, G. A. Multiscale simulations reveal key features of the proton-pumping mechanism in cytochrome c oxidase. Proceedings of the National Academy of Sciences of the United States of America 113, 7420–7425, doi:10.1073/pnas.1601982113 (2016).

65 Lynch, C. I., Rao, S. & Sansom, M. S. P. Water in Nanopores and Biological Channels: A Molecular Simulation Perspective. Chemical reviews, doi:10.1021/acs.chemrev.9b00830 (2020).

66 Chen, H. et al. Charge delocalization in proton channels, I: the aquaporin channels and proton blockage. Biophysical journal 92, 46–60, doi:10.1529/biophysj.106.091934 (2007).

67 Murata, K. et al. Structural determinants of water permeation through aquaporin-1. Nature 407, 599–605, doi:10.1038/35036519 (2000).

68 Mondal, D., Kolev, V. & Warshel, A. Combinatorial Approach for Exploring Conformational Space and Activation Barriers in Computer-Aided Enzyme Design. ACS Catal 10, 6002–6012, doi:10.1021/acscatal.0c01206 (2020).

69 Tunuguntla, R. H., Allen, F. I., Kim, K., Belliveau, A. & Noy, A. Ultrafast proton transport in sub-1-nm diameter carbon nanotube porins. Nature nanotechnology 11, 639–644, doi:10.1038/nnano.2016.43 (2016).

70 Geng, J. et al. Stochastic transport through carbon nanotubes in lipid bilayers and live cell membranes. Nature 514, 612–615, doi:10.1038/nature13817 (2014).

71 Jiang, T. et al. Single-chain heteropolymers transport protons selectively and rapidly. Nature 577, 216–220, doi:10.1038/s41586-019-1881-0 (2020).

