## Extended Data for "Transient Water Wires Mediate Selective Proton Transport in Designed Channel Proteins"

**Extended Data Table 1. Length ($l_{obs}$) of longer and shorter ($l_{obs,sh}$) hydrophobic stretches in designed channels.** The longest hydrophobic length for all peptides and shorter hydrophobic length ($l_{obs,sh}$) for each peptide LQLL, LLQL, QQLL, and QLQL were calculated from the classical MD simulations (see Materials and Methods).

| **Design** | $l_{obs}$ | $l_{obs,sh}$ |
|---|---|---|
| LLLL | 33.9 ± 0.3 Å | - |
| QLLL | 32.0 ± 0.6 Å | - |
| LQLL | 20.7 ± 0.1 Å | 11.6 ± 0.4 Å |
| LLQL | 22.4 ± 0.1 Å | 9.3 ± 0.5 Å |
| QQLL | 20.9 ± 0.1 Å | 8.2 ± 0.2 Å |
| QLQL | 20.3 ± 0.2 Å | 9.2 ± 0.2 Å |

**Extended Data Table 2. Data collection and refinement statistics for all structures.** All structures were determined from single protein crystals. Data in parentheses denote statistics for outermost shell. All listed data represent isotropic statistics unless denoted by *, which is anisotropic.

| Property | QLLL<br>PDB: 7UDY<br>APS 23-ID-D | LQLL<br>PDB: 7UDZ<br>APS 23-ID-B | LLQL<br>PDB: 7UDV<br>APS 23-ID-D | QQLL<br>PDB: 7UDW<br>ALS 8.3.1 | QLQL<br>PDB: 7UDX<br>ALS 8.3.1 |
|---|---|---|---|---|---|
| **Data Collection** | | | | | |
| Space Group | $C\ 2\ 2\ 2_1$ | C2 | $P\ 2_1\ 2_1\ 2_1$ | $P2_1$ | $C\ 2\ 2\ 2_1$ |
| **Cell dimensions** | | | | | |
| *a, b, c* (Å) | 56.39 84.84 149.56 | 86.77 46.66 66.32 | 52.53 55.36 82.71 | 48.246 100.175 50.498 | 55.72 83.68 147.03 |
| *α, β, γ* (°) | 90.0 90.0 90.0 | 90.00 108.96 90.00 | 90.0 90.0 90.0 | 90.0 110.506 90.0 | 90.0 90.0 90.0 |
| Resolution (Å) | 74.78 – 2.40 (2.53-2.40) | 41.03 – 2.47 (2.61-2.47) | 46.00 – 2.40 (2.46-2.40) | 42.77 – 3.00 (3.18 – 3.00) | 73.24 – 2.99 (3.04-2.99) |
| $R_{merge}$ | 0.155 (1.249) | 0.114 (0.708) | 0.111 (0.826) | 0.222 (1.207) | 0.145 (0.884) |
| < I /σI > | 8.1 (2.2) | 5.5 (1.8) | 7.6 (2.0) | 6.6 (1.8) | 13.4 (3.5) |
| Completeness (%) | 99.5 (97.7) | 98.1 (93.4) | 91.2 (97.3)* | 98.8 (98.5) | 99.9 (100.0) |
| CC1/2 | 0.998 (0.831) | 0.991(0.800) | 0.999 (0.591) | 0.998 (0.815) | 0.999 (0.921) |
| Redundancy | 8.5 (8.5) | 3.2 (3.0) | 3.3 (3.6) | 6.7 (6.4) | 13.1 (13.9) |
| **Refinement** | | | | | |
| Resolution (Å) | 2.40 | 2.47 | 2.40 | 3.00 | 2.99 |
| No. reflections | 14307 | 8896 | 8048 | 8355 | 7234 |
| $R_{work}/R_{free}$ | 0.230/0.251 | 0.248/0.269 | 0.246/0.268 | 0.251/0.286 | 0.231/0.273 |
| **No. atoms** | | | | | |
| Protein | 3098 | 2038 | 2035 | 4093 | 3093 |
| Ligand | 63 | 21 | 0 | 0 | 14 |
| Water | 0 | 4 | 4 | 6 | 0 (6)* |
| ***B*-Factors** | | | | | |
| Protein | 67.13 | 55.05 | 52.91 | 62.97 | 55.87 |
| Ligand/ion | 82.94 | 61.56 | N/A | N/A | 48.21 |
| Water | N/A | 56.71 | 43.92 | 41.17 | N/A (30.74)* |
| **R.M.S. deviations** | | | | | |
| Bond lengths (Å) | 0.001 | 0.002 | 0.003 | 0.004 | 0.003 |
| Bond angles (°) | 0.324 | 0.414 | 0.498 | 0.583 | 0.490 |
| **Ramachandran statistics** | | | | | |
| Outliers (%) | 0 | 0 | 0 | 0 | 0 |
| Allowed (%) | 0 | 0 | 0 | 0.43 | 0 |
| Favored (%) | 100 | 100 | 100 | 99.57 | 100 |

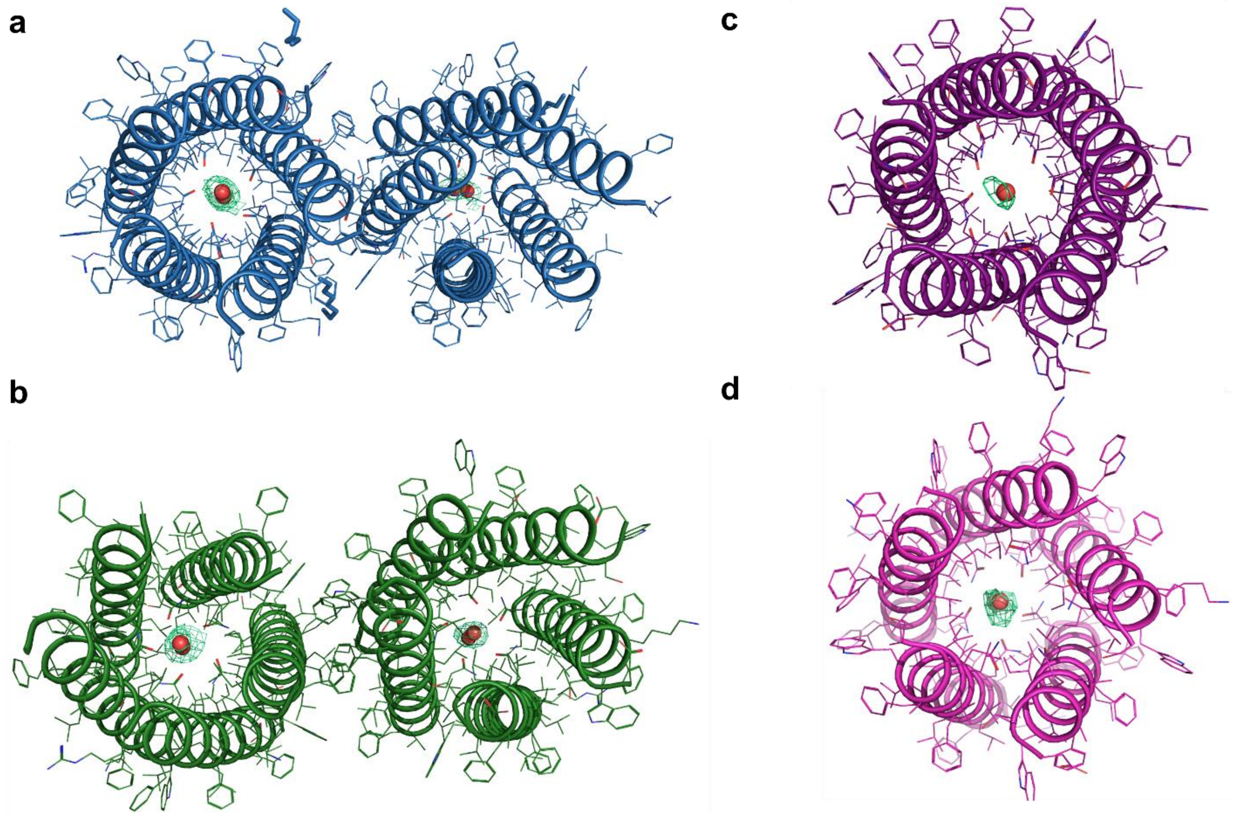


**Extended Data Fig. 1. Composite omit maps (2mFo-DFc) of designed proton channels.** Composite omit maps of the asymmetric unit for **a,** LQLL, **b**, LLQL, and one pentamer from the asymmetric unit for **c,** QQLL, and **d,** QLQL (shown only for the waters for clarity). All contours at σ = 1.0. Omit maps with simulated cartesian annealing were generated using Phenix, using methodology described in Hodel, et al.[104]

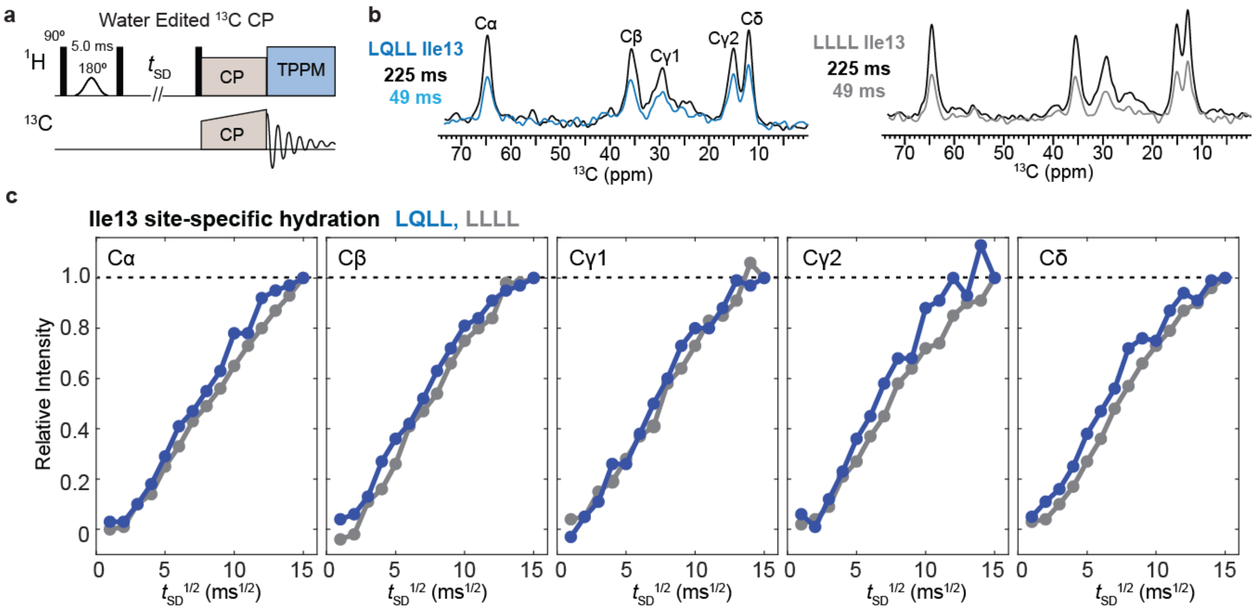


**Extended Data Fig. 2. Pulse diagram and water-edited 13C spectra, water buildup curves of membrane-bound LQLL and LLLL peptides**. **a,** Pulse diagram of water-edited 13C CP experiment. **b,** Representative water-edited 13C spectra of I13 in LQLL and LLLL, measured with 225 ms and 49 ms $^{1}$H mixing. The relative intensities of the 49 ms spectrum to the 225 ms spectrum are higher for LQLL Ile13 than LLLL Ile13, especially for the sidechain Cγ2 and Cδ carbons. **c,** Site-resolved water buildup curves for Ile13 in LQLL and LLLL. For all $^{13}$C sites, LQLL shows a faster water buildup than LLLL, consistent with water molecules in the pore lumen due to the Gln10 PLS.

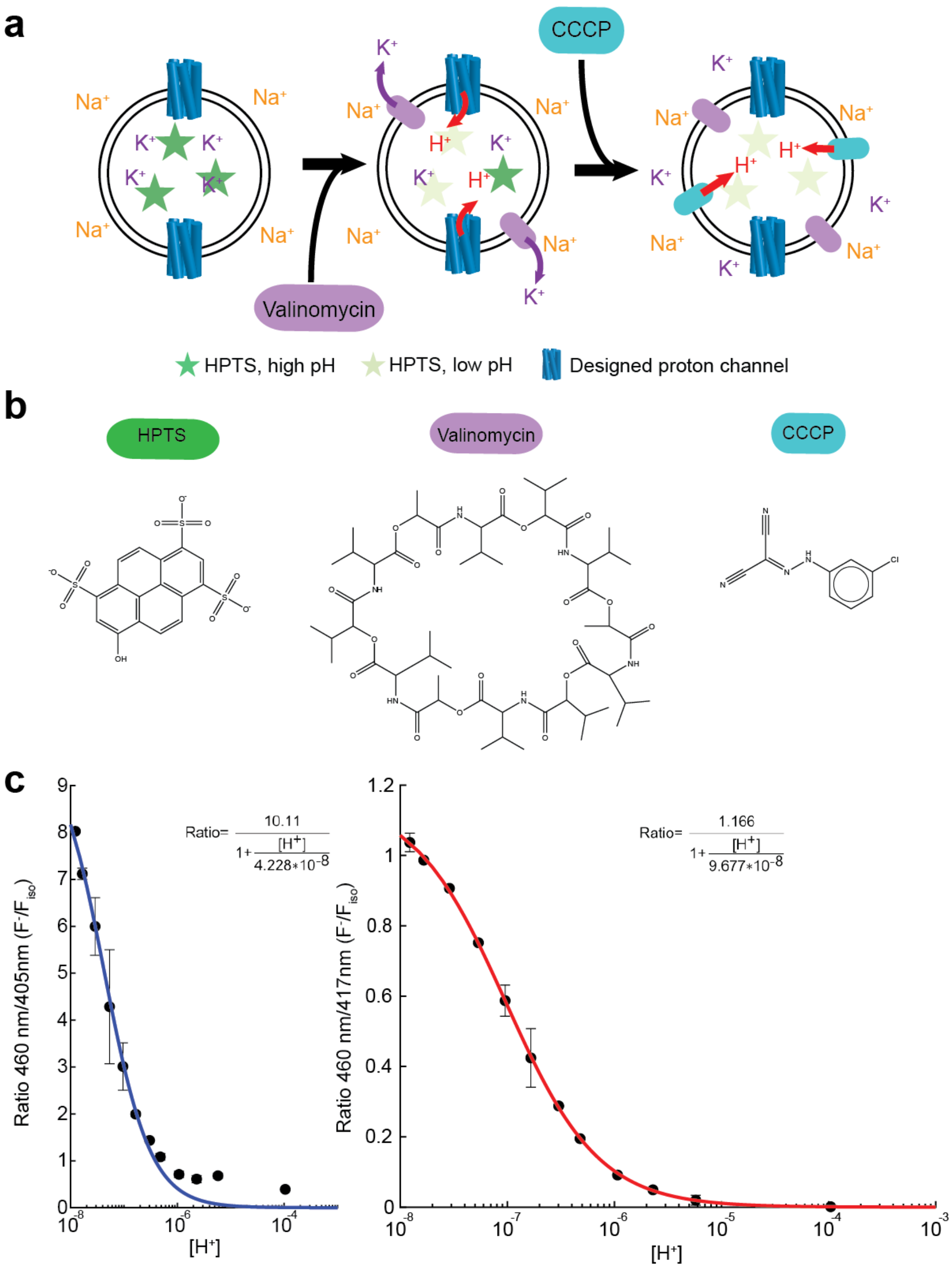


**Extended Data Fig. 3. Overview of proton flux measurements. a,** Full schematic for proton flux measurement including CCCP step, which is included to check vesicle leakiness and confirm proton selectivity. **b,** Chemical structures of key components of vesicle assay. **c,** Calibration curves for HPTS at ~5 µM in 12 solutions of 50 mM $K_2SO_4$, 30 mM $K_2HPO_4$ at different pH values for two plate readers used in data collection process. Unless stated, all data collected with instrument that generated the blue calibration curve. Fits used for downstream data processing shown for each of the two instruments with adjusted R-squared values of 0.9866 and 0.9998 for the left and right curves, respectively.

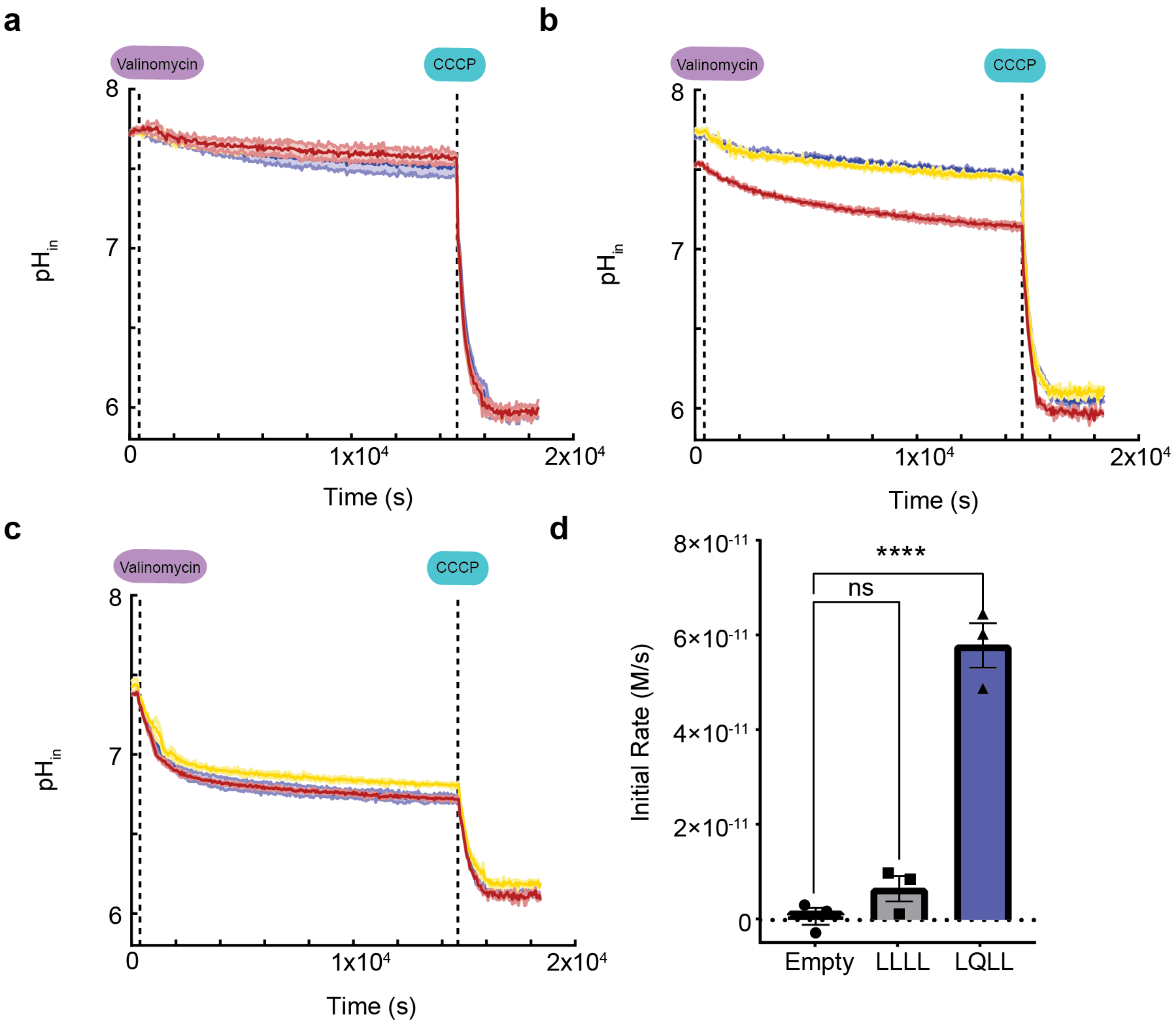


**Extended Data Fig. 4. All proton flux assay data for long kinetics runs.** Long kinetics runs of about 5 hours total for **a,** empty, **b,** LLLL, and **c,** LQLL vesicles. Dotted lines denote times in experiment when valinomycin and CCCP were added to the samples. Three samples of the different conditions were measured in triplicate. These long-time measurements reveal that the vesicles are not significantly leaky to $Na^+$, $K^+$, or $H^+$ and maintain their cargo and assembly over the entire course of the measurement. **d,** From the linear regression fits of the first 220 s following addition of valinomycin, all slopes (which give the initial rates (in M/s)) were used to calculate the mean and standard error. The one-way ANOVA analysis (with Dunn's test) reveals that LLLL rates are not significantly different ($p > 0.05$) when compared to the control empty vesicles. LQLL rates, however, are statistically significant ($p < 0.0001$) when compared to the control empty vesicles using one-way ANOVA analysis with Dunn's test.

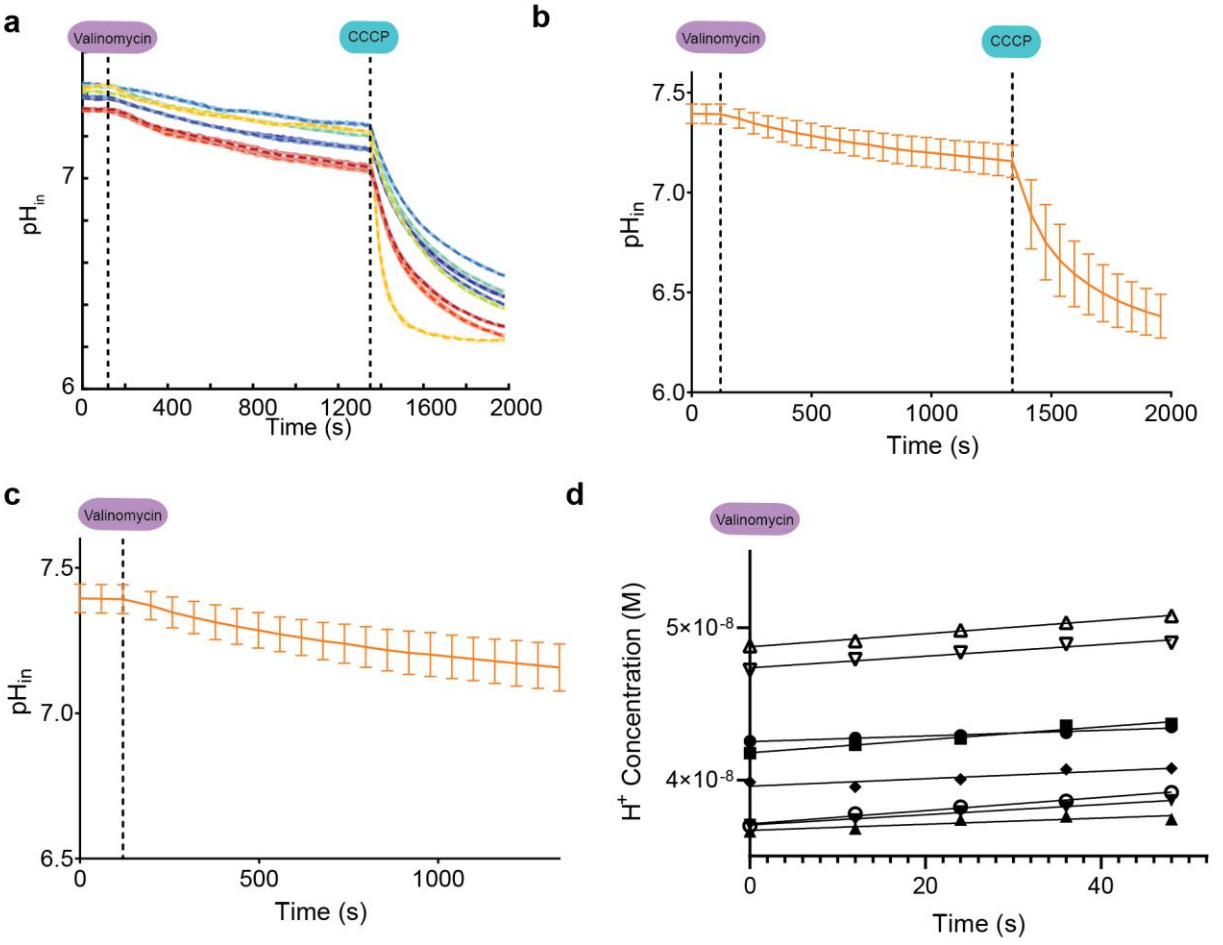


**Extended Data Fig. 5. All proton flux assay data for QLLL vesicle samples.** Nine samples (each run in triplicate with shaded error bars shown) containing 1:500 peptide:lipid ratio; samples were run independently in the assay. **a,** $pH_{in}$ as a function of time throughout the measurement for each independent sample. **b,** Mean and standard deviation for data collection. **c,** Data prior to CCCP addition shows little change in $pH_{in}$ after addition of valinomycin. **d,** Fits for the initial 50 seconds following addition of valinomycin. From the linear regression fits, all slopes (which give the initial rates (in M/s)) were used to calculate mean and standard error presented in Fig. 6g and Supplementary Table S3.

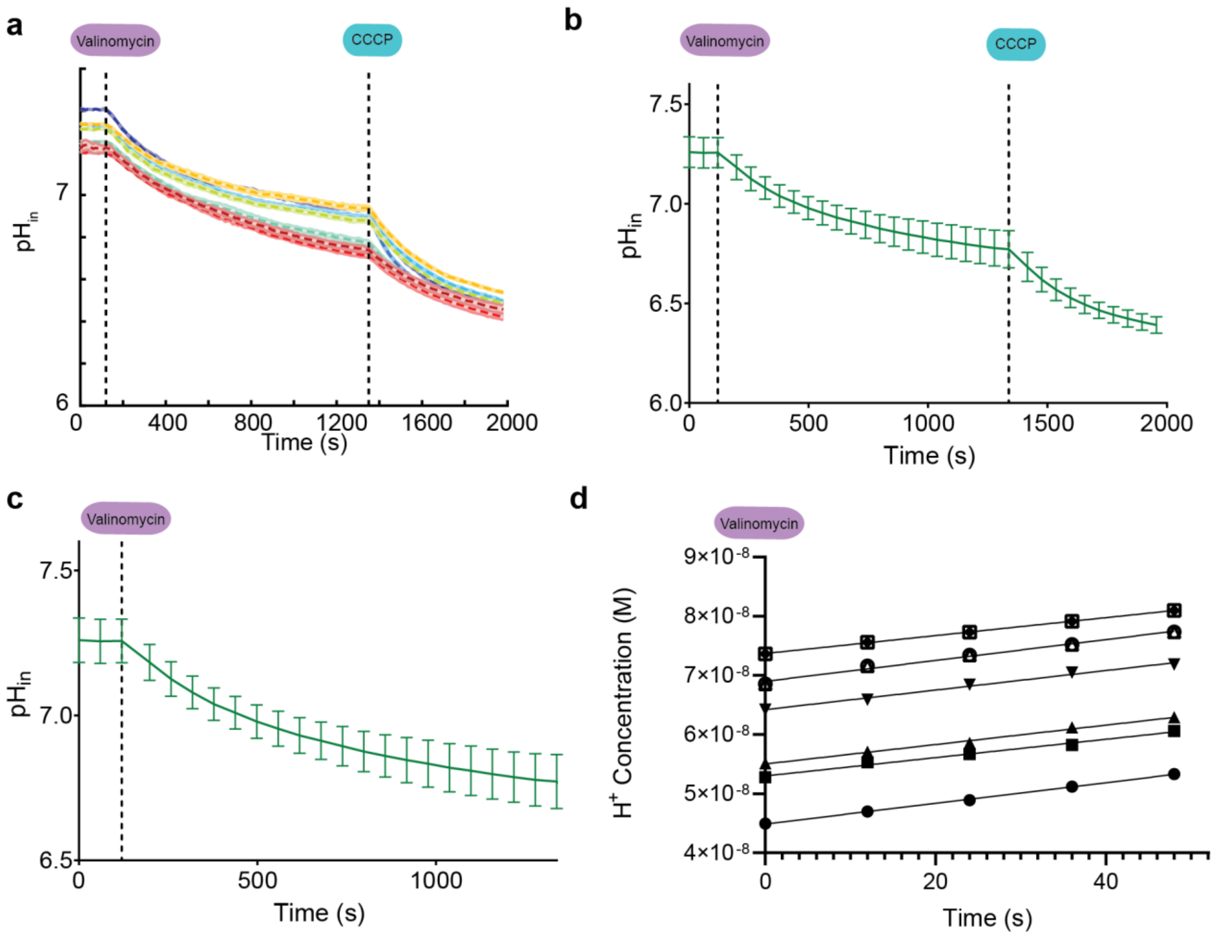


**Extended Data Fig. 6. All proton flux assay data for LLQL vesicle samples.** Eight samples (each run in triplicate with shaded error bars shown) containing 1:500 peptide:lipid ratio; samples were run independently in the assay. **a,** $pH_{in}$ as a function of time throughout the measurement for each independent sample. **b,** Mean and standard deviation for data collection. **c,** Data prior to CCCP addition shows significant change in $pH_{in}$ after addition of valinomycin. **d,** Fits for the initial 50 seconds following addition of valinomycin. From the linear regression fits, all slopes (which give the initial rates (in M/s)) were used to calculate mean and standard error presented in Fig. 6g and Supplementary Table S3.

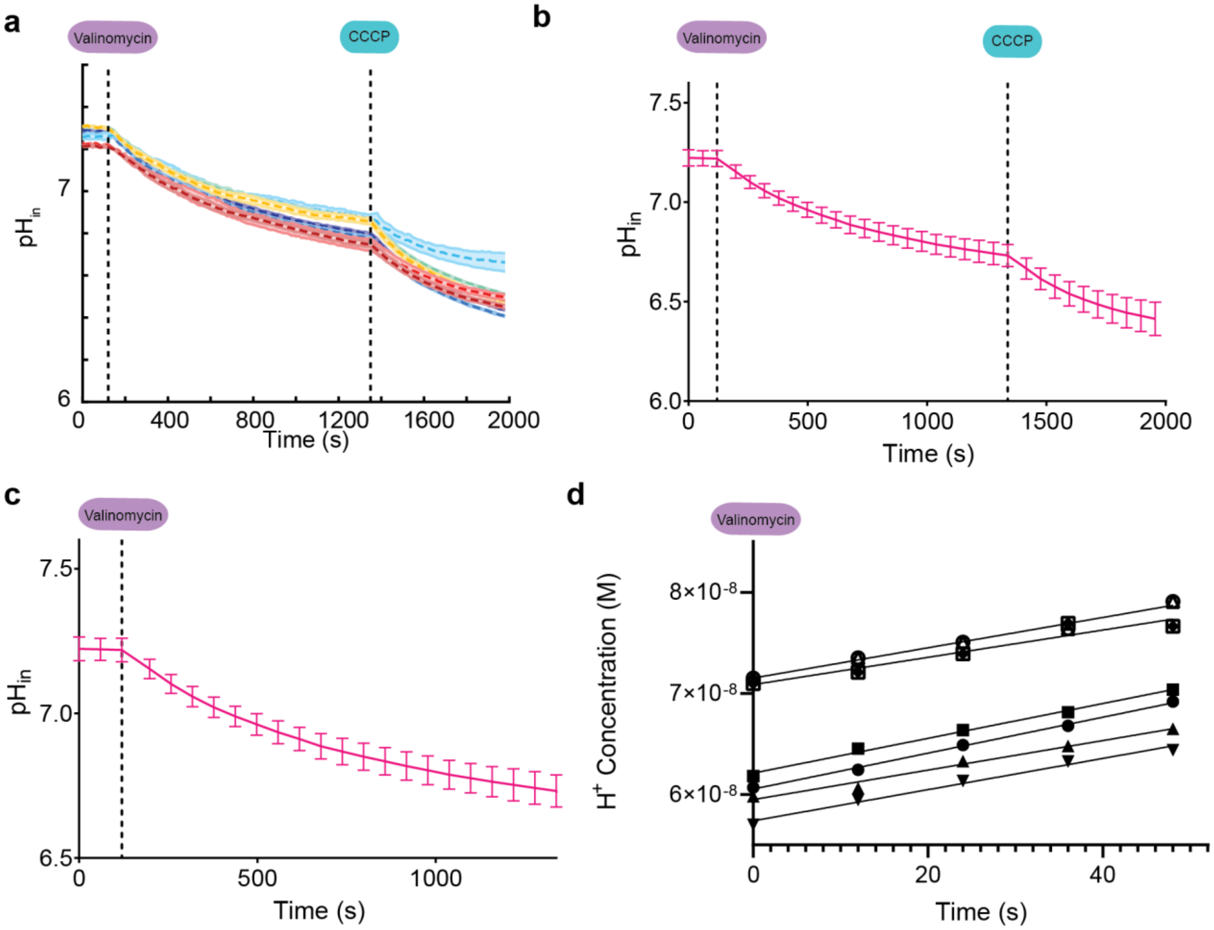


**Extended Data Fig. 7. All proton flux assay data for QLQL vesicle samples.** Eight samples (each run in triplicate with shaded error bars shown) containing 1:500 peptide:lipid ratio; samples were run independently in the assay. **a,** $pH_{in}$ as a function of time throughout the measurement for each independent sample. **b,** Mean and standard deviation for data collection. **c,** Data prior to CCCP addition shows significant change in $pH_{in}$ after addition of valinomycin. **d,** Fits for the initial 50 seconds following addition of valinomycin. From the linear regression fits, all slopes (which give the initial rates (in M/s)) were used to calculate mean and standard error presented in Fig. 6g and Supplementary Table S3.

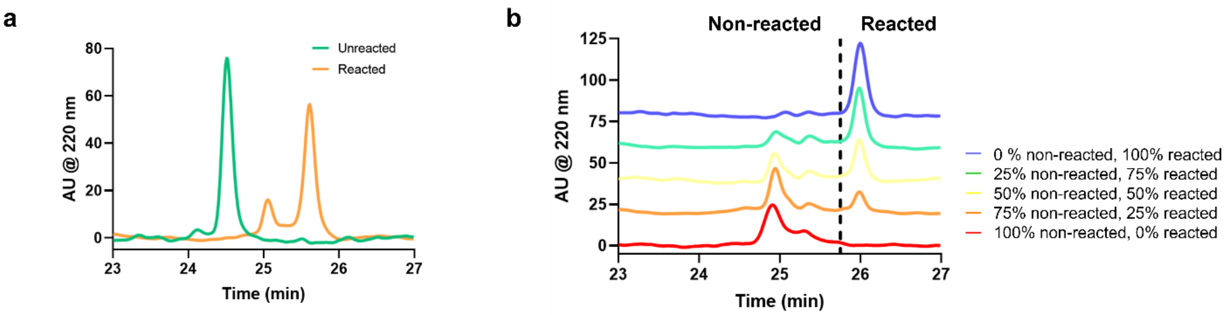


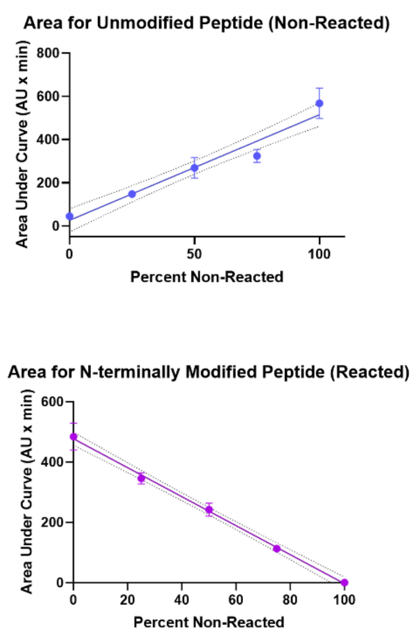


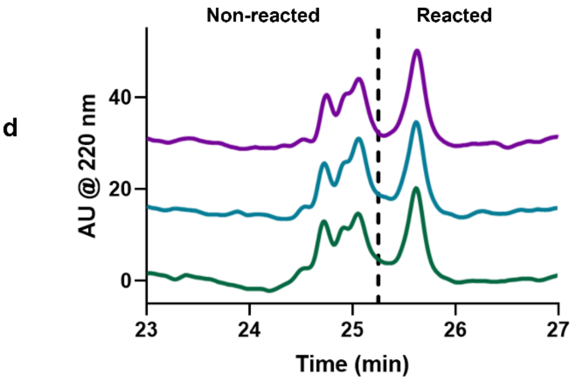


**e**

| Sample | Area Under Curve Non-Reacted (NR) | % NR Protein | Area Under Curve Reacted (R) | % R Protein |
|---|---|---|---|---|
| 1 | 204.351 | 56.8 | 256.032 | 47.1 |
| 2 | 199.730 | 57.8 | 276.955 | 51.4 |
| 3 | 229.975 | 51.5 | 271.431 | 50.3 |

**Extended Data Fig. 8. Determining orientation of pentamers in vesicles. a,** HPLC trace of unreacted and reacted peptides following reaction with the highly polar, amine-reactive methyltetrazine 3-sulfo-N-hydroxysuccinimide ester (methyltetrazine sulfo-NHS, see Materials and Methods). The only amine-reactive groups are the N-terminus or the N-terminal lysine sidechain. Thus, only peptides in which N-terminus is exposed on the outside of the vesicle should react to the dye. **b,** HPLC traces of mixtures of different ratios of non-reacted and reacted peptides. **c,** Calibration curves for area under the curve for nonreacted and reacted peaks in the HPLC traces corresponding to the different mixtures in **b. d,** Traces of three independent samples of LQLL pentamers from vesicles after reaction with methyltetrazine sulfo-NHS. **e,** Using the calibration curves in **c**, the area under the curve was determined for each sample. The data indicate that half the amines react, as expected from a random orientation of pentamers in the lipid vesicle.
