## Supplementary Materials for "Transient Water Wires Mediate Selective Proton Transport in Designed Channel Proteins"

### Materials and Methods

Synthesis of Proton Channel Peptides

Proton channel peptides, shown in Fig. 2 and Supplementary Table S1, were synthesized using Fmoc-based solid-phase peptide synthesis on either a Biotage Initiator Alstra microwave synthesizer or a Syro II parallel peptide synthesizer. All peptides were synthesized with a free amine N-terminus and a C-terminal carboxamide using Tentagel S-RAM resin (Chem-Impex Int'l Inc) with 0.22-0.24 mmol/g. Following synthesis, the peptides were cleaved from the resin with a trifluoroacetic acid: triisopropylsilane: water (TFA:TIPS:$H_2O$, 95:2.5:2.5) solution, precipitated out with cold diethyl ether, redissolved in 50% 1,1,1,3,3,3-hexafluoroisopropanol (HFIP) and water, and purified using reverse phase HPLC on a C4 prep column (Vydac) with a gradient of solvents A (water, 0.1% TFA) and B (isopropanol:acetonitrile:water:TFA, 60:30:9.9:0.1) at a flow rate of 10 mL/min. Peptides were then lyophilized until dry. Following successful purification, peptides were confirmed to be >90% pure with analytical HPLC over a C4 column (Phenomenex) and confirmed to have the correct mass with MALDI mass spectrometry (Shimadzu AXIMA Performance) using α-cyano-4-hydroxycinnamic acid (Sigma) as the matrix (Supplementary Table S1). Peptides were then dissolved in ethanol to the appropriate stock concentrations and subsequently used for all experiments as described.

Expression of Full-Length Influenza A M2 Polypeptide

The Influenza A M2 polypeptide from previous work [49] was expressed and purified from *E. coli* cells. Briefly, the gene containing the A/Udorn/72 Cys-free W15F variant (W15F, C17S, C19S, C50S) and a C-terminal 6x His tag was cloned into a pEXP-5-NT plasmid and transformed into chemically competent *E. coli* BL21(DE3) cells (Invitrogen) using heat shock methods. The cells were grown in TB media (Invitrogen) with ampicillin at 37°C. When an $OD_{600}$ of 0.6-0.8 was achieved, the expression culture was induced with 1 mM isopropyl ß-D-1-thiogalactopyranoside (IPTG). The cells were then harvested no more than 2.5 hours after induction and centrifuged down at 4°C for 10 minutes. Pellets were then frozen prior to purification. For purification, the cells were resuspended in lysis buffer (50 mM Tris pH 8, 300 mM NaCl, 5% glycerol, 25 mM imidazole, 0.1% decyl maltose neopentyl glycol (DMNG), 8 M urea) and sonicated with a microtip sonicator for 5 minutes at 20% amplitude (2 s on/off). Lysed cells were then spun down for 15 minutes at 18000 rpm and purified using Ni-NTA beads using batch purification methods. Wash buffer was the same as the lysis buffer. The protein was eluted in elution buffer (50 mM Tris pH 7.5, 150 mM NaCl, 5% glycerol, 250 mM imidazole, 0.1% DMNG, 8 M urea) and mixed with HFIP prior to loading onto the C4 column for secondary purification by HPLC. Polypeptide was lyophilized until dry and reconstituted in ethanol to appropriate stock concentrations. SDS-PAGE and MALDI-MS were used to confirm the expression of the polypeptide.

<u>SDS-PAGE Gels of Proton Channels</u>

25-50 µg of peptide were lyophilized from the peptide stocks and dissolved in 25 mM Tris pH 7.5 (Fisher Scientific) with 50 mM octyl-β-glucopyranoside (OG, Carbosynth Limited) for a final concentration of 2.5 µg/µL. An equal volume of 2x LDS (Invitrogen) was added to each sample and the sample was subsequently boiled for 15 minutes at 95ºC. 5 µg of peptide was loaded in each well of a 12% Bis-Tris NuPAGE gel (Thermo Fisher Scientific) with Precision Plus Protein Dual Xtra prestained protein standards (Bio-Rad Laboratories); the gel was subsequently run for 30 minutes at 200 V.

<u>Proton Channel Liposomal Flux Assays and Analysis</u>

The proton channel liposomal flux assays were adapted from references [48,49], and the exact methodologies for preparation and running of these assays are described below.

*Preparation of Proteoliposomes*

All proteoliposome samples were made as a stock solution of 1:500 peptide:lipids with final concentrations of 10 µM and 5 mM, respectively. To prepare the proteoliposome stock, 10 uL of 100 mM of a ratiometric pH-sensitive dye, 8-hydroxypyrene-1,3,6-trisulfonic acid trisodium salt (HPTS) was added to total of 2 mL of $K^+$ buffer (50 mM $K_2SO_4$, 30 mM $K_2HPO_4$, pH 7.5). The solution was then added to a dried film of 3:1 1-

palmitoyl-2-oleoyl-sn-glycero-3-phosphocholine:1-palmitoyl-2-oleoyl-sn-glycero-3-phosphoglycerol (POPC:POPG, Avanti Lipids) to make a 10 mM lipid solution, which was subsequently vortexed for 1 minute before sonicating with microtip for 5 min at 20% amplitude for 2 s on/off. The sonicated solution was then divided into 200 uL aliquots where 1M OG was added to a final concentration of 26 mM OG. After one hour on the rotisserie at RT, peptide in 26 mM OG in $K^+$ buffer (or, in the case of empty, 15 µL of 26 mM OG in $K^+$ buffer) was added to the solution of detergent-solubilized lipids for a final concentration of 20 µM peptide. The peptide-detergent-lipid (PDL) solution was equilibrated for 1 hour at RT on the rotisserie. Following the incubation of the PDL solution, 50 µL of XAD biobeads solution (Sigma-Aldrich) in $K^+$ buffer was added to the PDL mixture every 5-10 minutes five times for a final volume of around 500 µL and final peptide and lipid concentrations as described above. After the final addition of biobeads, the solution was left to rotate in the 4˚C rotisserie overnight. In the following morning, the sample was spun down in an ultracentrifuge at 96k rpm for 10 minutes to pellet the liposomes. The liposome pellet was resuspended in the same amount of dye-free $K^+$ buffer to afford the stock solution used for the liposomal assays. Dynamic light scattering (Malvern Panalytical) of the vesicle samples enabled size determination prior to proton flux assays to ensure liposome formation.

*Preparation of Working Solutions for Proton Flux Assays*

A working 3 µM valinomycin solution was made by dissolving 1 mg/mL (~900 mM) valinomycin stock in DMSO (purchased from Sigma-Aldrich) into $Na^+$ buffer (50 mM $Na_2SO_4$, 30 mM $Na_2HPO_4$, pH 7.5). Similarly, a 3 µM CCCP working solution was generated by dissolving 600 µM CCCP stock in DMSO into $Na^+$ buffer.

For the assay, 17.3 µL of the stock proteoliposome solution was diluted into 632.7 µL of $Na^+$ buffer mixed with 10 mM p-xylene-bis-pyridinium bromide (DPX, Invitrogen) for a total of 650 uL of sample at the working concentrations. The presence of membrane-impermeable DPX quenches any extraliposomal HPTS fluorescence. Therefore, the fluorescence measured in this experiment comes only from intraliposomal HPTS. The sample was allowed to equilibrate for 20 minutes before 190 µL was aliquoted into three wells on a black, u-shaped bottom 96-well plate (Greiner) for fluorescence measurements (collected every 55 s).

*Instrumentation*

All data reported in Fig. 6, Extended Figs. 3-5, Supplementary Figs. S19-S23 were collected using the Biotek Synergy 2 equipped with a 405 nm/20 nm bandwidth and 460 nm/40 nm bandwidth excitation filters and a 528 nm/20 nm bandwidth emission filter (Biotek). The valinomycin and CCCP solutions (10 uL each) were added to the wells using an injector system at 120 s and 1350 s into the assay, respectively, with 3 s of vigorous shaking following addition. The fluorescence was measured at the two excitation wavelengths every 12 s.

For the long-time kinetics run (Extended Data Fig. 2), the data was collected on a Biotek Synergy H1 using monochromators. The excitation wavelengths were set at 417 nm and 460 nm, and the emission wavelength was set to 515 nm.

*Calibration and Analysis of Flux Data*

For all data reported in Figs. 6, Extended Data Figs. 3-5, and Supplemental Figs. S19-S23, the fluorescence signals were calibrated using the ratio of the fluorescence signals of the deprotonated dye, $F^-$ ($\lambda_{excitation}$ = 460 nm, $\lambda_{emission}$ = 528 nm) and the isosbestic point, $F_{iso}$ ($\lambda_{excitation}$ = 405 nm, $\lambda_{emission}$ = 528 nm) as a function of pH from pH 8 to pH 4 (Extended Data Fig. 1). Similarly, for the long-time kinetics runs, a calibration curve of the ratio of the fluorescence signals of $F^-$ ($\lambda_{excitation}$ = 460 nm, $\lambda_{emission}$ = 515 nm) and $F_{iso}$ ($\lambda_{excitation}$ = 417 nm, $\lambda_{emission}$ = 515 nm) was derived as a function of pH (Extended Data Fig. 1). All fluorescence measurements for each reported sample was taken in triplicate and were then converted into values of $pH_{in}$ using the defined calibration curves.

For data reported in Fig. 6, Extended Data Fig. 3-5, and Supplemental Figs. S19-S23, initial rates of proton conduction were determined for each sample using data collected within the first sixty seconds following the addition of valinomycin. Similarly, for the long-time kinetics runs (Extended Data Fig. 2), initial rates were determined by fitting the first 220 seconds of data following the addition of valinomycin. All data were fit using linear regression and the initial rates were extracted from the fitted slope.

Orientation Determination of Peptides in Vesicles

Liposomes containing 108 uM of peptide (P:L ratio of 1:50) were used for the following experiment. Calibration curves were prepared as follows: 75 uL of liposome solution was added to 37.5 uL of 1 M OG and 37.5 uL of water (for non-reacted or NR samples) or 37.5 uL of 20 mM methyltetrazine-sulfo NHS ester (Click Chemistry Tools) in water (for reacted or R samples) for 30 minutes. Samples were then quenched with 16.67 uL of 1 M Tris pH 8 for 5 minutes. Non-reacted and reacted samples were mixed in differing ratios and diluted in 50% HFIP in water before running on reverse phase HPLC on an analytical C4 column with a gradient of solvents A (water, 0.1% TFA) and B (isopropanol:acetonitrile:water:TFA, 60:30:9.9:0.1) at a flow rate of 1 mL/min. The area under the curves for the non-reacted and reacted peaks were calculated for each sample to generate the calibration curve (Extended Data Fig. 8). Traces in figures are background-subtracted and offset for ease of viewing.

To determine orientation of the samples, 50 uL of the liposome sample was mixed with 50 uL of 10 mM methyltetrazine sulfo-NHS ester in water for 30 minutes and subsequently quenched with 5.56 uL of 1 M Tris pH 8 for 5 minutes. The samples were then diluted with 50:50 1 M OG and HFIP before running on HPLC. The area under the curve was calculated for the non-reacted and reacted peaks and used with the calibration curves to estimate the orientation of the peptide (Extended Data Fig. 8d,e). All samples were measured in duplicates.

Lipidic Cubic Phase (LCP) Crystallography of Proton Channels

The methods described for LCP crystallization are originally described in detail in Caffrey and Cherezov and have been slightly modified as follows [72,73]. Peptide from the ethanol stock was mixed with 60 mg of monoolein (Sigma-Aldrich) until clear. The solution containing peptide and monoolein was then dried under a gentle stream of $N_2$ and lyophilized overnight. To prepare the LCP, the monoolein-peptide mixture was heated to 42ºC until it became liquid, and subsequently extruded ~200 times with 2/3 times the volume of 50 mM OG in coupled gastight Hamilton syringes at room temperature. Successful LCP formation was confirmed as the solution became clear and was not birefringent in the cross-polarizer. The final concentration of peptide for each sample is listed in Supplementary Table S2.

For crystallization, 50 nL of the LCP mixture was dispensed onto 96-well Laminex plastic sandwich plates (Molecular Dimensions) with 0.5 to 1 uL of precipitant solution using the TTP Labtech LCP Mosquito robot at room temperature. Plates were sealed using plastic coverslips (Molecular Dimensions) and monitored using a Formulatrix RockImager at 20ºC. Crystals of each construct were harvested from conditions noted in Supplementary Table S2 with and cryoprotected with 30% v/v PEG 400, if necessary, and flash-frozen with liquid nitrogen.

X-Ray Diffraction Data Collection and Analysis

Crystals were mounted under a cryostream at 100K and data was collected at both the 8.3.1 beamline at the Advanced Light Source at Lawrence Berkeley National Lab at a wavelength of 1.115832 Å or at 23-ID-B/D at Argonne National Laboratory at a wavelength of 1.03318 Å. The data was processed with the XDS package [74] and reduced with AIMLESS within the CCP4 suite [75]. The structures were determined by molecular replacement with Phaser [76] using a previously designed de novo protein (pdb code: 6mct) as a model. The models were rebuilt in Coot [77] and the structures were then refined with PHENIX [78]. The data processing and structural refinement statistics are described in Extended Data Table 2.

Solid-state NMR (ssNMR) Experiments

LQLL and LLLL peptides with the appropriate site-specific $^{13}C$, $^{15}N$ labels at the Ile6, Ile13 or Gln10 positions were made using SPPS methods as described previously. The purified peptides were reconstituted into d54-DMPC lipids at a protein/lipid molar ratio of 1:12 with a final peptide weight of ~5 mg. The samples are concentrated to a hydration level of ~40% (w/w) and pelleted into 3.2 mm rotors for NMR experiments.

Magic-angle-spinning (MAS) solid-state nuclear magnetic resonance (NMR) spectra were measured on a Bruker Avance III HD 600 MHz (14.1 T) spectrometer with a Bruker 3.2

mm HXY MAS probe operating in double-resonance $^{1}H/^{13}C$ mode. All $^{13}C$ spectra were externally referenced to to the adamantane $CH_2$ resonance at 38.48 ppm on the trimethylsilane scale. All spectra were recorded under 10.5 kHz MAS at a sample temperature of 277 K. The sample temperature was estimated based on the $^{1}H$ chemical shift of bulk water at 4.97 ppm, according to the equation T (K) = 96.9 × (7.83 ppm – $\delta_{H2O}$) [79]. Typical radiofrequency field strengths were 50-80 kHz for $^{1}H$ and 50-60 kHz for $^{13}C$. All spectra used a recycle delay of 3 s.

Water-edited $^{13}C$ cross-polarization (CP) experiments were used to probe the water accessibility and hydration of the channel[80-84]. These consisted of a $^{1}H$ excitation pulse at 71.4 kHz, followed by a 52 rotor period (5.0 ms) Hahn echo with a selective Gaussian 180˚ pulse of 4.8 ms placed on resonance with water. This water-selective echo period is followed by a $^{1}H$ spin diffusion period, whose duration ($t_{SD}$) was varied from 1 to 225 ms. The $^{1}H$ spin diffusion period was followed by a 500 μs $^{1}H$-$^{13}C$ cross-polarization (CP) for $^{13}C$ detection, during which $^{1}H$ two-phase modulation (TPPM) decoupling was applied at 71.4 kHz.

Water-edited $^{13}C$ spectral intensities were analyzed as integrated intensities from 0-75 ppm for all samples and as peak heights for the resolved Ile13 signals in LQLL and LLLL samples. These intensities are corrected for water $^{1}H$ $T_1$ relaxation by dividing each value by $\exp(-t_{SD}/T_1)$. The $T_1$ corrected intensities are then normalized to the value at 225 ms. The water $^{1}H$ $T_1$ values were measured using the inversion-recovery experiment and ranged from 1.0–1.5 s.

Classical Molecular Dynamics Simulations

The model for the engineered pentameric membrane protein LLLL was taken from its crystal structure (pdb: 6mct). Molecular models of the Leu-to-Gln (LQ) variant proteins were built and sidechains repacked to the global energetic minima using Scrwl4 [85], with the LLLL crystal structure as the input template. The simulations were performed prior to LCP X-ray structures being solved, and thus modeled *a priori*. The initial transmembrane orientation of LLLL in a lipid bilayer was predicted by the OPM PPM 2.0 server [86], and the LQ variant proteins were modeled with identical geometry as LLLL relative to the membrane by structural superposition.

The MD system was built through an automatic script merging protein and membrane components using VMD [87] and the GROMACS engine [88]. First, a pre-assembled 7.5 x 7.5 nm 1-palmitoyl-2-oleoyl-sn-glycero-3-phosphocholine (POPC) bilayer was modeled using VMD's membrane builder application and aligned with the implicit bilayer used to predict the membrane proteins' insertion geometry. The oriented membrane protein was merged with the lipid bilayer and lipid molecules which clashed with protein atoms (<1.2 Å overlap) were removed. The system was treated as a periodic box, 7.0 nm in the Z direction, and hydrated with TIP3P water (ca. 20 Å of water regions above and below the bilayer). KCl was added to the system to neutralize protein charge and to yield a final 0.15 M ion concentration. The system used CHARMM36 parameters [89] and the GROMACS 2018 engine was used for minimization and dynamics simulations. The recommended CHARMM36 cut-offs (rcoulomb, rvdw = 1.2 nm), switching (1.0 nm), and Particle-Mesh Ewald distances were used. A 2 fs time step was used for Langevin dynamics.

The system was minimized with the steepest decent algorithm (5000 steps max, tolerance of < 1000.0 kJ/mol/nm) without atomic position restraints. Harmonic positional restraints of 1 kcal $mol^{-1}$ $Å^{-2}$ on all non-hydrogen protein atoms. A 50 ps NVT dynamics simulation was initiated from the minimized model using harmonic position restraints on all non-hydrogen protein atoms (1 kcal $mol^{-1}$ $Å^{-2}$) using the Velocity-rescale thermostat fixed at 298.15° K with a 0.1 ps coupling constant. Next, a 15 ns restraint NPT equilibration simulation was run using a semiisotropic Berendsen barostat (P = 1 bar, pressure coupling time constant = 5 ps, compressibility = 4.5E-5 bar-1) and a Berendsen thermostat (T = 298.15° K, 1.0 ps time constant) while maintaining 1 kcal mol-1 Å-2 harmonic restraints on protein Cα atoms relative to the input structure. Unrestrained production dynamics simulations were then run for 200 ns with a Nose-Hoover thermostat at 298.15° K with a 1.0 ps time constant and a semiisotropic Parrinello-Rahman barostat fixed at 1 bar with a pressure coupling constant of 5 ps. Coordinate frames were extracted at 20 ps intervals in these production simulation trajectories. Three independent simulation trajectories were launched for each unique protein sequence, using different initial atomic velocities.

The backbone atoms from every frame in the trajectory versus the initial frame had an atomic RMSD in the range of 0.6-1.2 Å for all variant studies, ansd each triplicate trajectory. Likewise, across all variants, the average RMSD between any two frames throughout the MD trajectory was <1 Å. By these backbone RMSD metrics we can determine the protein fold and tertiary structure is very stable for all variants and deviates from the LLLL model on the same order as thermal fluctuations. The backbone RMSD of the medoid frame of each triplicate trajectory versus the LCP X-ray structures for the LQ variants were all <1.2 Å.

Analysis of Classical MD Simulations

Analysis of the channels and their water content was done using the Channel Annotation Package (CHAP, https://www.channotation.org) available from the Sansom Lab [44]. The trajectories from the production runs were centered to the protein using GROMACS software and sampled every 200 ps to create a compressed xtc trajectory file for CHAP analysis. Water density plots (like those seen in Figs. 3 and 6 and Supplementary Figs. S1-S6) were generated for each 3 x 200 ns trajectory for each of the pentameric bundle designs. Similarly, the time-averaged water density profiles (i.e. panel c of Supplemental Figs. S1-S6) were generated by calculating the average water density for a given s over the course of the 200 ns simulation. These time-averaged water density profiles were then used to calculate the observed hydrophobic lengths ($l_{obs}$) reported in Fig. 2c and Extended Data Table 1 using a second derivative method.

Multiscale Reactive MD Simulations

The X-ray crystal structures for LQLL (pdb 7udz) and LLLL (pdb 6mct) were used as the starting structures for simulation. Classical simulations were first performed to equilibrate the protein structures and the simulation systems. Each protein was embedded in a 1-palmitoyl-2-oleoyl-sn-glycero-3-phosphocholine (POPC) bilayer and solvated with water using the CHARMM GUI [90-93], and the membrane and water were equilibrated using a standard equilibration protocol. Classical equilibration with no restraints was performed for 500ns. The CHARMM36 forcefield [94] was used to model all interactions, and simulations were run at 298K in the NPT ensemble using GROMACS [95].

MS-RMD [45-47,96] was subsequently used to model the water and excess proton in all simulations used in our analyses. The MS-RMD method captures proton delocalization in water by allowing hydrogen-oxygen bonds to break and form. This is done by taking a linear combination of possible bonding topology states at every timestep. See our previous work for a detailed description and theory. The MS-EVB 3.2 parameters were used to describe the hydrated excess proton [97]. The excess proton center of excess charge (CEC) is defined as [98]:

$$\vec{r}_{CEC} = \sum_{i}^{N} c_i^2 \, \vec{r}_{COC}^{\,i} \,, \qquad (1)$$

Where $\vec{r}_{COC}^{\,i}$ is the coordinate of the center of excess charge of the ith diabatic state and $c_i^2$ is the amplitude of that state. The CEC defines the position of the delocalized excess proton. The CEC defines the position of the delocalized excess proton. The CHARMM36 forcefield was used to model the remaining interactions. Simulations were run at 298K in the NVT ensemble using LAMMPS [99] with the MS-RMD package.

Umbrella sampling simulations were performed in two dimensions to model the PT process. The first collective variable (CV) used is the CEC position along the channel axis, Z'CEC. The channel axis for each system was defined as the average principal component of the protein from a 750ps MS-RMD simulation after equilibration. The position along this axis is calculated in reference to the center of mass of the Ile13 alpha-carbons, such that Z'CEC = 0Å at that point. The second CV used, φ, is a recently developed CV that measures the water connectivity within a channel using graph theory [8]. This CV is a significant improvement over water density, which does not directly bias the formation of a continuous water wire and can result in unphysical water “clumps” as the bias increases. Instead, the new water connectivity CV measures the length of transient water wire formations on a scale from 0 to 1, where 0 is no water and 1 is water fully connected throughout the channel, agnostic to the number of water molecules. We refer the reader to reference [8] for further theoretical details.

The open-source, community developed PLUMED library [100,101] was used to define the umbrella bias potentials. Umbrella sampling windows were set up every 0.5Å in the range [-22, 22] Å for Z'CEC and every 0.035 in the range [0.140,0.980] and [0.245,0.980] for φ for LLLL and LQLL, respectively, for a total of ~2000 windows for each system. Initial windows were generated by using steered MD to pull water into the channel, and the excess proton was placed at each point along the channel. Subsequent windows were pulled from nearby windows. Windows were equilibrated for 100ps, and then run for 0.5-3.5ns. A few additional windows were added to ensure sampling overlap.

The 2D PMFs were calculated using WHAM-2D [102]. Error bars were calculated using the block method with four blocks. The minimum free energy path (MFEP) was calculated using the string method.

Figs. 5, Supplemental Figs. S17, S18, and S24 were generated with Matplotlib [103]. Fig. 5 was generated using Visual Molecular Dynamics (VMD) [87].

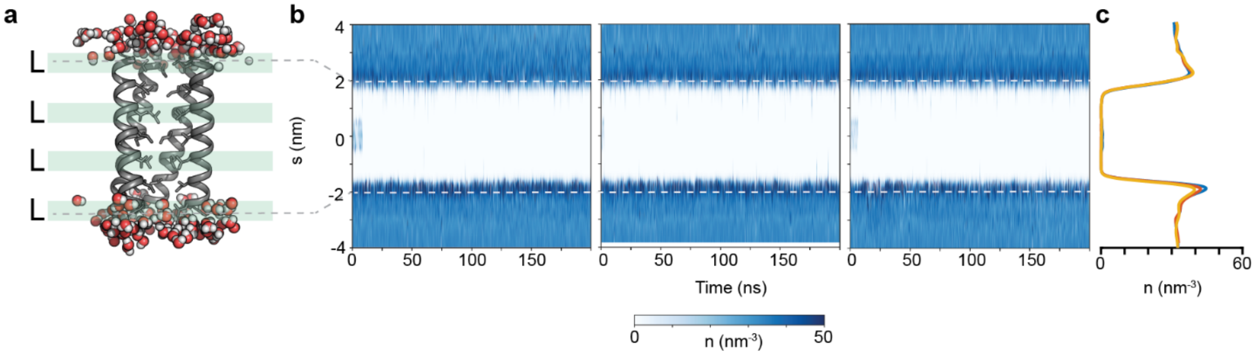

**Supplementary Fig. S1. Analysis of classical MD simulations of LLLL. a,** Snapshot taken from production run #1 of 3 of LLLL simulations. Fifth helix removed for clarity. **b,** Water density traces for all 3 x 200 ns trajectories show the channel (which spans s = -2 to 2 nm) devoid of water over the course of the simulation. **c,** Time-averaged water density as a function of s corresponding to the three independent simulations (blue, red, and yellow are productions #1, #2, and #3, respectively). From the second derivative of the trace, the $l_{obs}$ of the channel was calculated to be 33.9 ± 0.3 Å. The leftmost water density plot is shown in Figs. 3a and 6a of the main text.

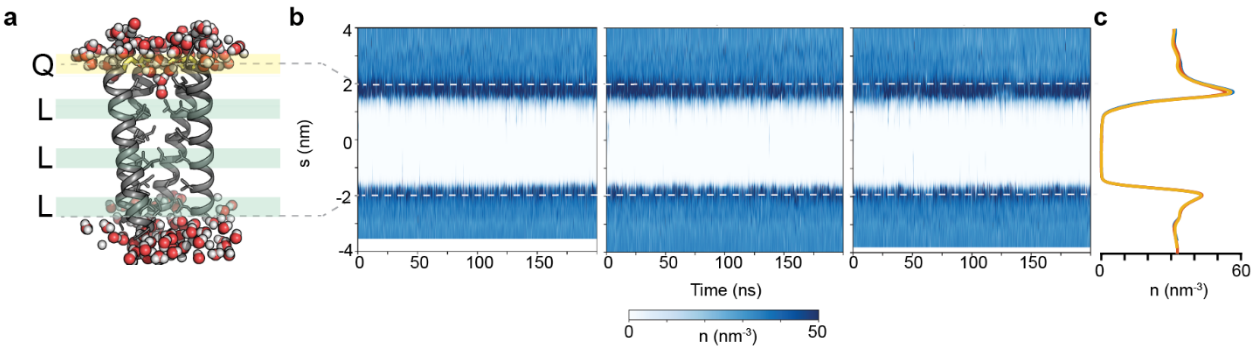

**Supplementary Fig. S2. Analysis of classical MD simulations of QLLL. a,** Snapshot taken from production run #1 of 3 of QLLL simulations. Fifth helix removed for clarity. **b,** Water density traces for all 3 x 200 ns trajectories show the channel (which spans s = -2 to 2 nm), like LLLL, devoid of water over the course of the simulation. Notably, there is an increase in water density near the Gln site, at s = +2 nm. **c,** Time-averaged water density as a function of s corresponding to the three independent simulations (blue, red, and yellow are productions #1, #2, and #3, respectively). From the second derivative of the trace, the $l_{obs}$ of the channel was calculated to be 32.0 ± 0.6 Å. The leftmost water density plot is shown in Fig. 3b.

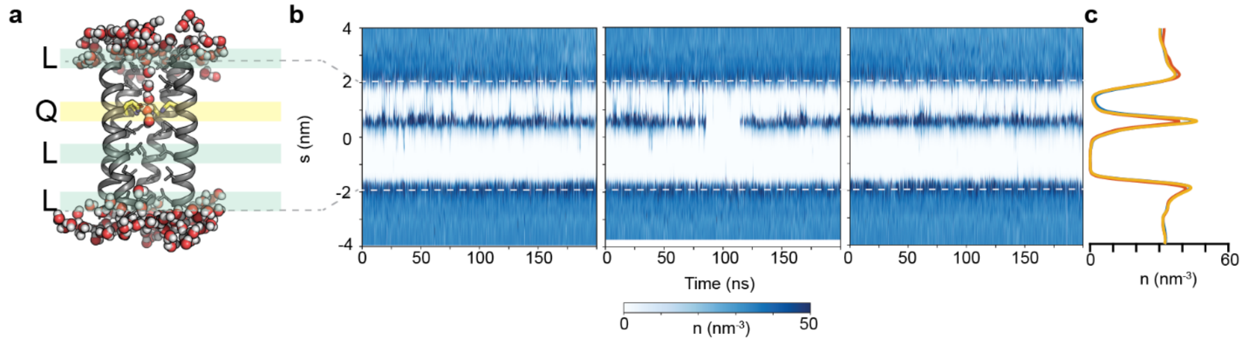

**Supplementary Fig. S3. Analysis of classical MD simulations of LQLL. a,** Snapshot taken from production run #2 of 3 of LQLL simulations. Fifth helix removed for clarity. **b,** Water density traces for all 3 x 200 ns trajectories show the channel (which spans s = -2 to 2 nm) with increased water densities within the pore near the Gln site, s = +0.5 nm. Moreover, there seems to be events in which multiple waters span the shorter segment. **c,** Time-averaged water density as a function of s corresponding to the three independent simulations (blue, red, and yellow are productions #1, #2, and #3, respectively). From the second derivative of the trace, the $l_{obs}$ of the channel was calculated to be 20.7 ± 0.1 Å. The shorter hydrophobic path at the N-terminal end of the channel was calculated to be 11.6 ± 0.4 Å. The middle water density plot is shown in Figs. 3c and 6e.

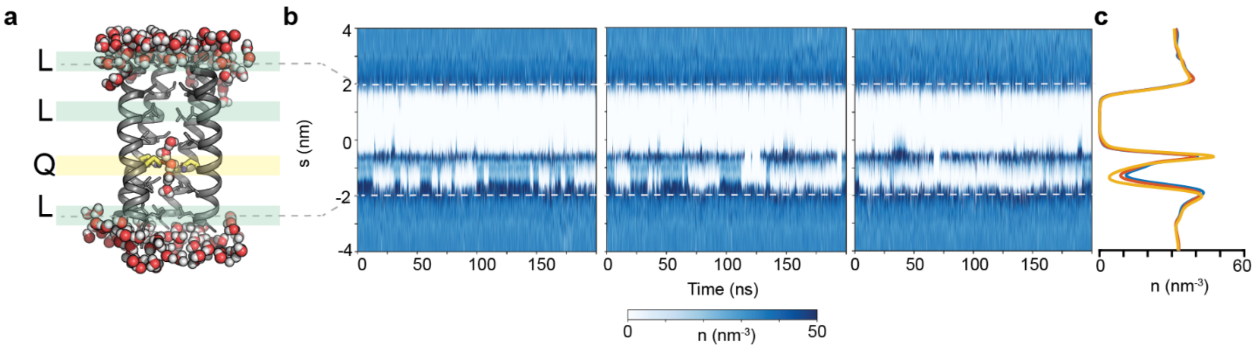

**Supplementary Fig. S4. Analysis of classical MD simulations of LLQL. a,** Snapshot taken from production run #1 of 3 of LLQL simulations. Fifth helix removed for clarity. **b,** Water density traces for all 3 x 200 ns trajectories show the channel (which spans s = -2 to 2 nm) with increased water densities within the pore near the Gln site, s = -0.5 nm. Moreover, there seems to be events in which multiple waters span the shorter segment. **c,** Time-averaged water density as a function of s corresponding to the three independent simulations (blue, red, and yellow are productions #1, #2, and #3, respectively). From the second derivative of the trace, the $l_{obs}$ of the channel was calculated to be 22.4 ± 0.1 Å. The shorter hydrophobic path at the C-terminal end of the channel was calculated to be 9.3 ± 0.5 Å. The rightmost water density plot is shown in Fig. 3d.

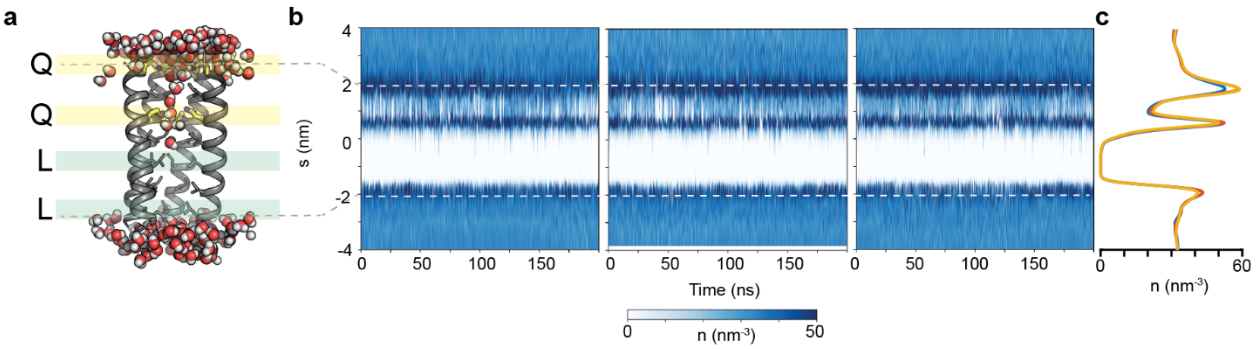

**Supplementary Fig. S5. Analysis of classical MD simulations of QQLL. a,** Snapshot taken from production run #1 of 3 of QQLL simulations. Fifth helix removed for clarity. **b,** Water density traces for all 3 x 200 ns trajectories show the channel (which spans s = -2 to 2 nm) with increased water densities within the pore near the Gln sites, at s = +2 and +0.5 nm. Moreover, there seems to be events in which multiple waters span the shorter segment. **c,** Time-averaged water density as a function of s corresponding to the three independent simulations (blue, red, and yellow are productions #1, #2, and #3, respectively). From the second derivative of the trace, the $l_{obs}$ of the channel was calculated to be 20.9 ± 0.1 Å. The shorter hydrophobic path at the C-terminal end of the channel was calculated to be 8.2 ± 0.2 Å. The leftmost water density plot is shown in Figs. 3e and 6f.

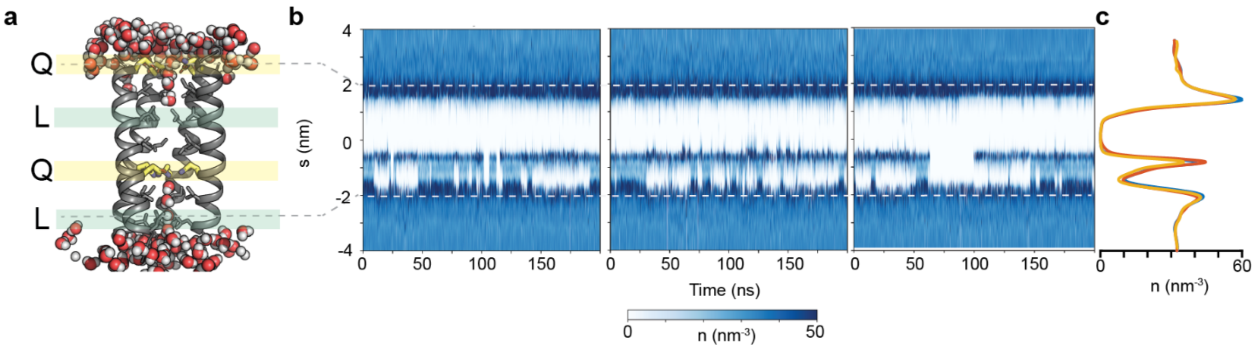

**Supplementary Fig. S6. Analysis of classical MD simulations of QLQL. a,** Snapshot taken from production run #1 of 3 of QLQL simulations. Fifth helix removed for clarity. **b,** Water density traces for all 3 x 200 ns trajectories show the channel (which spans s = -2 to 2 nm) with increased water densities within the pore near the Gln sites, at s = +2 and -0.5 nm. Moreover, there seems to be events in which multiple waters span the shorter segment. **c,** Time-averaged water density as a function of s corresponding to the three independent simulations (blue, red, and yellow are productions #1, #2, and #3, respectively). From the second derivative of the trace, the $l_{obs}$ of the channel was calculated to be 20.3 ± 0.2 Å. The shorter hydrophobic path at the C-terminal end of the channel was calculated to be 9.2 ± 0.2 Å. The rightmost water density plot is shown in Fig. 3f.

a
b

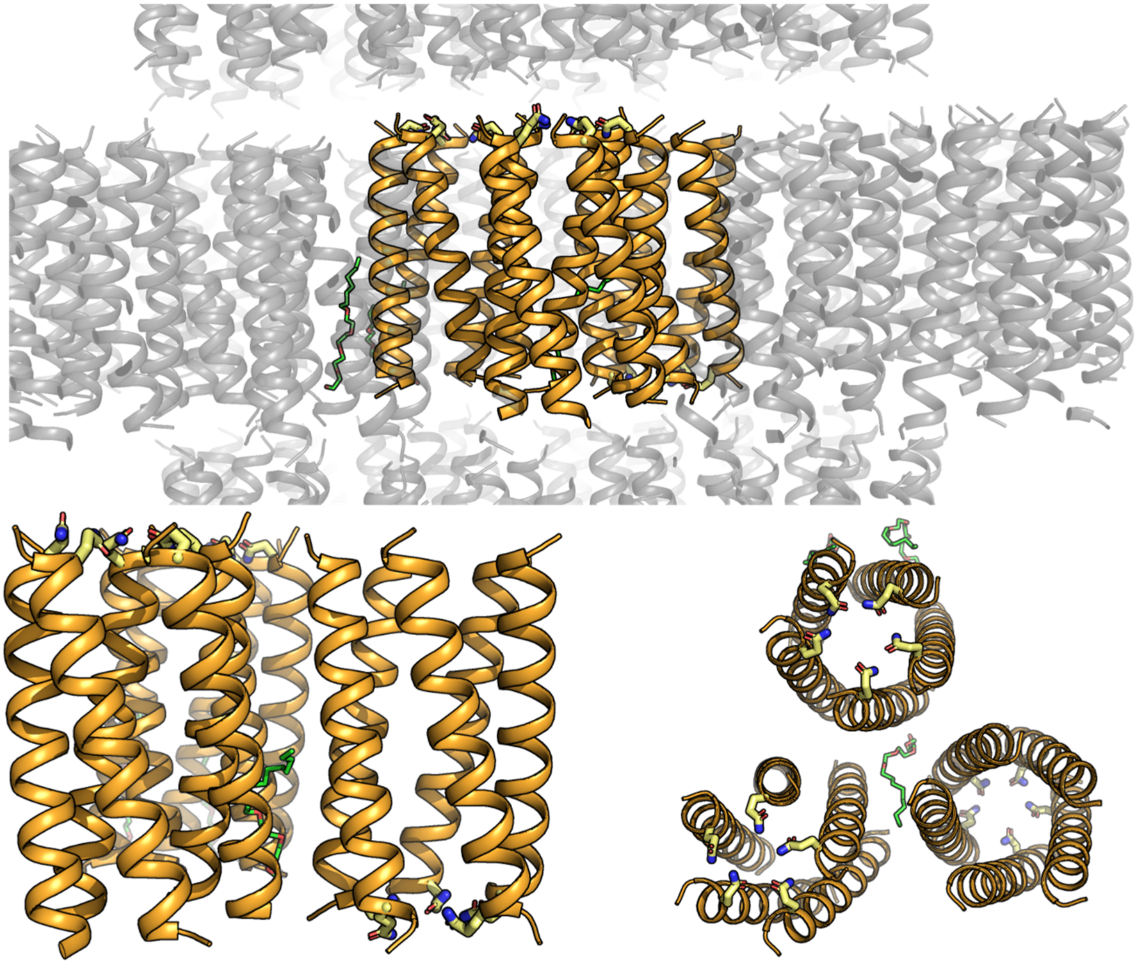

**Supplementary Fig. S7. Crystal lattice of QLLL. a,** Asymmetric unit (orange) in crystal lattice reveals three pentameric QLLL assemblies. **b,** Asymmetric unit is composed of three QLLL pentamers (2 up and 1 down). The three pentamers are structurally similar with rmsd values from 0.252 to 0.336 Å.

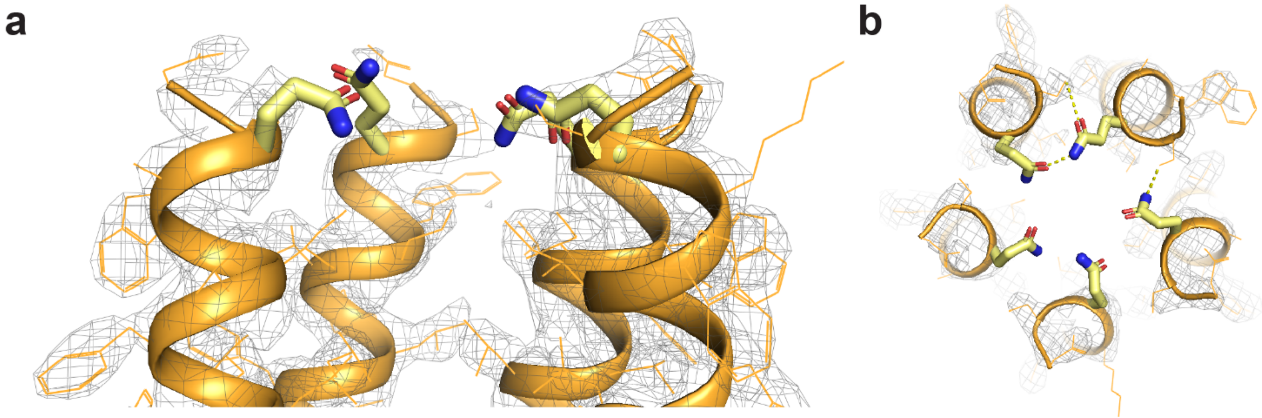

**Supplementary Fig. S8. Structure of QLLL with electron density. a,** Close-up of sideview with electron density (σ = 1.0) of one of the QLLL assemblies. **b,** Top view of Gln layer. Hydrogen bonds of < 3.2 Å shown in yellow dashes.

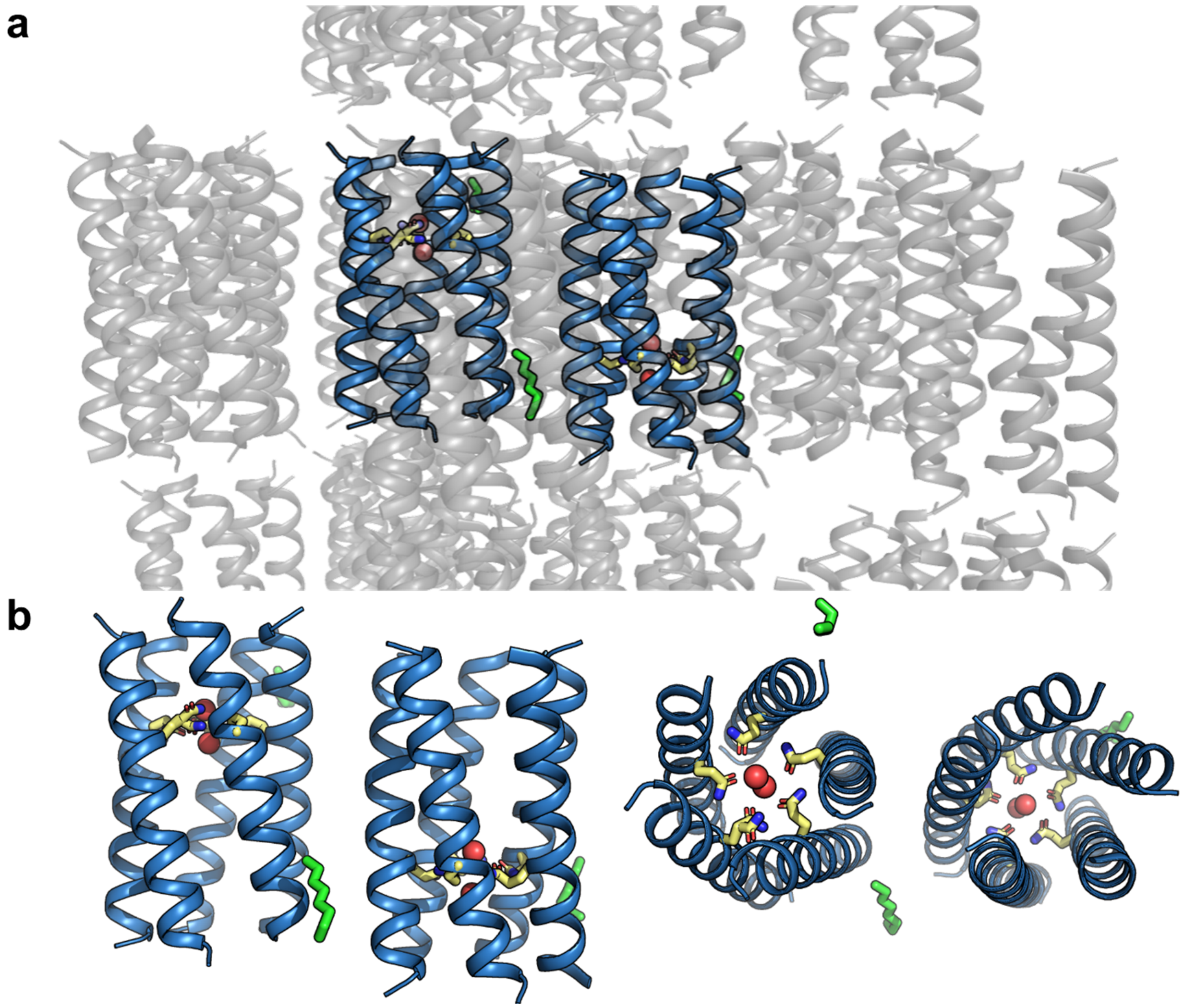

**Supplementary Fig. S9. Crystal lattice of LQLL. a,** Asymmetric unit (blue) in crystal lattice reveals two pentameric LQLL assemblies. **b,** Asymmetric unit is composed of two LQLL pentamers (one up and one down). The two structures in the asymmetric unit have an rmsd of 0.189 Å.

5

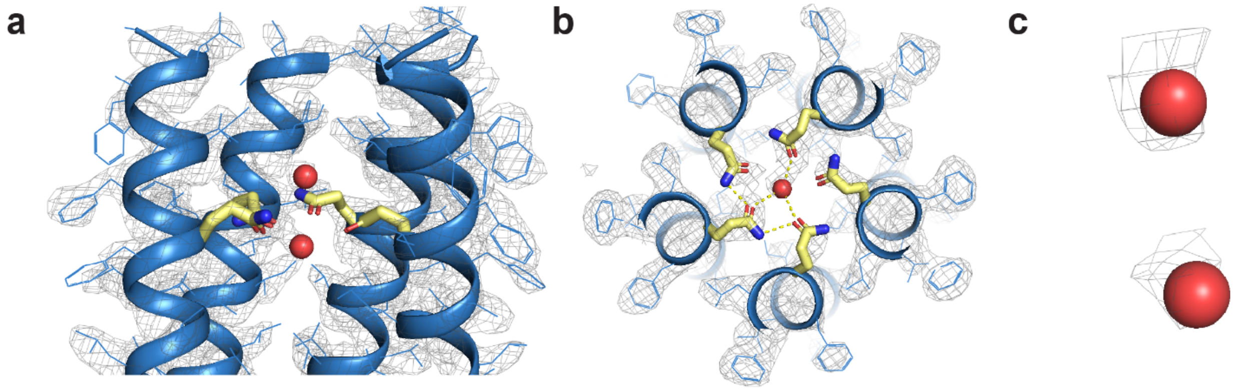

**Supplementary Fig. S10. Structure of LQLL with electron density. a,** Close-up of sideview with electron density ($\sigma$ = 1.0) of one of the LQLL assemblies. **b,** Top view of Gln layer. Hydrogen bonds of < 3.2 Å shown in yellow dashes. **c,** Electron density ($\sigma$ = 1.0) around waters in crystal structure.

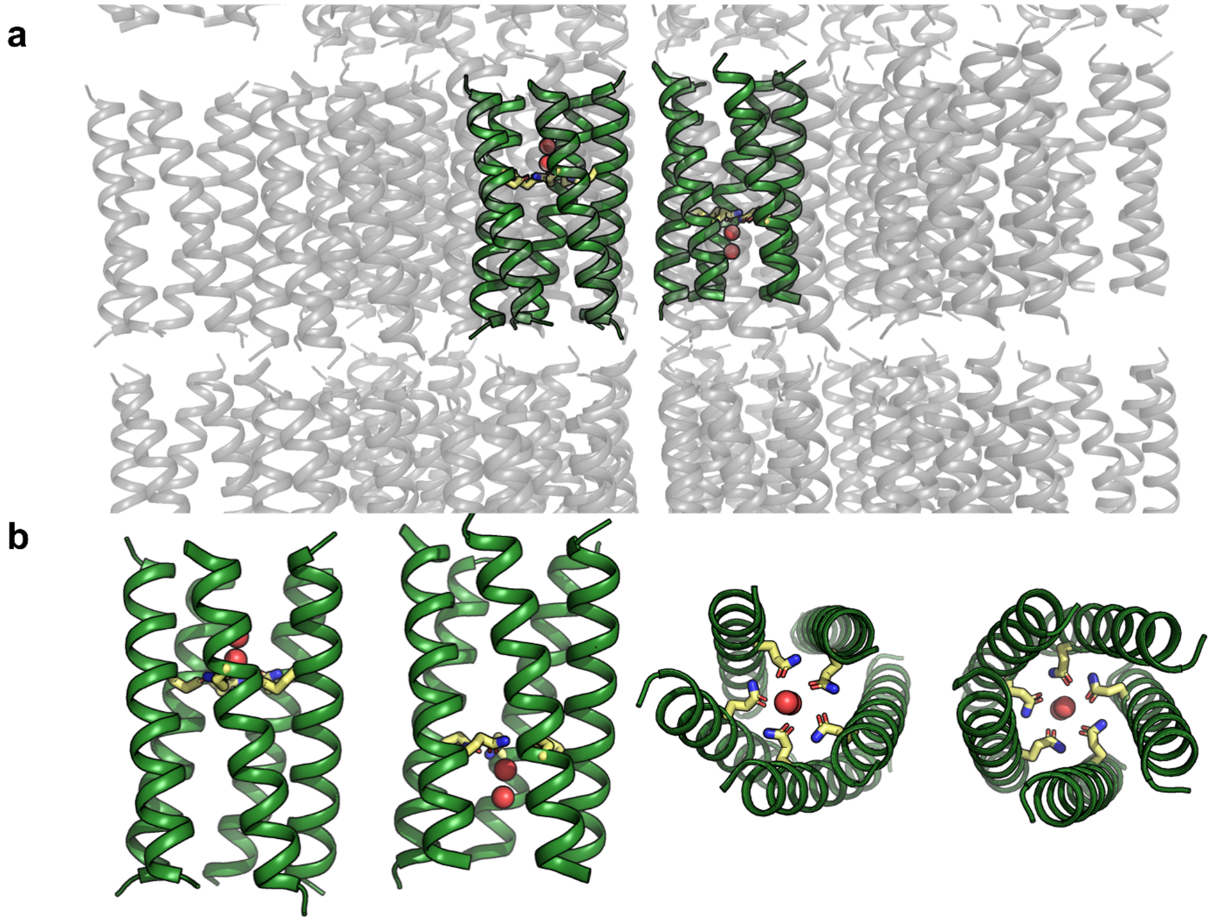

**Supplementary Fig. S11. Crystal lattice of LLQL. a,** Asymmetric unit (green) in crystal lattice reveals two pentameric LLQL assemblies. **b,** Asymmetric unit is composed of two LLQL pentamers (one up and one down). The structures of the two pentamers are within 0.337 Å rmsd of each other.

5

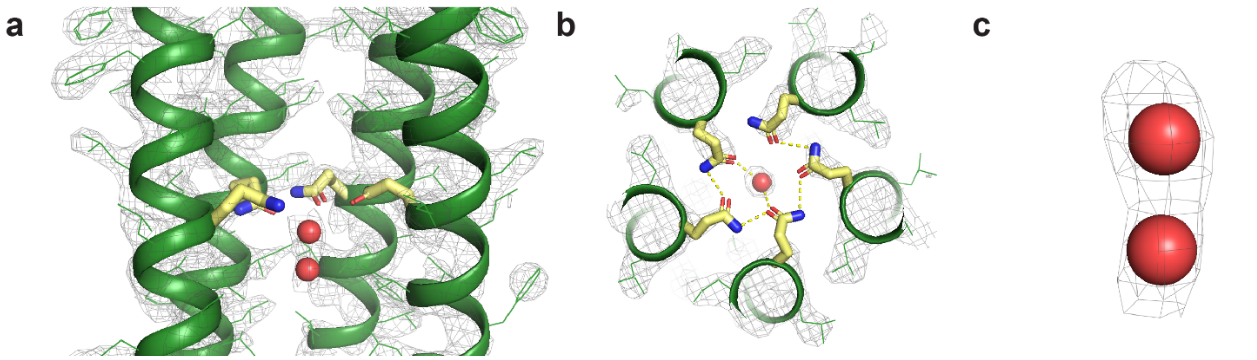

**Supplementary Fig. S12. Structure of LLQL with electron density. a,** Close-up of sideview with electron density (σ = 1.0) of one of the LLQL assemblies. **b,** Top view of Gln layer. Hydrogen bonds of < 3.2 Å shown in yellow dashes. **c,** Electron density (σ = 1.0) around waters in crystal structure.

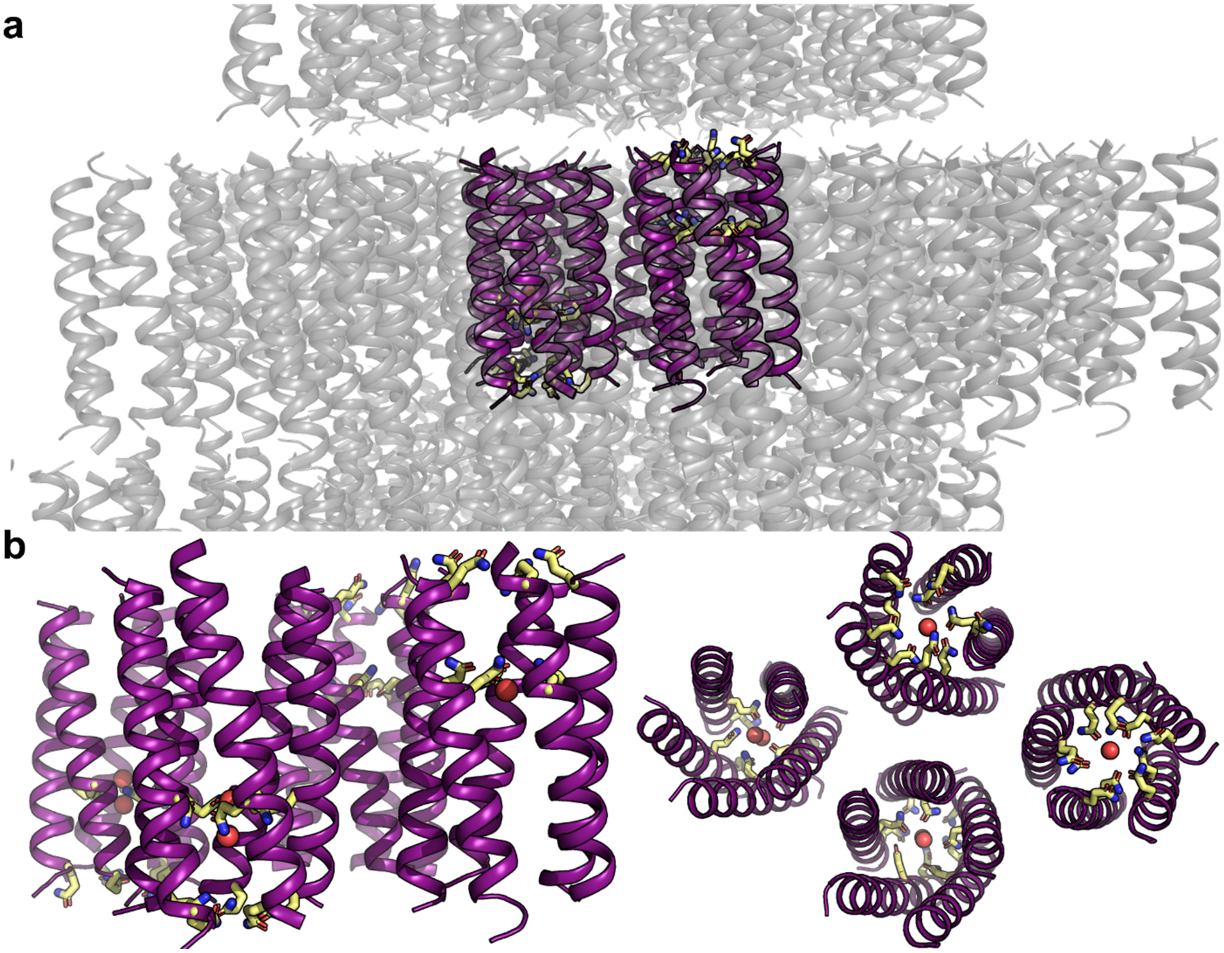

**Supplementary Fig. S13. Crystal lattice of QQLL. a,** Asymmetric unit (purple) in crystal lattice reveals four pentameric QQLL assemblies. **b,** Asymmetric unit is composed of four QQLL pentamers (2 up and 2 down). The backbone rmsd of the four bundles in the asymmetric unit vary from 0.275 to 1.62 Å.

5

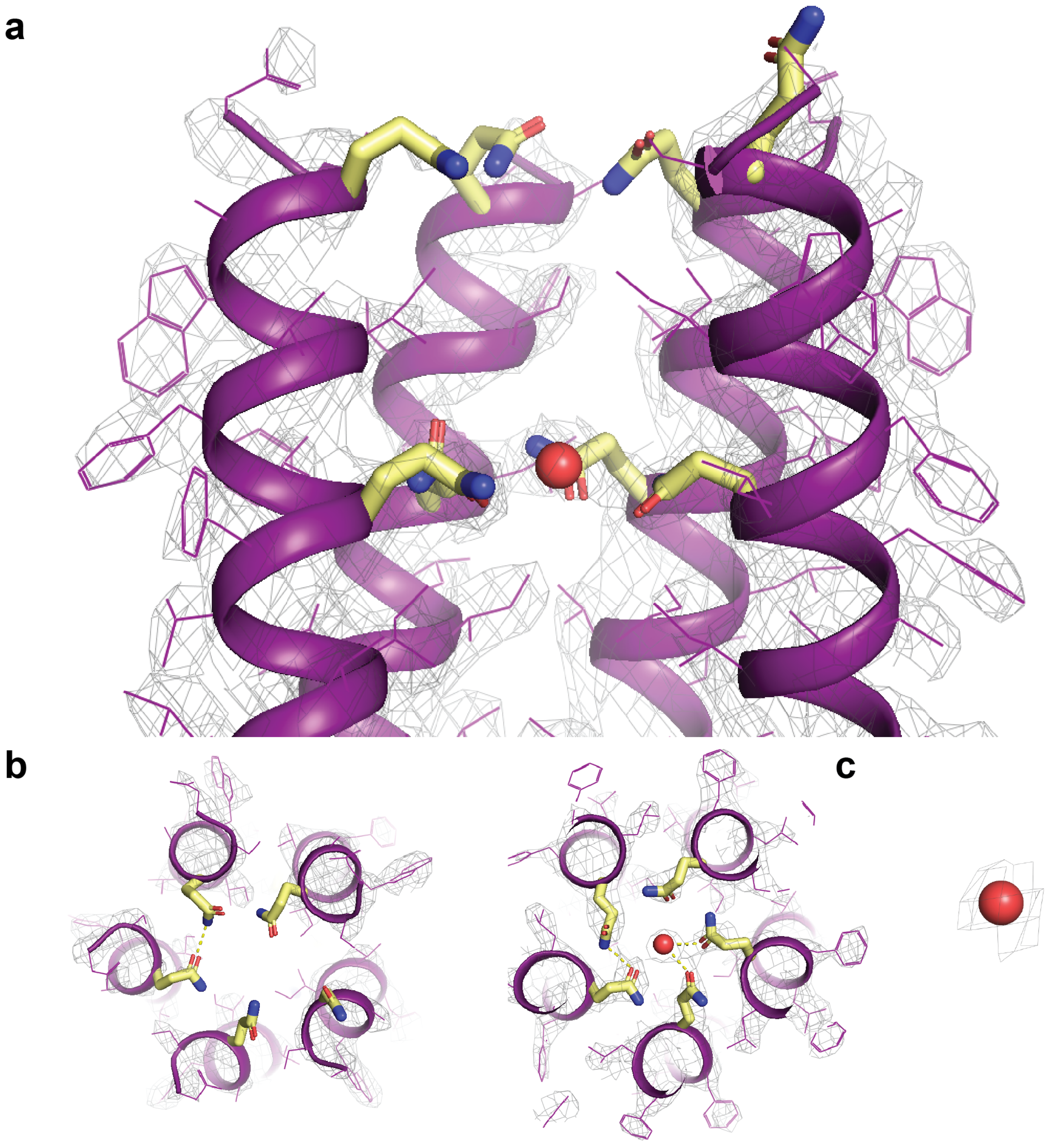

**Supplementary Fig. S14. Structure of QQLL with electron density. a,** Close-up of sideview with electron density of one of the QQLL assemblies. **b,**. Top view of Gln layer 1 (left) and Gln layer 2 (right). **c,** Electron density at σ = 1.0 for water.

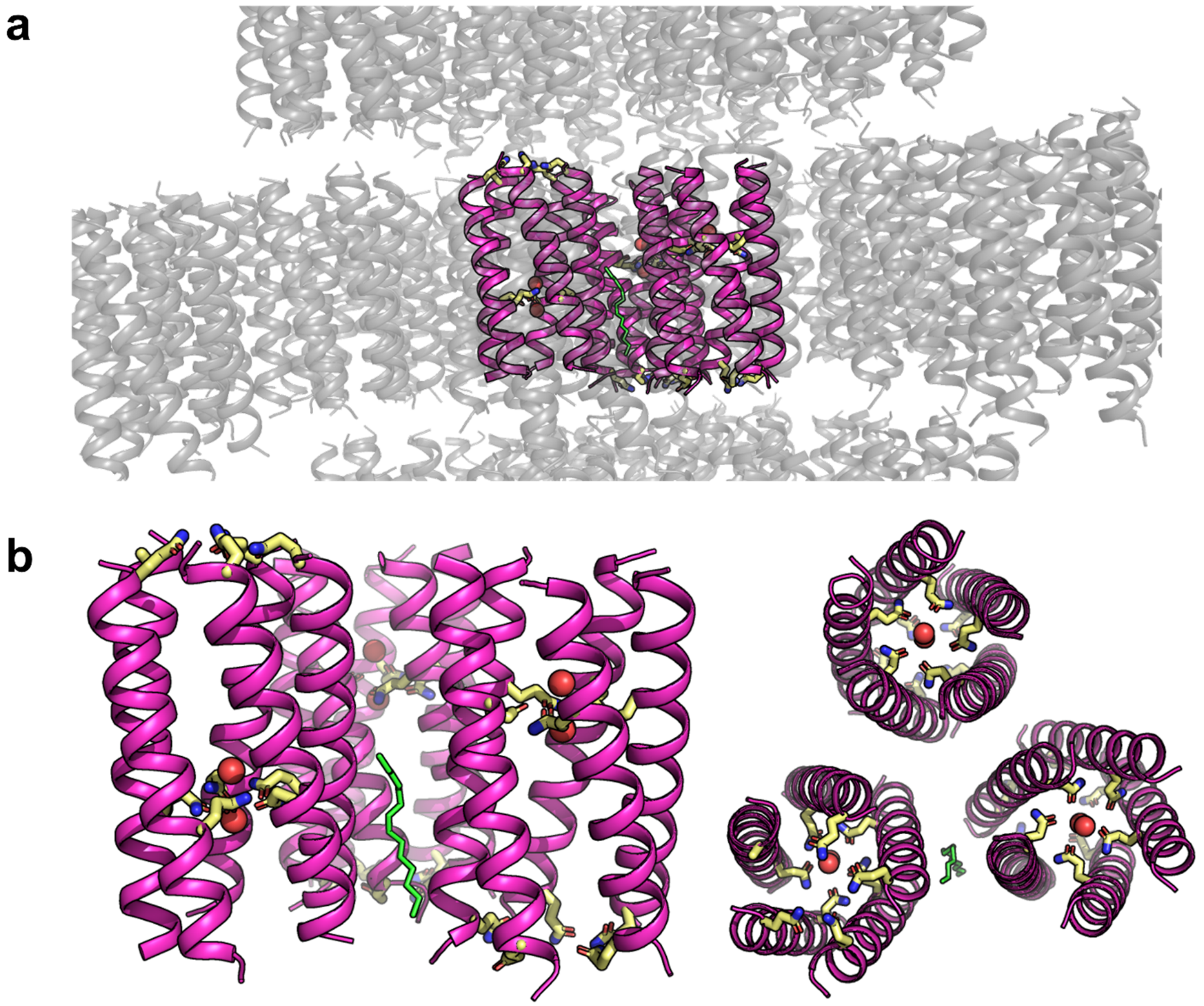

**Supplementary Fig. S15. Crystal lattice of QLQL. a,** Asymmetric unit (pink) in crystal lattice reveals three pentameric QLQL assemblies. **b,** Asymmetric unit is composed of three QLQL pentamers (2 up and 1 down). The three pentamers in the asymmetric unit have an rmsd between 0.376-0.427 Å.

5

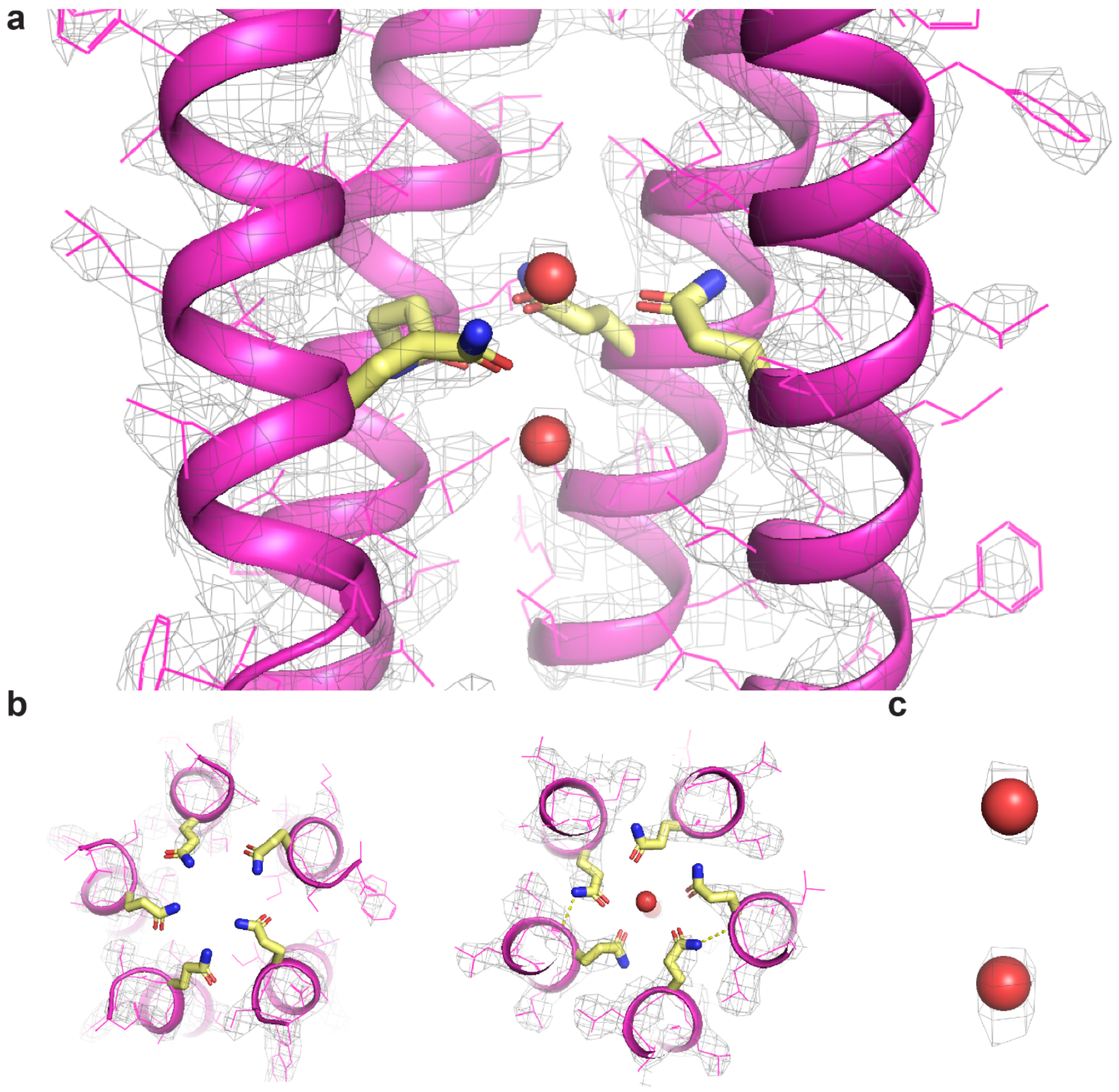

**Supplementary Fig. S16. Structure of QLQL with electron density. a,** Close-up of sideview with electron density (σ = 1.0) of one of the QLQL assemblies. **b,** Top view of Gln layer 1 (left) and Gln layer 2 (right). Hydrogen bonds of < 3.2 Å shown in yellow dashes. **c,** Electron density (σ = 1.0) around waters in crystal structure.

5

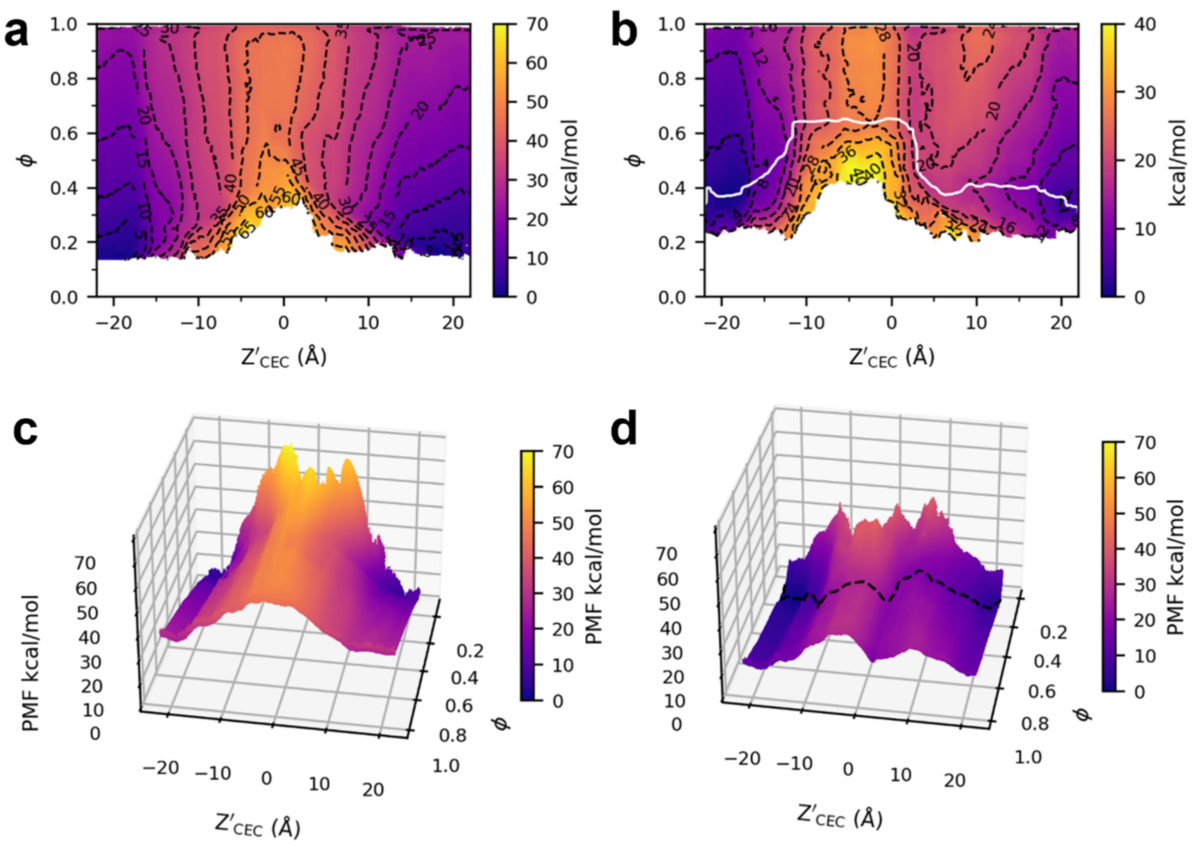

**Supplementary Fig. S17. Generating MFEP from 2D PMFs.** 2D PMFs for **a,** LLLL and **b,** LQLL, respectively. Note the different color scheme used for each for better contrast. Contours shown and labeled in each. In **b,**, the white line is the MFEP calculated using string theory. 3D depictions of the same PMFs for **c,** LLLL and **d,** LQLL, respectively, using the same scaling. The black dashed line in **d,** represents the MFEP.

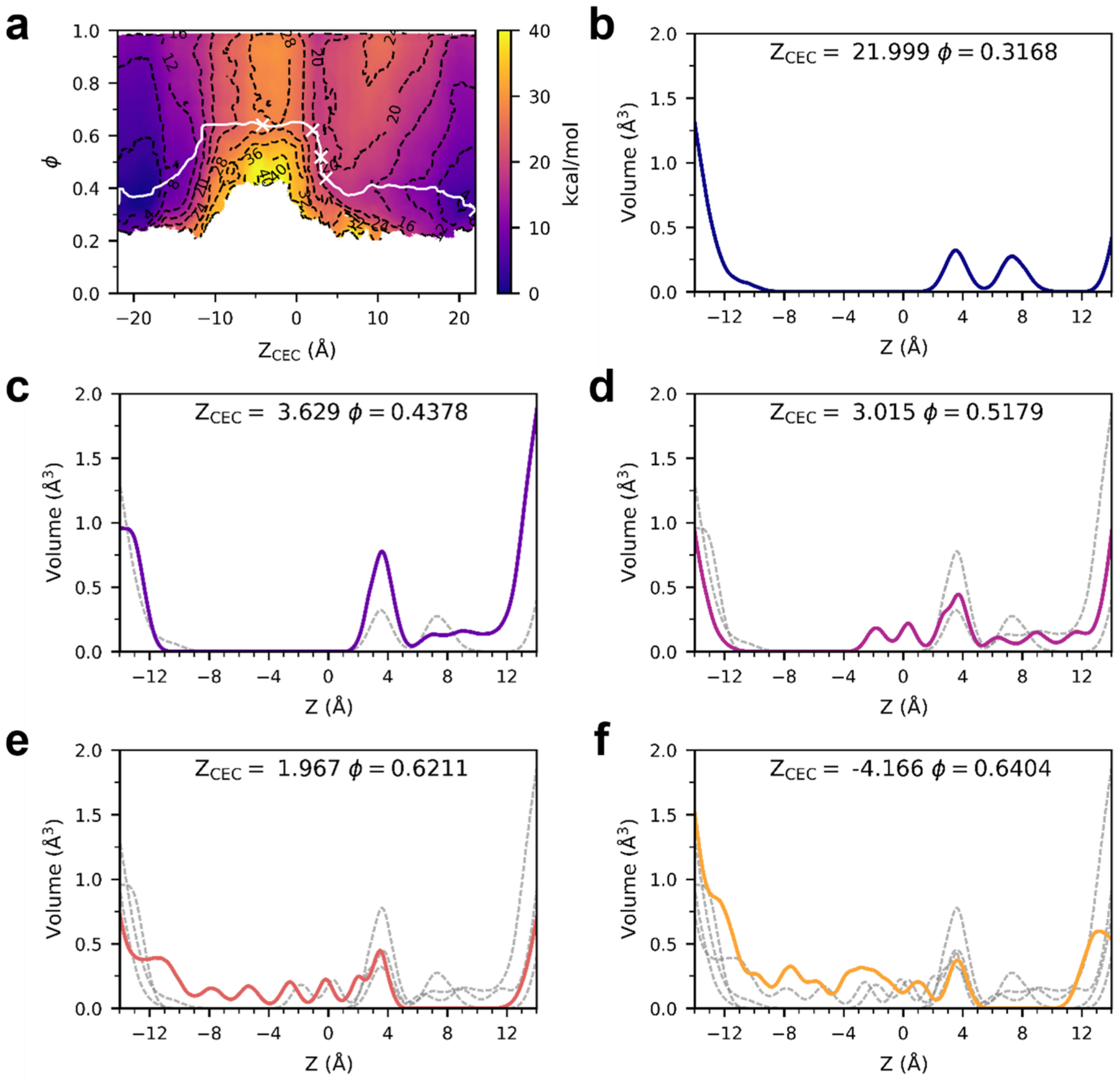

**Supplementary Fig. S18. Average water volume as a function of Z'$_{CEC}$ and ϕ. a,** Average volume of water in the channel shown for 5 different points along the LQLL MFEP, as indicated by white X's. **b,** The excess proton is out of the channel and the few waters in the channel are not connected to bulk. **c,** The proton is just below Gln and water connects to the top. **d,** The water structure transitions from the top to the bottom while the proton remains below Gln. **e,** The water is full connected in the bottom. **f,** The proton moves through the bottom-connected water wire. The dashed lines in C-F are the curves shown in all previous panels, b-e.

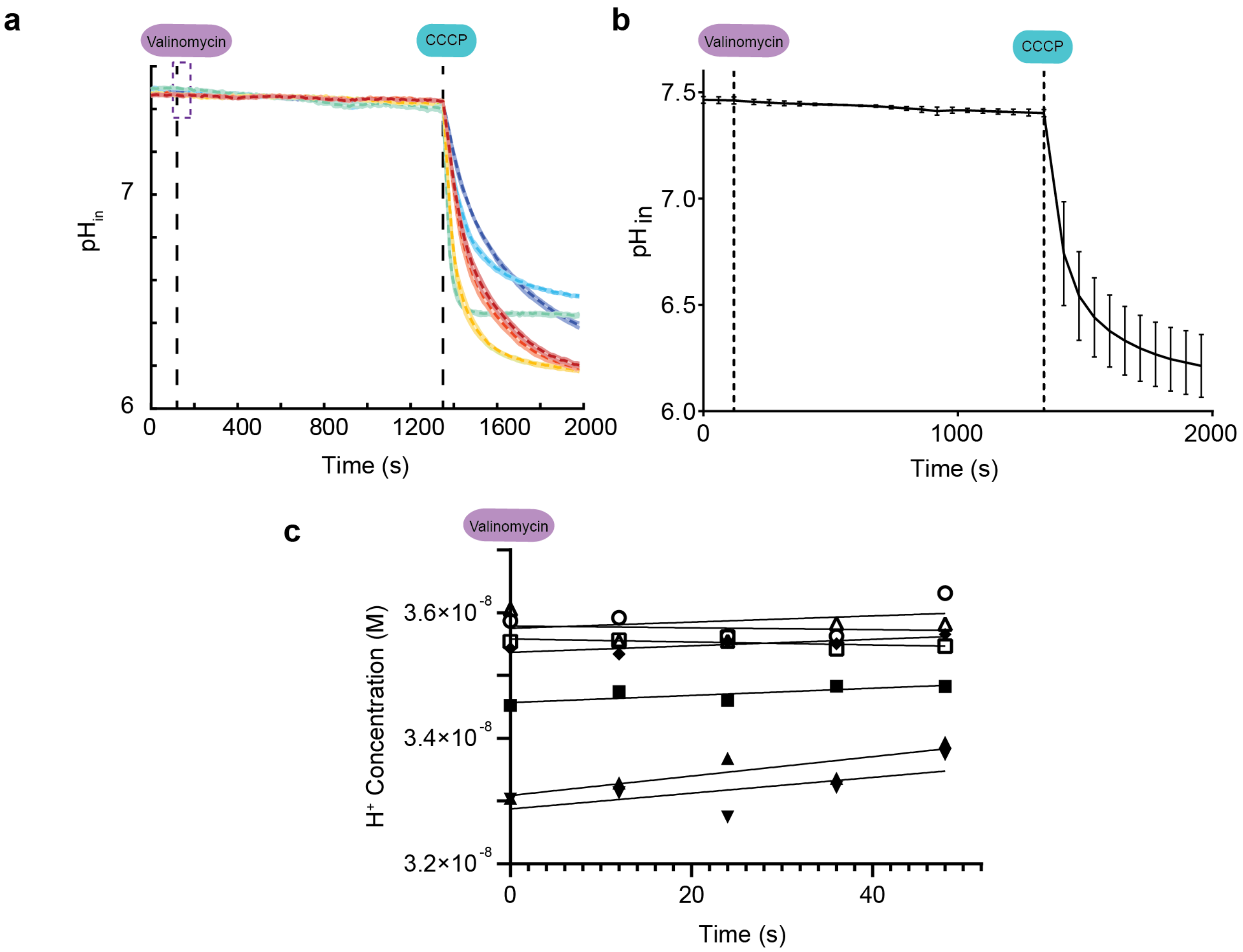

**Supplementary Fig. S19. All proton flux assay data for "Empty" vesicles.** Seven samples (each run in triplicate with shaded error bars shown) for empty (non-proteinaceous) vesicles that were run independently in assay. **a,** $pH_{in}$ as a function of time throughout the measurement for each independent sample. Purple box shows time points used in analysis of initial rates in **c**. **b,** Mean and standard deviation for data collection. Data prior to CCCP step shown in Fig. 4b. **c,** Fits for the initial 50 seconds following addition of valinomycin. From the linear regression fits, all slopes (which give the initial rates (in M/s)) were used to calculate mean and standard error presented in Fig. 6g and Supplementary Table S3.

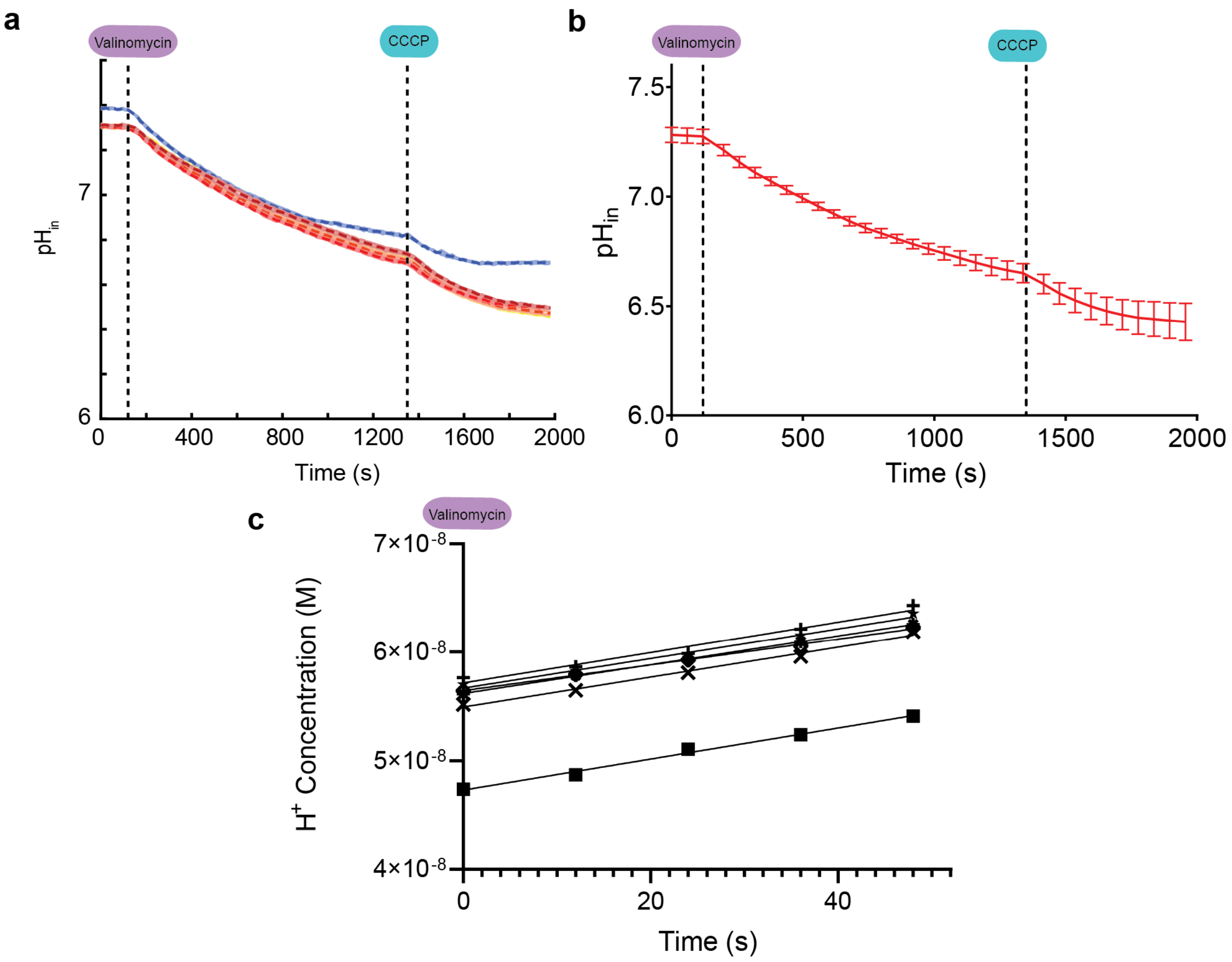

**Supplementary Fig. S20. All proton flux assay data for Influenza A M2 channel vesicle samples.** Six samples (each run in triplicate with shaded error bars shown) containing 1:500 polypeptide:lipid ratio; samples were run independently in the assay. **a,** $pH_{in}$ as a function of time throughout the measurement for each independent sample. **b,** Mean and standard deviation for data collection. Data prior to CCCP step shown in Fig. 4c. **c,** Fits for the initial 50 seconds following addition of valinomycin. From the linear regression fits, all slopes (which give the initial rates (in M/s)) were used to calculate mean and standard error presented in Fig. 6g and Supplementary Table S3.

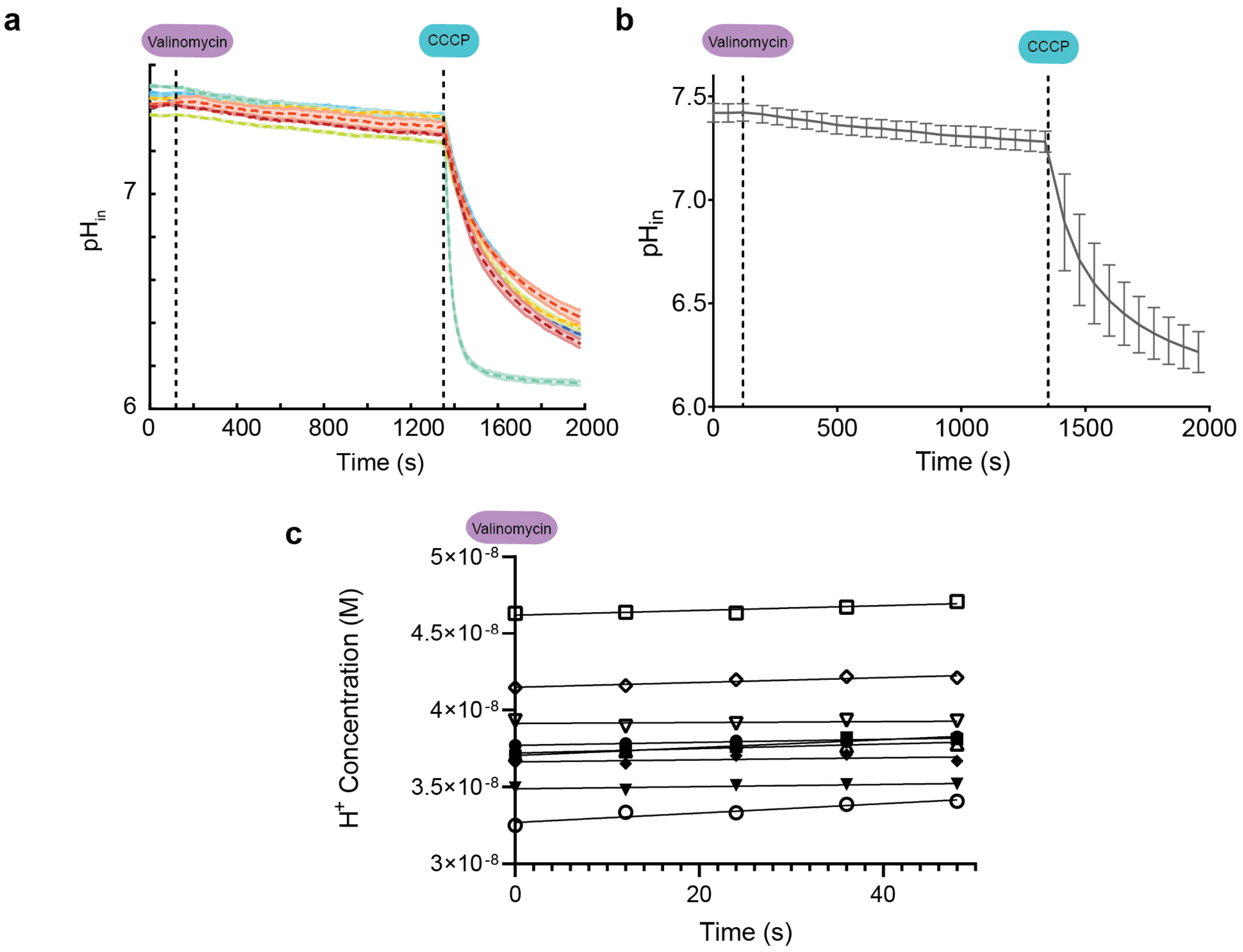

**Supplementary Fig. S21. All proton flux assay data for LLLL vesicle samples.** Nine samples (each run in triplicate with shaded error bars shown) containing 1:500 peptide:lipid ratio; samples were run independently in the assay. **a,** $pH_{in}$ as a function of time throughout the measurement for each independent sample. **b,** Mean and standard deviation for data collection. Data prior to CCCP step shown in Fig. 4d. **c,** Fits for the initial 50 seconds following addition of valinomycin. From the linear regression fits, all slopes (which give the initial rates (in M/s)) were used to calculate mean and standard error presented in Fig. 6g and Supplementary Table S3.

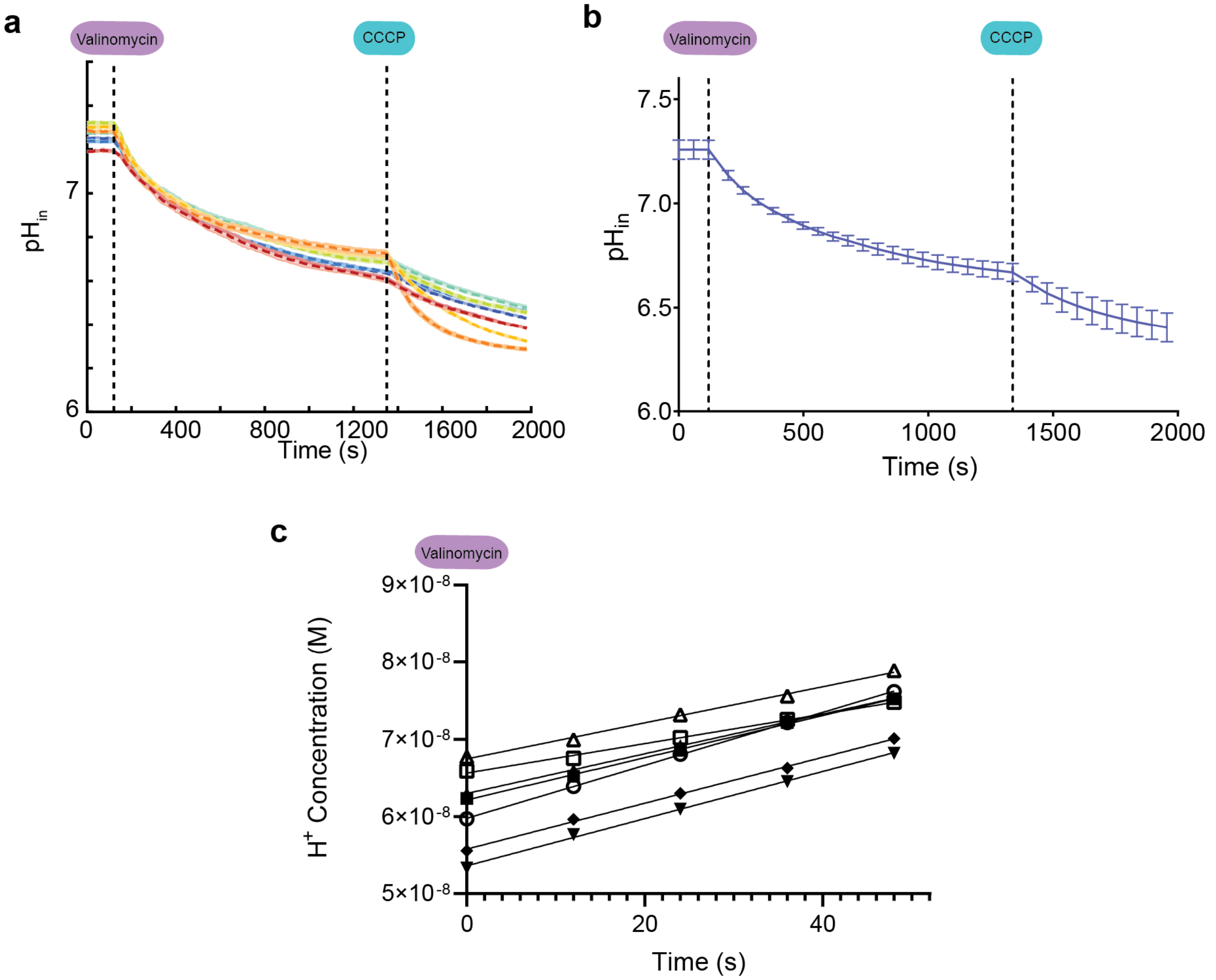

**Supplementary Fig. S22. All proton flux assay data for LQLL vesicle samples.** Seven samples (each run in triplicate with shaded error bars shown) containing 1:500 peptide:lipid ratio; samples were run independently in the assay. **a,** $pH_{in}$ as a function of time throughout the measurement for each independent sample. **b,** Mean and standard deviation for data collection. Data prior to CCCP addition shown in Fig. 4e. **d,** Fits for the initial 50 seconds following addition of valinomycin. From the linear regression fits, all slopes (which give the initial rates (in M/s)) were used to calculate mean and standard error presented in Fig. 6g and Supplementary Table S3.

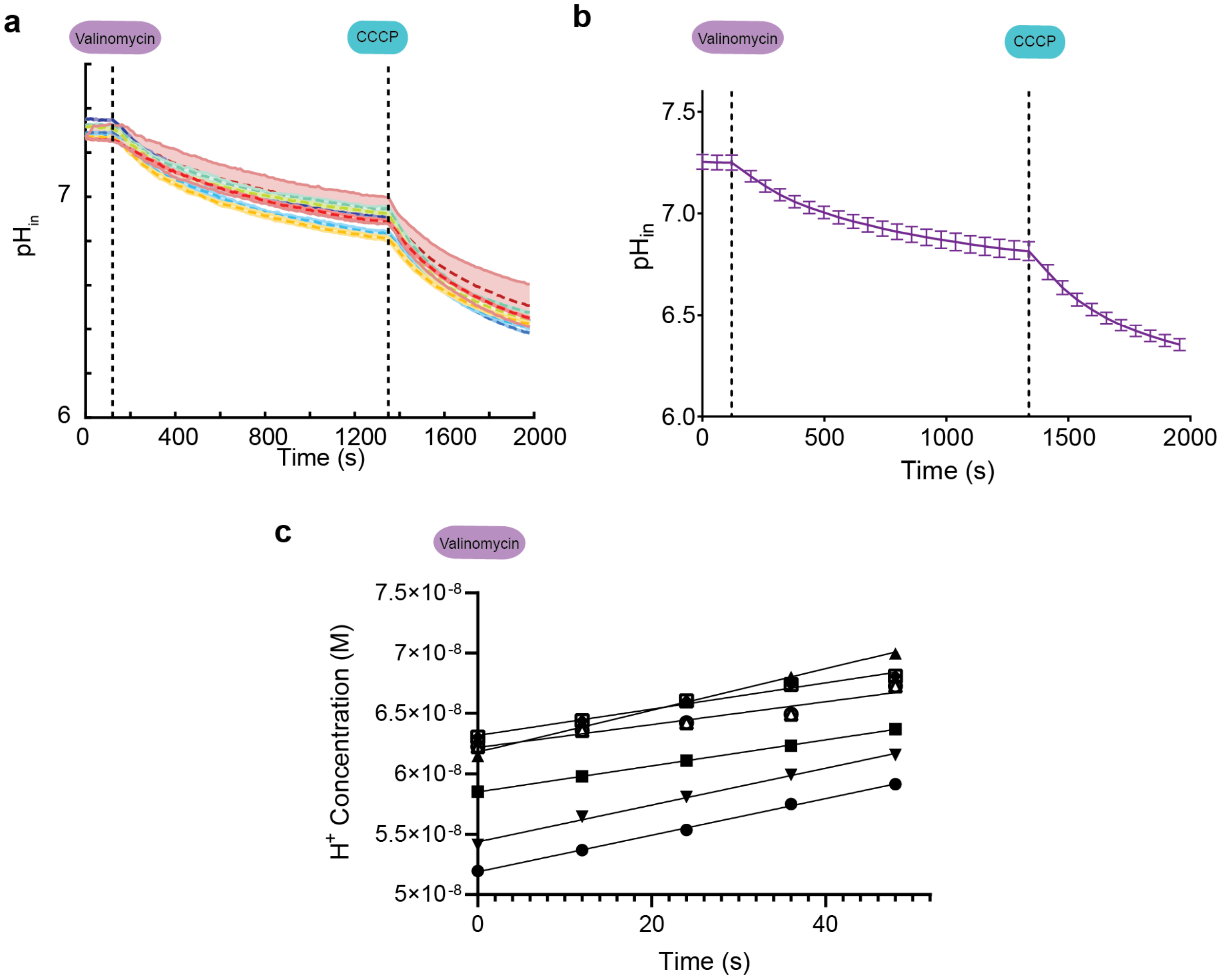

**Supplementary Fig. S23. All proton flux assay data for QQLL vesicle samples.** Eight samples (each run in triplicate with shaded error bars shown) containing 1:500 peptide:lipid ratio; samples were run independently in the assay. **a,** $pH_{in}$ as a function of time throughout the measurement for each independent sample. **b,** Mean and standard deviation for data collection. Data prior to CCCP addition shown in Fig. 4f. **d,** Fits for the initial 50 seconds following addition of valinomycin. From the linear regression fits, all slopes (which give the initial rates (in M/s)) were used to calculate mean and standard error presented in Fig. 6g and Supplementary Table S3.

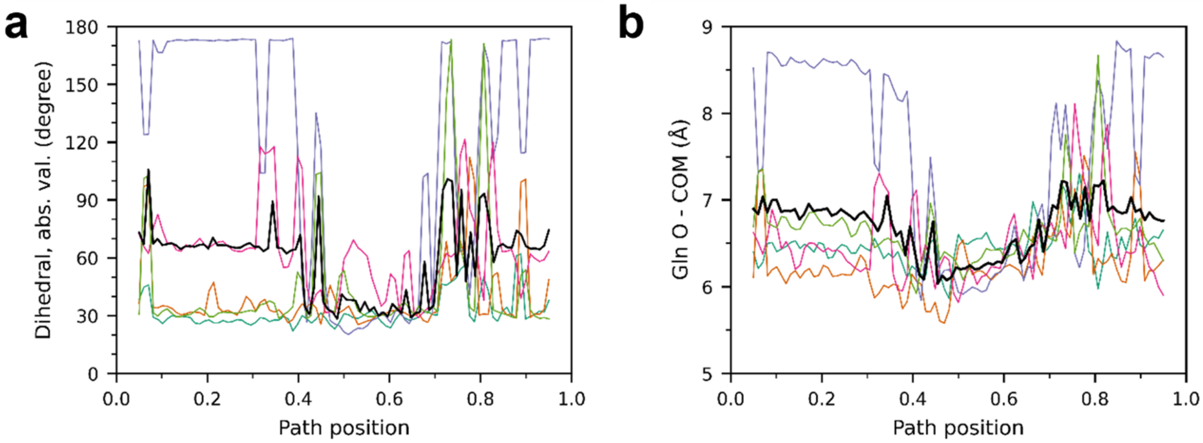

**Supplementary Fig. S24. Analysis of Gln sidechain dihedrals reveals asymmetry in Gln conformations. a,** The absolute value of the Gln sidechain dihedral for each Gln, as a function of the system position along the path. The average over all 5 residues is shown in black. **b,** The distance between the Gln carbonyl oxygen and the Ile13 center of mass, shown for each Gln. The average over all 5 residues is shown in black

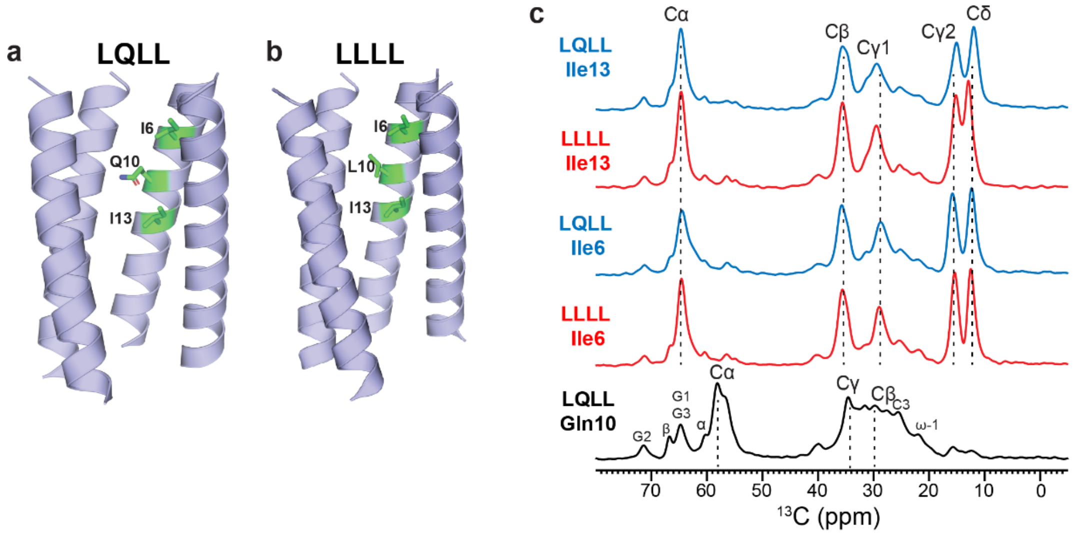

**Supplementary Figure 25. Structure and 1D $^{13}$C NMR spectra of membrane-bound LQLL and LLLL peptides**. (a) Crystal structure of pentameric LQLL (pdb: 7udz), highlighting Ile6, Gln10, Ile13, which are isotopically labeled in this study. Fifth helix removed for clarity. (b) Crystal structure of pentameric LLLL (PDB: 6MCT). Ile6 and Ile13 are isotopically labeled in this study. One chain is removed for clarity. (c) 1D $^{13}$C CP-MAS spectra of five DMPC membrane-bound LQLL and LLLL peptides, each containing a single labeled residue. All Ile and Gln Cα and Cβ chemical shifts indicate α-helical conformation. Minor signals of the DMPC natural abundance headgroup and glycerol backbone carbons are also observed in the Gln10 spectra. Ile6 in LLLL shows narrower linewidths than the other Ile residues in the other peptides, indicating higher conformational homogeneity. Gln10 exhibits broader linewidths, indicating conformational heterogeneity.

**Supplementary Table S1. Designed peptide sequences and observed masses.**

Peptides used for the experiments were synthesized and purified using protocols described in Materials and Methods. Samples were calibrated to bovine insulin (Sigma Aldrich) $[M+H]^{+}$ and $[M+2H]^{2+}$peaks.

| Design | Sequence | Expected Mass (Da) | Observed Mass (Da) |
|---|---|---|---|
| LLLL | DS $^{3}$LKWIVFL $^{10}$LFLIVLL $^{17}$LLAIVFL LRG | 3028.1 | 3030.4 |
| QLLL | DS $^{3}$QKWIVFL $^{10}$LFLIVLL $^{17}$LLAIVFL LRG | 3043.1 | 3045.0 |
| LQLL | DS $^{3}$LKWIVFL $^{10}$QFLIVLL $^{17}$LLAIVFL LRG | 3043.1 | 3045.2 |
| LLQL | DS $^{3}$LKWIVFL $^{10}$LFLIVLL $^{17}$QLAIVFL LRG | 3043.1 | 3044.4 |
| QQLL | DS $^{3}$QKWIVFL $^{10}$QFLIVLL $^{17}$LLAIVFL LRG | 3058.0 | 3059.2 |
| QLQL | DS $^{3}$QKWIVFL $^{10}$LFLIVLL $^{17}$QLAIVFL LRG | 3058.0 | 3059.3 |

**Supplementary Table S2. Protein crystallization conditions**

Solutions and conditions for crystallization of all designed proton channels. All crystals prepped using LCP methods described in Materials and Methods at 30 mg/mL in MAG 9.9 with 50 mM OG.

| Design | Crystallization Condition |
| --- | --- |
| QLLL | 0.22 M sodium citrate, 0.1 M Tris pH 8.0, 35% v/v PEG 400 |
| LQLL | 0.1 M calcium chloride, 0.1 M MES, 29% v/v PEG 400 |
| LLQL | 0.2 M magnesium chloride, 0.1 M potassium chloride, 0.03 M sodium citrate pH 4, 33% v/v PEG 400 |
| QQLL | 0.22 M sodium citrate, 0.1 M Tris pH 8.0 |
| QLQL | 0.2 M ammonium sulfate, 0.1 M sodium chloride, 0.1 M sodium citrate pH 6.0, 20% w/v PEG 2000 |

**Supplementary Table S3. Descriptive statistics for all vesicle samples in proton flux assays.** Mean and standard deviation/error of initial rates for all vesicle samples run for data presented in Fig. 6, Extended Data Figs. 3-5, Supplementary Figs. S19-S23.

| | Empty | Influenza A M2 | LLLL | QLLL | LQLL | LLQL | QQLL | QLQL |
|---|---|---|---|---|---|---|---|---|
| **Number of values** | 7 | 6 | 9 | 8 | 7 | 8 | 8 | 8 |
| **Minimum** | -2.4E-12 | 1.19E-10 | 2.97E-12 | 1.86E-11 | 1.91E-10 | 1.52E-10 | 9.49E-11 | 1.35E-10 |
| **Maximum** | 1.55E-11 | 1.43E-10 | 3.06E-11 | 4.34E-11 | 3.43E-10 | 1.77E-10 | 1.73E-10 | 1.78E-10 |
| **Range** | 1.78E-11 | 2.38E-11 | 2.76E-11 | 2.49E-11 | 1.52E-10 | 2.49E-11 | 7.77E-11 | 4.28E-11 |
| **Mean** | 5.72E-12 | 1.35E-10 | 1.43E-11 | 3.3E-11 | 2.72E-10 | 1.65E-10 | 1.24E-10 | 1.53E-10 |
| **Std. Deviation** | 6.58E-12 | 8.49E-12 | 9.05E-12 | 1.05E-11 | 5.02E-11 | 1.08E-11 | 3.01E-11 | 1.56E-11 |
| **Std. Error of Mean** | 2.49E-12 | 3.47E-12 | 3.02E-12 | 3.7E-12 | 1.9E-11 | 3.81E-12 | 1.07E-11 | 5.51E-12 |
| **Lower 95% CI of mean** | -3.7E-13 | 1.26E-10 | 7.36E-12 | 2.42E-11 | 2.25E-10 | 1.56E-10 | 9.88E-11 | 1.4E-10 |
| **Upper 95% CI of mean** | 1.18E-11 | 1.44E-10 | 2.13E-11 | 4.17E-11 | 3.18E-10 | 1.74E-10 | 1.49E-10 | 1.66E-10 |

**Supplementary Table S4. Unpaired t-test for control samples: empty vesicle initial rates vs. Influenza A M2 proton channel.** Comparison of initial rates for the control samples reveals that rates measured for vesicles containing the natural proton channel (influenza A M2 channel) are statistically significant from empty vesicles.

| Unpaired t test between Influenza A M2 channel and empty vesicles | |
|---|---|
| **P value** | <0.0001 |
| **P value summary** | **** |
| **Significantly different (P < 0.05)?** | Yes |
| **One- or two-tailed P value?** | Two-tailed |
| **t, df** | t=30.93, df=11 |

**Supplementary Table S5. Summary of ordinary one-way ANOVA for multiple comparisons (Dunn's test) of LLLL vs. designed channels.** Comparison of initial rates from Figure 6g shows that initial rates of LLLL and QLLL are not statistically significant. However, initial rates of conduction for LQLL, LLQL, QQLL and QLQL are statistically significant when compared to LLLL.

| Dunnett's multiple comparisons test | Mean Diff. | 95.00% CI of diff. | Summary | Adjusted P Value |
|---|---|---|---|---|
| LLLL vs. QLLL | -1.9E-11 | -5.001E-011 to 1.270E-011 | ns | 0.405 |
| LLLL vs. LQLL | -2.6E-10 | -2.898E-010 to -2.248E-010 | **** | <0.0001 |
| LLLL vs. LLQL | -1.5E-10 | -1.816E-010 to -1.189E-010 | **** | <0.0001 |
| LLLL vs. QQLL | -1.1E-10 | -1.411E-010 to -7.836E-011 | **** | <0.0001 |
| LLLL vs. QLQL | -1.4E-10 | -1.696E-010 to -1.069E-010 | **** | <0.0001 |

**Supplementary Table S6. Summary of ordinary one-way ANOVA for multiple comparisons (Dunn's test) of QLLL vs. designed channels.** Comparing the initial rates of proton flux of all channels to QLLL. Similarly, this analysis reveals that there is a statistically significant difference between initial rates of the non-conductive QLLL/LLLL with the other four proton-conductive designed channels.

| Dunnett's multiple comparisons test | Mean Diff. | 95.00% CI of diff. | Summary | Adjusted P Value |
|---|---|---|---|---|
| QLLL vs. LLLL | 1.87E-11 | -1.258E-011 to 4.988E-011 | ns | 0.3947 |
| QLLL vs. LQLL | -2.4E-10 | -2.719E-010 to -2.054E-010 | **** | <0.0001 |
| QLLL vs. LLQL | -1.3E-10 | -1.638E-010 to -9.948E-011 | **** | <0.0001 |
| QLLL vs. QQLL | -9.1E-11 | -1.232E-010 to -5.893E-011 | **** | <0.0001 |
| QLLL vs. QLQL | -1.2E-10 | -1.517E-010 to -8.747E-011 | **** | <0.0001 |

**Supplementary Movie S1.** Snapshots from MS-RMD simulations along MFEP (as shown in Fig. 5) depict the proton (represented as a yellow hydronium in the movie) inducing the formation of water wires in hydrophobic regions of the channel lumen. The proton moves along these transiently forming water wires to make its way down to a region just below the PLS formed by the green Gln residues. Once near this site, the presence of the proton enables the formation of a secondary water wire through the longer hydrophobic region. This allows the proton to traverse the rest of the way across the channel.

**References and notes:**

1 Moriyama, Y. & Futai, M. H+-ATPase, a primary pump for accumulation of neurotransmitters, is a major constituent of brain synaptic vesicles. *Biochemical and biophysical research communications* **173**, 443-448, doi:10.1016/s0006-291x(05)81078-2 (1990).

2 Nishi, T. & Forgac, M. The vacuolar (H+)-ATPases--nature's most versatile proton pumps. *Nature reviews. Molecular cell biology* **3**, 94-103, doi:10.1038/nrm729 (2002).

3 Mitchell, P. Coupling of phosphorylation to electron and hydrogen transfer by a chemi-osmotic type of mechanism. *Nature* **191**, 144-148, doi:10.1038/191144a0 (1961).

4 Nicholls, D. G. Mitochondrial ion circuits. *Essays in biochemistry* **47**, 25-35, doi:10.1042/bse0470025 (2010).

5 Diering, G. H. & Numata, M. Endosomal pH in neuronal signaling and synaptic transmission: role of Na(+)/H(+) exchanger NHE5. *Frontiers in physiology* **4**, 412, doi:10.3389/fphys.2013.00412 (2014).

6 Agmon, N. The Grotthuss mechanism. *Chemical Physics Letters* **244**, 456-462, doi:10.1016/0009-2614(95)00905-j (1995).

7 Calio, P. B., Li, C. & Voth, G. A. Resolving the Structural Debate for the Hydrated Excess Proton in Water. *J. Am. Chem. Soc.* **143**, 18672-18683, doi:10.1021/jacs.1c08552 (2021).

8 Li, C. & Voth, G. A. A quantitative paradigm for water-assisted proton transport through proteins and other confined spaces. *Proceedings of the National Academy of Sciences of the United States of America* **118**, doi:10.1073/pnas.2113141118 (2021).

9 Wraight, C. A. Chance and design--proton transfer in water, channels and bioenergetic proteins. *Biochimica et biophysica acta* **1757**, 886-912, doi:10.1016/j.bbabio.2006.06.017 (2006).

10 Decoursey, T. E. Voltage-gated proton channels and other proton transfer pathways. *Physiol Rev* **83**, 475-579, doi:10.1152/physrev.00028.2002 (2003).

11 Peng, Y., Swanson, J. M., Kang, S. G., Zhou, R. & Voth, G. A. Hydrated Excess Protons Can Create Their Own Water Wires. *The journal of physical chemistry. B* **119**, 9212-9218, doi:10.1021/jp5095118 (2015).

12 Banh, R. *et al.* Hydrophobic gasket mutation produces gating pore currents in closed human voltage-gated proton channels. *Proceedings of the National Academy of Sciences of the United States of America* **116**, 18951-18961, doi:10.1073/pnas.1905462116 (2019).

13 Garczarek, F. & Gerwert, K. Functional waters in intraprotein proton transfer monitored by FTIR difference spectroscopy. *Nature* **439**, 109-112, doi:10.1038/nature04231 (2006).

14 Kaur, D., Khaniya, U., Zhang, Y. & Gunner, M. R. Protein Motifs for Proton Transfers That Build the Transmembrane Proton Gradient. *Frontiers in chemistry* **9**, 660954, doi:10.3389/fchem.2021.660954 (2021).

15 Kalra, A., Garde, S. & Hummer, G. Osmotic water transport through carbon nanotube membranes. *Proceedings of the National Academy of Sciences of the*

*United States of America* **100**, 10175-10180, doi:10.1073/pnas.1633354100 (2003).

16 Ben-Abu, Y., Zhou, Y., Zilberberg, N. & Yifrach, O. Inverse coupling in leak and voltage-activated K+ channel gates underlies distinct roles in electrical signaling. *Nature structural & molecular biology* **16**, 71-79, doi:10.1038/nsmb.1525 (2009).

17 Jensen, M. O. *et al.* Principles of conduction and hydrophobic gating in K+ channels. *Proceedings of the National Academy of Sciences of the United States of America* **107**, 5833-5838, doi:10.1073/pnas.0911691107 (2010).

18 Aryal, P., Sansom, M. S. & Tucker, S. J. Hydrophobic gating in ion channels. *J Mol Biol* **427**, 121-130, doi:10.1016/j.jmb.2014.07.030 (2015).

19 Zhu, F. & Hummer, G. Drying transition in the hydrophobic gate of the GLIC channel blocks ion conduction. *Biophysical journal* **103**, 219-227, doi:10.1016/j.bpj.2012.06.003 (2012).

20 Rasaiah, J. C., Garde, S. & Hummer, G. Water in nonpolar confinement: from nanotubes to proteins and beyond. *Annual review of physical chemistry* **59**, 713-740, doi:10.1146/annurev.physchem.59.032607.093815 (2008).

21 Wang, T. *et al.* Deprotonation of D96 in bacteriorhodopsin opens the proton uptake pathway. *Structure* **21**, 290-297, doi:10.1016/j.str.2012.12.018 (2013).

22 Weinert, T. *et al.* Proton uptake mechanism in bacteriorhodopsin captured by serial synchrotron crystallography. *Science* **365**, 61-65, doi:10.1126/science.aaw8634 (2019).

23 Freier, E., Wolf, S. & Gerwert, K. Proton transfer via a transient linear water-molecule chain in a membrane protein. *Proceedings of the National Academy of Sciences of the United States of America* **108**, 11435-11439, doi:10.1073/pnas.1104735108 (2011).

24 Regan, L. & DeGrado, W. F. Characterization of a helical protein designed from first principles. *Science* **241**, 976-978, doi:10.1126/science.3043666 (1988).

25 Walsh, S. T., Cheng, H., Bryson, J. W., Roder, H. & DeGrado, W. F. Solution structure and dynamics of a de novo designed three-helix bundle protein. *Proceedings of the National Academy of Sciences of the United States of America* **96**, 5486-5491, doi:10.1073/pnas.96.10.5486 (1999).

26 Kuhlman, B. *et al.* Design of a novel globular protein fold with atomic-level accuracy. *Science* **302**, 1364-1368, doi:10.1126/science.1089427 (2003).

27 Vorobieva, A. A. *et al.* De novo design of transmembrane beta barrels. *Science* **371**, doi:10.1126/science.abc8182 (2021).

28 Yang, C. *et al.* Bottom-up de novo design of functional proteins with complex structural features. *Nat Chem Biol* **17**, 492-500, doi:10.1038/s41589-020-00699-x (2021).

29 Polizzi, N. F. & DeGrado, W. F. A defined structural unit enables de novo design of small-molecule-binding proteins. *Science* **369**, 1227-1233, doi:10.1126/science.abb8330 (2020).

30 Cao, L. *et al.* De novo design of picomolar SARS-CoV-2 miniprotein inhibitors. *bioRxiv : the preprint server for biology*, doi:10.1101/2020.08.03.234914 (2020).

31 Fleishman, S. J. *et al.* Computational design of proteins targeting the conserved stem region of influenza hemagglutinin. *Science* **332**, 816-821, doi:10.1126/science.1202617 (2011).

32 Jiang, L. *et al.* De novo computational design of retro-aldol enzymes. *Science* **319**, 1387-1391, doi:10.1126/science.1152692 (2008).

33 Lassila, J. K., Privett, H. K., Allen, B. D. & Mayo, S. L. Combinatorial methods for small-molecule placement in computational enzyme design. *Proceedings of the National Academy of Sciences of the United States of America* **103**, 16710-16715, doi:10.1073/pnas.0607691103 (2006).

34 Polizzi, N. F. *et al.* De novo design of a hyperstable non-natural protein-ligand complex with sub-A accuracy. *Nature chemistry* **9**, 1157-1164, doi:10.1038/nchem.2846 (2017).

35 Leaver-Fay, A. *et al.* ROSETTA3: an object-oriented software suite for the simulation and design of macromolecules. *Methods in enzymology* **487**, 545-574, doi:10.1016/B978-0-12-381270-4.00019-6 (2011).

36 Koga, N. *et al.* Principles for designing ideal protein structures. *Nature* **491**, 222-227, doi:10.1038/nature11600 (2012).

37 Scott, A. J. *et al.* Constructing ion channels from water-soluble alpha-helical barrels. *Nature chemistry* **13**, 643-650, doi:10.1038/s41557-021-00688-0 (2021).

38 Xu, C. *et al.* Computational design of transmembrane pores. *Nature* **585**, 129-134, doi:10.1038/s41586-020-2646-5 (2020).

39 Joh, N. H. *et al.* De novo design of a transmembrane Zn2+-transporting four-helix bundle. *Science* **346**, 1520-1524, doi:10.1126/science.1261172 (2014).

40 Lu, P. *et al.* Accurate computational design of multipass transmembrane proteins. *Science* **359**, 1042-1046, doi:10.1126/science.aaq1739 (2018).

41 Thomaston, J. L. *et al.* X-ray Crystal Structure of the Influenza A M2 Proton Channel S31N Mutant in Two Conformational States: An Open and Shut Case. *J. Am. Chem. Soc.* **141**, 11481-11488, doi:10.1021/jacs.9b02196 (2019).

42 Saotome, K. *et al.* Structures of the otopetrin proton channels Otop1 and Otop3. *Nature structural & molecular biology* **26**, 518-525, doi:10.1038/s41594-019-0235-9 (2019).

43 Mravic, M. *et al.* Packing of apolar side chains enables accurate design of highly stable membrane proteins. *Science* **363**, 1418-1423, doi:10.1126/science.aav7541 (2019).

44 Klesse, G., Rao, S., Sansom, M. S. P. & Tucker, S. J. CHAP: A Versatile Tool for the Structural and Functional Annotation of Ion Channel Pores. *J Mol Biol* **431**, 3353-3365, doi:10.1016/j.jmb.2019.06.003 (2019).

45 Lee, S., Liang, R., Voth, G. A. & Swanson, J. M. Computationally Efficient Multiscale Reactive Molecular Dynamics to Describe Amino Acid Deprotonation in Proteins. *Journal of chemical theory and computation* **12**, 879-891, doi:10.1021/acs.jctc.5b01109 (2016).

46 Knight, C., Lindberg, G. E. & Voth, G. A. Multiscale reactive molecular dynamics. *J Chem Phys* **137**, 22A525, doi:10.1063/1.4743958 (2012).

47 Yamashita, T., Peng, Y., Knight, C. & Voth, G. A. Computationally Efficient Multiconfigurational Reactive Molecular Dynamics. *Journal of chemical theory and computation* **8**, 4863-4875, doi:10.1021/ct3006437 (2012).

48 Moffat, J. C. *et al.* Proton transport through influenza A virus M2 protein reconstituted in vesicles. *Biophysical journal* **94**, 434-445, doi:10.1529/biophysj.107.109082 (2008).

49 Ma, C. *et al.* Identification of the functional core of the influenza A virus A/M2 proton-selective ion channel. *Proceedings of the National Academy of Sciences of the United States of America* **106**, 12283-12288, doi:10.1073/pnas.0905726106 (2009).

50 Leiding, T., Wang, J., Martinsson, J., DeGrado, W. F. & Arskold, S. P. Proton and cation transport activity of the M2 proton channel from influenza A virus. *Proceedings of the National Academy of Sciences of the United States of America* **107**, 15409-15414, doi:10.1073/pnas.1009997107 (2010).

51 Slope, L. N. & Peacock, A. F. De Novo Design of Xeno-Metallo Coiled Coils. *Chemistry, an Asian journal* **11**, 660-666, doi:10.1002/asia.201501173 (2016).

52 Pinter, T. B. J., Koebke, K. J. & Pecoraro, V. L. Catalysis and Electron Transfer in De Novo Designed Helical Scaffolds. *Angewandte Chemie* **59**, 7678-7699, doi:10.1002/anie.201907502 (2020).

53 Khurana, E. *et al.* Molecular dynamics calculations suggest a conduction mechanism for the M2 proton channel from influenza A virus. *Proceedings of the National Academy of Sciences of the United States of America* **106**, 1069-1074, doi:10.1073/pnas.0811720106 (2009).

54 Yi, M., Cross, T. A. & Zhou, H. X. A secondary gate as a mechanism for inhibition of the M2 proton channel by amantadine. *The journal of physical chemistry. B* **112**, 7977-7979, doi:10.1021/jp800171m (2008).

55 Ramsey, I. S. *et al.* An aqueous H+ permeation pathway in the voltage-gated proton channel Hv1. *Nature structural & molecular biology* **17**, 869-875, doi:10.1038/nsmb.1826 (2010).

56 Chamberlin, A. *et al.* Hydrophobic plug functions as a gate in voltage-gated proton channels. *Proceedings of the National Academy of Sciences of the United States of America* **111**, E273-282, doi:10.1073/pnas.1318018111 (2014).

57 Takeshita, K. *et al.* X-ray crystal structure of voltage-gated proton channel. *Nature structural & molecular biology* **21**, 352-357, doi:10.1038/nsmb.2783 (2014).

58 Wikstrom, M., Krab, K. & Sharma, V. Oxygen Activation and Energy Conservation by Cytochrome c Oxidase. *Chemical reviews* **118**, 2469-2490, doi:10.1021/acs.chemrev.7b00664 (2018).

59 Hofacker, I. & Schulten, K. Oxygen and proton pathways in cytochrome c oxidase. *Proteins: Structure, Function, and Genetics* **30**, 100-107, doi:10.1002/(sici)1097-0134(199801)30:1<100::aid-prot9>3.0.co;2-s (1998).

60 Wikström, M., Verkhovsky, M. I. & Hummer, G. Water-gated mechanism of proton translocation by cytochrome c oxidase. *Biochimica et Biophysica Acta (BBA) - Bioenergetics* **1604**, 61-65, doi:10.1016/s0005-2728(03)00041-0 (2003).

61 Tashiro, M. & Stuchebrukhov, A. A. Thermodynamic properties of internal water molecules in the hydrophobic cavity around the catalytic center of cytochrome c oxidase. *The journal of physical chemistry. B* **109**, 1015-1022, doi:10.1021/jp0462456 (2005).

62 Goyal, P., Lu, J., Yang, S., Gunner, M. R. & Cui, Q. Changing hydration level in an internal cavity modulates the proton affinity of a key glutamate in cytochrome c oxidase. *Proceedings of the National Academy of Sciences of the United States of America* **110**, 18886-18891, doi:10.1073/pnas.1313908110 (2013).

63 Liang, R., Swanson, J. M. J., Wikstrom, M. & Voth, G. A. Understanding the essential proton-pumping kinetic gates and decoupling mutations in cytochrome c oxidase. *Proceedings of the National Academy of Sciences of the United States of America* **114**, 5924-5929, doi:10.1073/pnas.1703654114 (2017).

64 Liang, R., Swanson, J. M., Peng, Y., Wikstrom, M. & Voth, G. A. Multiscale simulations reveal key features of the proton-pumping mechanism in cytochrome c oxidase. *Proceedings of the National Academy of Sciences of the United States of America* **113**, 7420-7425, doi:10.1073/pnas.1601982113 (2016).

65 Lynch, C. I., Rao, S. & Sansom, M. S. P. Water in Nanopores and Biological Channels: A Molecular Simulation Perspective. *Chemical reviews*, doi:10.1021/acs.chemrev.9b00830 (2020).

66 Chen, H. *et al.* Charge delocalization in proton channels, I: the aquaporin channels and proton blockage. *Biophysical journal* **92**, 46-60, doi:10.1529/biophysj.106.091934 (2007).

67 Murata, K. *et al.* Structural determinants of water permeation through aquaporin-1. *Nature* **407**, 599-605, doi:10.1038/35036519 (2000).

68 Mondal, D., Kolev, V. & Warshel, A. Combinatorial Approach for Exploring Conformational Space and Activation Barriers in Computer-Aided Enzyme Design. *ACS Catal* **10**, 6002-6012, doi:10.1021/acscatal.0c01206 (2020).

69 Tunuguntla, R. H., Allen, F. I., Kim, K., Belliveau, A. & Noy, A. Ultrafast proton transport in sub-1-nm diameter carbon nanotube porins. *Nature nanotechnology* **11**, 639-644, doi:10.1038/nnano.2016.43 (2016).

70 Geng, J. *et al.* Stochastic transport through carbon nanotubes in lipid bilayers and live cell membranes. *Nature* **514**, 612-615, doi:10.1038/nature13817 (2014).

71 Jiang, T. *et al.* Single-chain heteropolymers transport protons selectively and rapidly. *Nature* **577**, 216-220, doi:10.1038/s41586-019-1881-0 (2020).

72 Caffrey, M. & Cherezov, V. Crystallizing membrane proteins using lipidic mesophases. *Nature protocols* **4**, 706-731, doi:10.1038/nprot.2009.31 (2009).

73 Caffrey, M. Crystallizing membrane proteins for structure determination: use of lipidic mesophases. *Annual review of biophysics* **38**, 29-51, doi:10.1146/annurev.biophys.050708.133655 (2009).

74 Kabsch, W. XDS. *Acta crystallographica. Section D, Biological crystallography* **66**, 125-132, doi:10.1107/S0907444909047337 (2010).

75 Winn, M. D. *et al.* Overview of the CCP4 suite and current developments. *Acta crystallographica. Section D, Biological crystallography* **67**, 235-242, doi:10.1107/S0907444910045749 (2011).

76 McCoy, A. J. *et al.* Phaser crystallographic software. *Journal of applied crystallography* **40**, 658-674, doi:10.1107/S0021889807021206 (2007).

77 Emsley, P., Lohkamp, B., Scott, W. G. & Cowtan, K. Features and development of Coot. *Acta crystallographica. Section D, Biological crystallography* **66**, 486-501, doi:10.1107/S0907444910007493 (2010).

78 Afonine, P. V. *et al.* Towards automated crystallographic structure refinement with phenix.refine. *Acta crystallographica. Section D, Biological crystallography* **68**, 352-367, doi:10.1107/S0907444912001308 (2012).

79 Böckmann, A. *et al.* Characterization of different water pools in solid-state NMR protein samples. *J. Biomol. NMR* **45**, 319-327 (2009).

80 Luo, W. & Hong, M. Conformational changes of an ion channel detected through water-protein interactions using solid-state NMR spectroscopy. *J. Am. Chem. Soc.* **132**, 2378-2384, doi:10.1021/ja9096219 (2010).
81 Williams, J. K. & Hong, M. Probing membrane protein structure using water polarization transfer solid-state NMR. *Journal of magnetic resonance* **247**, 118-127, doi:10.1016/j.jmr.2014.08.007 (2014).
82 Mandala, V. S. *et al.* Structure and drug binding of the SARS-CoV-2 envelope protein transmembrane domain in lipid bilayers. *Nature structural & molecular biology* **27**, 1202-1208, doi:10.1038/s41594-020-00536-8 (2020).
83 Gelenter, M. D. *et al.* Water orientation and dynamics in the closed and open influenza B virus M2 proton channels. *Communications biology* **4**, 338, doi:10.1038/s42003-021-01847-2 (2021).
84 Hong, M. *et al.* Coupling amplification in 2D MAS NMR and its application to torsion angle determination in peptides. *Journal of magnetic resonance* **129**, 85-92, doi:10.1006/jmre.1997.1242 (1997).
85 Krivov, G. G., Shapovalov, M. V. & Dunbrack, R. L., Jr. Improved prediction of protein side-chain conformations with SCWRL4. *Proteins* **77**, 778-795, doi:10.1002/prot.22488 (2009).
86 Lomize, M. A., Pogozheva, I. D., Joo, H., Mosberg, H. I. & Lomize, A. L. OPM database and PPM web server: resources for positioning of proteins in membranes. *Nucleic Acids Res* **40**, D370-376, doi:10.1093/nar/gkr703 (2012).
87 Humphrey, W., Dalke, A. & Schulten, K. VMD-Visual Molecular Dynamics. *J. Molec. Graphics* **14**, 33-38 (1996).
88 Van Der Spoel, D. *et al.* GROMACS: fast, flexible, and free. *Journal of computational chemistry* **26**, 1701-1718, doi:10.1002/jcc.20291 (2005).
89 Huang, J. & MacKerell, A. D., Jr. CHARMM36 all-atom additive protein force field: validation based on comparison to NMR data. *Journal of computational chemistry* **34**, 2135-2145, doi:10.1002/jcc.23354 (2013).
90 Jo, S., Kim, T., Iyer, V. G. & Im, W. CHARMM-GUI: a web-based graphical user interface for CHARMM. *Journal of computational chemistry* **29**, 1859-1865, doi:10.1002/jcc.20945 (2008).
91 Jo, S., Kim, T. & Im, W. Automated builder and database of protein/membrane complexes for molecular dynamics simulations. *PloS one* **2**, e880, doi:10.1371/journal.pone.0000880 (2007).
92 Wu, E. L. *et al.* CHARMM-GUI Membrane Builder toward realistic biological membrane simulations. *Journal of computational chemistry* **35**, 1997-2004, doi:10.1002/jcc.23702 (2014).
93 Lee, J. *et al.* CHARMM-GUI Input Generator for NAMD, GROMACS, AMBER, OpenMM, and CHARMM/OpenMM Simulations Using the CHARMM36 Additive Force Field. *Journal of chemical theory and computation* **12**, 405-413, doi:10.1021/acs.jctc.5b00935 (2016).
94 Best, R. B. *et al.* Optimization of the additive CHARMM all-atom protein force field targeting improved sampling of the backbone phi, psi and side-chain chi(1) and chi(2) dihedral angles. *Journal of chemical theory and computation* **8**, 3257-3273, doi:10.1021/ct300400x (2012).

95 Abraham, M. J. *et al.* GROMACS: High performance molecular simulations through multi-level parallelism from laptops to supercomputers. *SoftwareX* **1-2**, 19-25, doi:10.1016/j.softx.2015.06.001 (2015).

96 Nelson, J. G., Peng, Y., Silverstein, D. W. & Swanson, J. M. Multiscale Reactive Molecular Dynamics for Absolute pKa Predictions and Amino Acid Deprotonation. *Journal of chemical theory and computation* **10**, 2729-2737, doi:10.1021/ct500250f (2014).

97 Biswas, R., Tse, Y. L., Tokmakoff, A. & Voth, G. A. Role of Presolvation and Anharmonicity in Aqueous Phase Hydrated Proton Solvation and Transport. *The journal of physical chemistry. B* **120**, 1793-1804, doi:10.1021/acs.jpcb.5b09466 (2016).

98 Day, T. J. F., Soudackov, A. V., Čuma, M., Schmitt, U. W. & Voth, G. A. A second generation multistate empirical valence bond model for proton transport in aqueous systems. *The Journal of Chemical Physics* **117**, 5839-5849, doi:10.1063/1.1497157 (2002).

99 Plimpton, S. Fast Parallel Algorithms for Short-Range Molecular Dynamics. *Journal of Computational Physics* **117**, 1-19, doi:10.1006/jcph.1995.1039 (1995).

100 Bonomi, M. B., G.; Camilloni, C.; et al. Promoting transparency and reproducibility in enhanced molecular simulations. *Nature methods* **16**, 670-673, doi:10.1038/s41592-019-0506-8 (2019).

101 Tribello, G. A., Bonomi, M., Branduardi, D., Camilloni, C. & Bussi, G. PLUMED 2: New feathers for an old bird. *Computer Physics Communications* **185**, 604-613, doi:10.1016/j.cpc.2013.09.018 (2014).

102 Grossfield, A. "WHAM: the weighted histogram analysis method", version 2.0.9, http://membrane.urmc.rochester.edu/wordpress/?page_id=126.

103 Hunter, J. D. Matplotlib: A 2D graphics environment. *Comput Sci Eng* **9**, 90-95, doi: 10.1109/Mcse.2007.55 (2007).

104 Hodel, A., Kim, S.-H., Brünger, A.T. *Acta Cryst.* A**48**, 851-858. doi: 10.1107/S0108767392006044 (1992).
